# A single cell *Arabidopsis* root atlas reveals developmental trajectories in wild type and cell identity mutants

**DOI:** 10.1101/2020.06.29.178863

**Authors:** Rachel Shahan, Che-Wei Hsu, Trevor M. Nolan, Benjamin J. Cole, Isaiah W. Taylor, Anna Hendrika Cornelia Vlot, Philip N. Benfey, Uwe Ohler

**Affiliations:** Department of Biology, Duke University, Durham, North Carolina 27708, USA; Department of Biology, Humboldt Universität zu Berlin, 10117 Berlin, Germany; The Berlin Institute for Medical Systems Biology, Max Delbrück Center for Molecular Medicine, 10115 Berlin, Germany; Department of Energy Joint Genome Institute, Walnut Creek, California 94598, USA; Technische Universität Berlin, Germany; Howard Hughes Medical Institute, Duke University, Durham, North Carolina 27708, USA; Department of Computer Science, Humboldt Universität zu Berlin, 10117 Berlin, Germany

## Abstract

Cell fate acquisition is a fundamental developmental process in all multicellular organisms. Yet, much is unknown regarding how a cell traverses the pathway from stem cell to terminal differentiation. Advances in single cell genomics^1^ hold promise for unraveling developmental mechanisms^2–3^ in tissues^4^, organs^5–6^, and organisms^7–8^. However, lineage tracing can be challenging for some tissues^9^ and integration of high-quality datasets is often necessary to detect rare cell populations and developmental states^10,11^. Here, we harmonized single cell mRNA sequencing data from over 110,000 cells to construct a comprehensive atlas for a stereotypically developing organ with indeterminate growth, the *Arabidopsis* root. To test the utility of the atlas to interpret new datasets, we profiled mutants for two key transcriptional regulators at single cell resolution, *shortroot* and *scarecrow*. Although both transcription factors are required for early specification of cell identity^12^, our results suggest the existence of an alternative pathway acting in mature cells to specify endodermal identity, for which *SHORTROOT* is required. Uncovering the architecture of this pathway will provide insight into specification and stabilization of the endodermis, a tissue analogous to the mammalian epithelium. Thus, the atlas is a pivotal advance for unraveling the transcriptional programs that specify and maintain cell identity to regulate organ development in space and time.

## Main text

Precisely controlled transcriptional networks specify cell identity, relate positional information, and regulate maturation^12^. Defining how these networks orchestrate organ development and function requires detailed knowledge of spatiotemporal gene expression patterns for each cell type and developmental state. Here, we present the first large-scale *Arabidopsis* root gene expression atlas at single cell resolution. Using a general-purpose data pre-processing pipeline and an iterative, integrative strategy for annotation, we show that the atlas provides enhanced resolution to identify gene expression dynamics underlying the differentiation of each cell type and tissue in wild type and in cell-identity mutants.

The cellular organization of the *Arabidopsis thaliana* root simplifies the study of its spatiotemporal development^13^ (Fig 1a). Cell types are arranged in concentric layers around a central vasculature. Cell lineages are ordered longitudinally along a temporal developmental axis, with the oldest cells closest to the shoot and the youngest cells adjacent to the stem cell niche at the root tip. With each new cell division at the root tip, older cells are displaced shootward from the stem cell niche. Thus, the root enables interrogation of the full trajectory from stem cell to differentiated tissue^14,15^.

**Figure 1.**
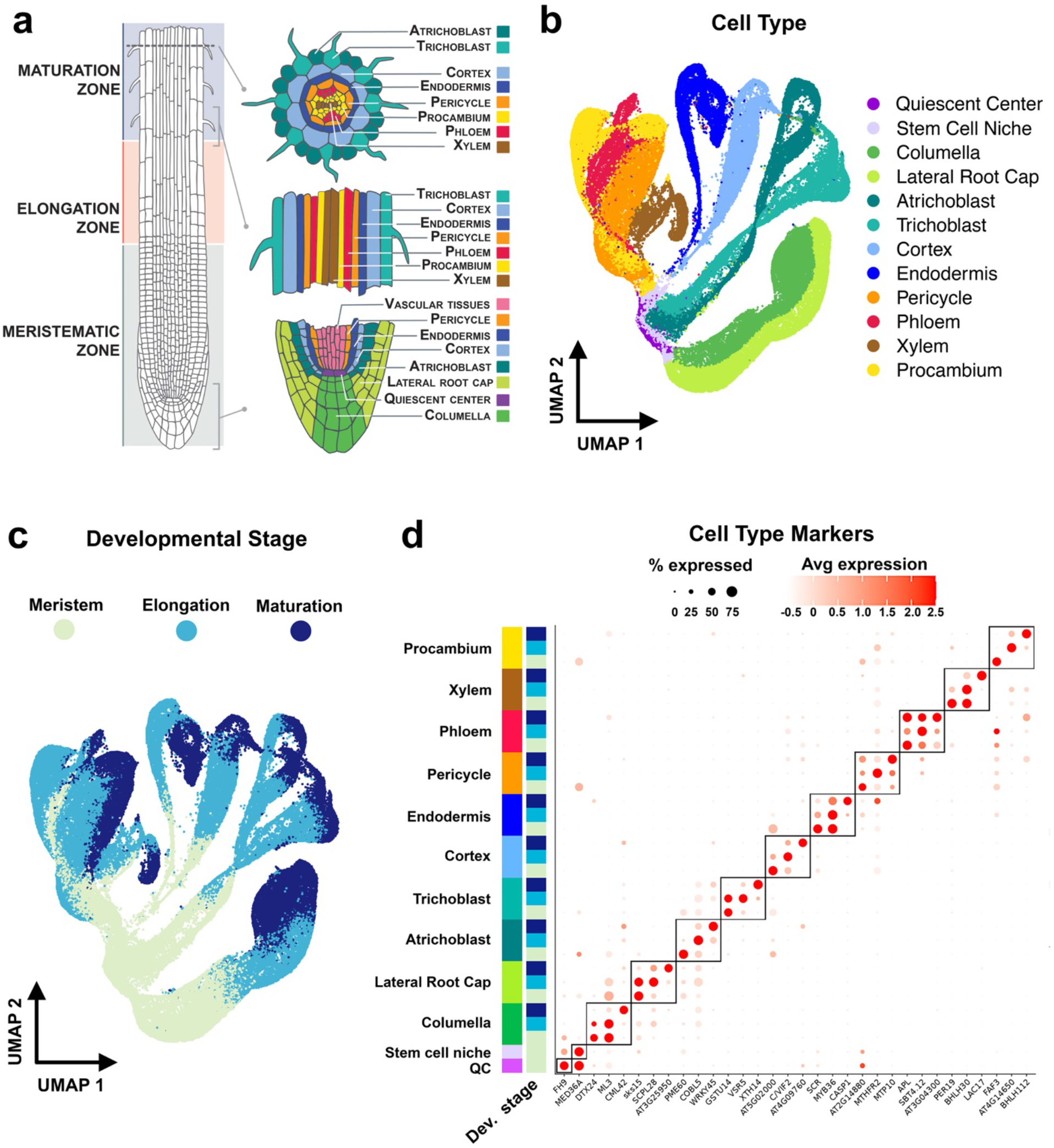
110,000 cell root atlas representing all major cell types. Given the simple structure of the *Arabidopsis* root (**a**; Illustration adapted from the Plant Illustrations repository^45^) the atlas UMAP provides an intuitive visualization with cell types (**b**) and developmental stages (**c**) separated on the X and Y axes, respectively. Although lateral root cap and columella cells are present only at the root tip, meristematic, elongation, and maturation developmental stage labels in (**b**) are consistent with other cell types for simplicity. The crossing over or apparent mixture between some cell types (**b**) is a result of 2D projection and absent in 3D (Supplementary Movie 1). **d**) Expression patterns of marker genes for each developmental stage/cell type combination (identified by Receiver Operating Characteristic (ROC) analysis). Black boxes denote markers from each cell type. Colors of side annotations indicate cell type and developmental stage.

The *Arabidopsis* root is a tractable model organ with established markers for most cell types as well as expression profiles for morphologically defined developmental stages^16–18^. Several groups have used the root to demonstrate the applicability of droplet-based single cell RNA sequencing (scRNA-seq) to plants^19–23^. However, a comprehensive root atlas encompassing all major cell types and developmental states is required to define the spatiotemporal transcriptional dynamics underlying organ development.

## Integration of 110,000 cells produces an organ-scale atlas

To build a harmonized atlas at single cell resolution, we used the 10X Genomics scRNA-seq platform to profile over 96,000 cells from 13 biological replicates of whole, WT roots ranging in age from five to seven days post-germination (Supplementary Dataset 1). Gene expression matrices calculated by kallisto^24^ and bustools^25^ served as input to <u>**C**</u>ell prepr<u>**O**</u>cessing <u>**PI**</u>peline ka<u>**L**</u>list<u>**O**</u> bus<u>**T**</u>ools (COPILOT), our pre-processing software, which incorporates detection and removal of low-quality cells and doublet cells (Supplementary Dataset 2; Methods).

Based on quality assessment by COPILOT, we additionally selected three published root scRNA-seq datasets^19,21^ to augment datasets we generated and to demonstrate the feasibility of integrating *Arabidopsis* data produced by different groups (Supplementary Dataset 1). After excluding mitochondrial and chloroplast genes, as well as genes affected by protoplasting (the process of dissociating plant cells from their cell walls^19^), we used the multi-dataset integration pipeline in Seurat^10,26^ (Methods) to harmonize the cells into an organ-scale atlas (Extended Data Fig. 1; Supplementary Dataset 1).

## Cell annotation places tissues in known developmental contexts

We assigned each cell to one of twelve root cell types (Fig 1b) and to each of the three developmental stages (Fig 1c) by combining information from three approaches for cell type annotation (Supplementary Datasets 1 and 3; Methods). In the first approach, we calculated the correlation coefficient of each cell’s expression profile to published gene expression profiles of root cell types isolated with fluorescent reporters^17,18^. Secondly, we used an information-theoretic approach to compute Index of Cell Identity (ICI) scores for each cell^27,28^ (Extended Data Fig. 2; Supplementary Datasets 4 and 5). The ICI score is quantitative and represents the relative contribution of cell identities as determined from a reference expression profile dataset. Third, we examined the expression of known cell type-specific marker genes in each cell (Supplementary Dataset 1). To assign developmental stage annotation labels, we compared each cell with published bulk gene expression profiles of manually dissected root tissue segments^17,18^.

After annotation, a striking feature of the cell atlas ordination, visible on a Uniform Manifold Approximation and Projection (UMAP) plot, is the presence of four major branches corresponding to four root tissues^13^ (Fig 1b). Lateral root cap (LRC) and columella cells comprise the root cap and form a single branch on the atlas UMAP (Fig 1b). Trichoblast (hair) and atrichoblast (nonhair) cells constitute the epidermis and form a second major branch. Cortex and endodermis cells, which together make up the ground tissue, form a third branch. Finally, the phloem, xylem, procambium, and pericycle cell types are present in the stele tissue and form a fourth branch. The branches originate from a collection of cells within a putative stem cell niche (Fig 1b). Young, dividing meristematic cells are at the base of each branch followed by elongating and finally mature, differentiated cells at the tips (Fig 1c). The branching pattern indicates that distinct cell lineages are transcriptionally distinguishable very early after stem cell division. Overall, the atlas ordination spatially recapitulates what is known about root development and suggests that the combined transcriptome data will be useful in describing relationships between and within individual cell types.

To test the quality of the atlas annotation, we used Seurat to perform differential gene expression analyses and asked if expected markers are enriched in each of the twelve cell type groups produced with our method as compared to the atlas as a whole. We observed enrichment of known cell type markers in their expected cell type groups^12^ (Fig 1d) and also identified many new genes that are enriched in a cell type or developmental stage (Supplementary Dataset 6). We subsequently asked if genes with cell-type specific expression patterns also show localized expression along the developmental gradient. In agreement with previous bulk expression data, we observed that gene expression is rarely specific to both a cell type and developmental stage (Fig 1d).

## Differentiation states and trajectories can be inferred across tissue types

The stereotypic development of the root, coupled with data from existing expression maps, facilitates reconstruction of cell lineages from whole root scRNA-seq data. In addition to improved classification of cells into cell types, the resulting atlas provides enhanced resolution to identify gene expression dynamics underlying the differentiation of each cell and tissue type^14,15^. To infer developmental trajectories, we quantified cell state progression using two tools, CytoTRACE^29^ and scVelo^30^. CytoTRACE predicts the differentiation state of each cell based on the diversity of expressed genes. scVelo is a likelihood-based dynamical model that infers gene-specific rates of transcription, splicing, and degradation for each cell. Using only transcriptional dynamics, scVelo predicts latent time, a representation of the actual time experienced by the cell during differentiation. The inferred latent time has been shown to successfully reconstruct timelines of cellular fate^30^.

When applied to the entire atlas, both packages produce trajectories that are not consistent with previously reported data associating developmental stage with specific gene expression profiles^12,31^ (Extended Data Fig. 3). This result may reflect different maturation rates among tissues/lineages, which could be specific to plant development, or indicate issues with implementing trajectory inference analyses across complex organs. We thus subdivided the atlas into four tissue/lineage groups^13^: stele (consisting of pericycle, procambium, xylem, and, phloem cells), ground tissue (consisting of cortex and endodermis cells), epidermis/lateral root cap (consisting of trichoblast, atrichoblast, and lateral root cap cells), and columella root cap (consisting of columella cells). Unlike the four major branches that are evident on the atlas UMAP ordination (Fig 1b), the four groups isolated here for trajectory inference are based on shared stem cell origin^13^. For example, although LRC and columella cells together comprise the root cap, the columella is patterned by unique stem cells. The LRC, by contrast, is patterned by divisions of the same stem cells that pattern the epidermis. Given the difficulty of distinguishing distinct stem cell types from the atlas (Methods), we incorporated all quiescent center (QC) and stem cell niche (SCN) cells into each group to ensure inclusion of stem cells that pattern each cell type.

As CytoTRACE and scVelo trajectories are strongly correlated (Supplementary Dataset 7), we calculated a ‘consensus’ trajectory for each tissue/lineage group (Extended Data Fig. 4), which represents an averaged developmental progression (Methods). The consensus trajectories agree with developmental stage annotations and reflect existing biological knowledge. For example, in the stele, the procambium and pericycle differentiate more slowly than the xylem and phloem^32,33^.

For closer examination of a predicted trajectory, we focused on the ground tissue (Fig 2), the development of which is well characterized^12^. The direction of the consensus trajectory (Fig 2d) for ground tissue cells within the atlas is consistent with the developmental stage annotation (Fig 2c) and with expected expression profiles of known endodermis and cortex markers, including *SCARECROW* (*SCR*), *MYB36*, and *CASPARIAN STRIP MEMBRANE DOMAIN PROTEIN 1-4* (Fig 2e,f). However, unlike the morphologically determined developmental zones, the consensus trajectory permits examination of gene expression dynamics at a fine resolution. For example, in consensus time group T1 at the beginning of the trajectory, 601 genes are enriched in cortex while 493 genes are enriched in endodermis (Supplementary Dataset 8; Fig 2e). Given that cortex and endodermis are patterned by asymmetric divisions of the same stem cell, genes uniquely expressed early in development for each cell type may include new regulators of cell specification. Together, the inferred trajectories lend credence to the utility of the atlas for downstream analyses of all tissue types, including the identification of gene regulatory networks underlying differentiation.

**Figure 2.**
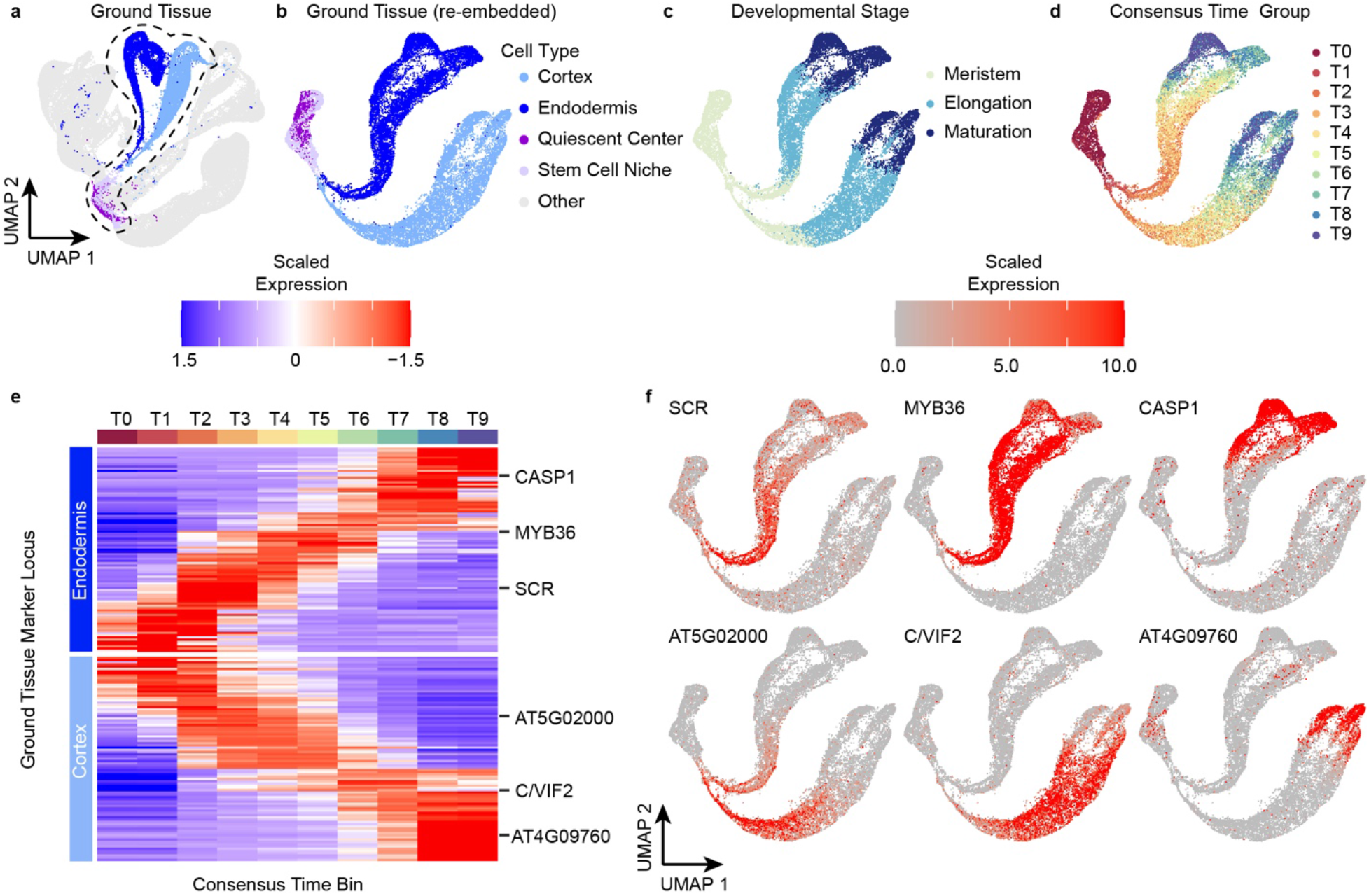
Inferred trajectories reflect the dynamics of cell type and tissue differentiation. Prior to trajectory inference, the atlas was subdivided into four tissues/lineages, one of which is the ground tissue **(a-c)**. The consensus time annotation for the ground tissue **(d)** corresponds with the developmental stage annotation (**c**) and with expression of known endodermis and cortex markers **(e, f)**. Differential expression analyses between ten subgroups (T0 to T9; generated by partitioning the trajectory into ten groups, each with equal numbers of cells) along the ground tissue trajectory identify genes dynamically expressed during cortex and endodermis differentiation **(e).**

## scRNA-seq reveals differentiation pathways of cell identity mutants

In addition to identifying new transcriptional regulators, scRNA-seq allows us to ask how known regulators control tissue and organ development. In the root, the transcription factors *SHORTROOT* (*SHR*) and *SCARECROW* (*SCR*) function in a transcriptional regulatory complex and are essential for stem cell niche maintenance and tissue patterning^34–37^. Using the atlas to inform interpretation of new datasets, we asked how the loss of *SHR* or *SCR* function affects tissue composition as well as cell identity and differentiation. We first transferred cell type and developmental stage annotation labels^10^ from the atlas to two biological replicates each of *shortroot-2* (*shr-2)* and *scarecrow-4* (*scr-4)* mutant roots and to five WT biological replicates profiled alongside the mutants (Supplementary Dataset 1). We performed reference-based integration with Seurat to harmonize these datasets (Methods).

Both *shr-2* and *scr-4* mutants lack the asymmetric cell division that patterns the ground tissue, resulting in a single mutant tissue layer instead of the cortex and endodermis cell layers. Previous detection of tissue-specific markers and morphologies revealed that the mutant layer has cortex-like attributes in *shr-2*^34^
 but a mixture of cortex and endodermis characteristics in *scr-4*^36^. These phenotypes are clearly reflected in the scRNA-seq data given the significant reduction of cells expressing endodermal markers in both *shr-2* and *scr-4* (Fig 3a, c). A second striking observation is the decrease in *shr-2* xylem, phloem, and pericycle cell abundance relative to WT (Fig 3 a,c). Similar changes are also detected for *scr-4* (Fig 3c). These results are consistent with reports of *shr-2* and *scr-4* defects in stele development^38–42^ but were not discernible in earlier *shr-2* scRNA-seq data^19^. Taken together, the resolution of the atlas annotation enables confident detection of major and subtle cell type changes in mutants.

**Figure 3.**
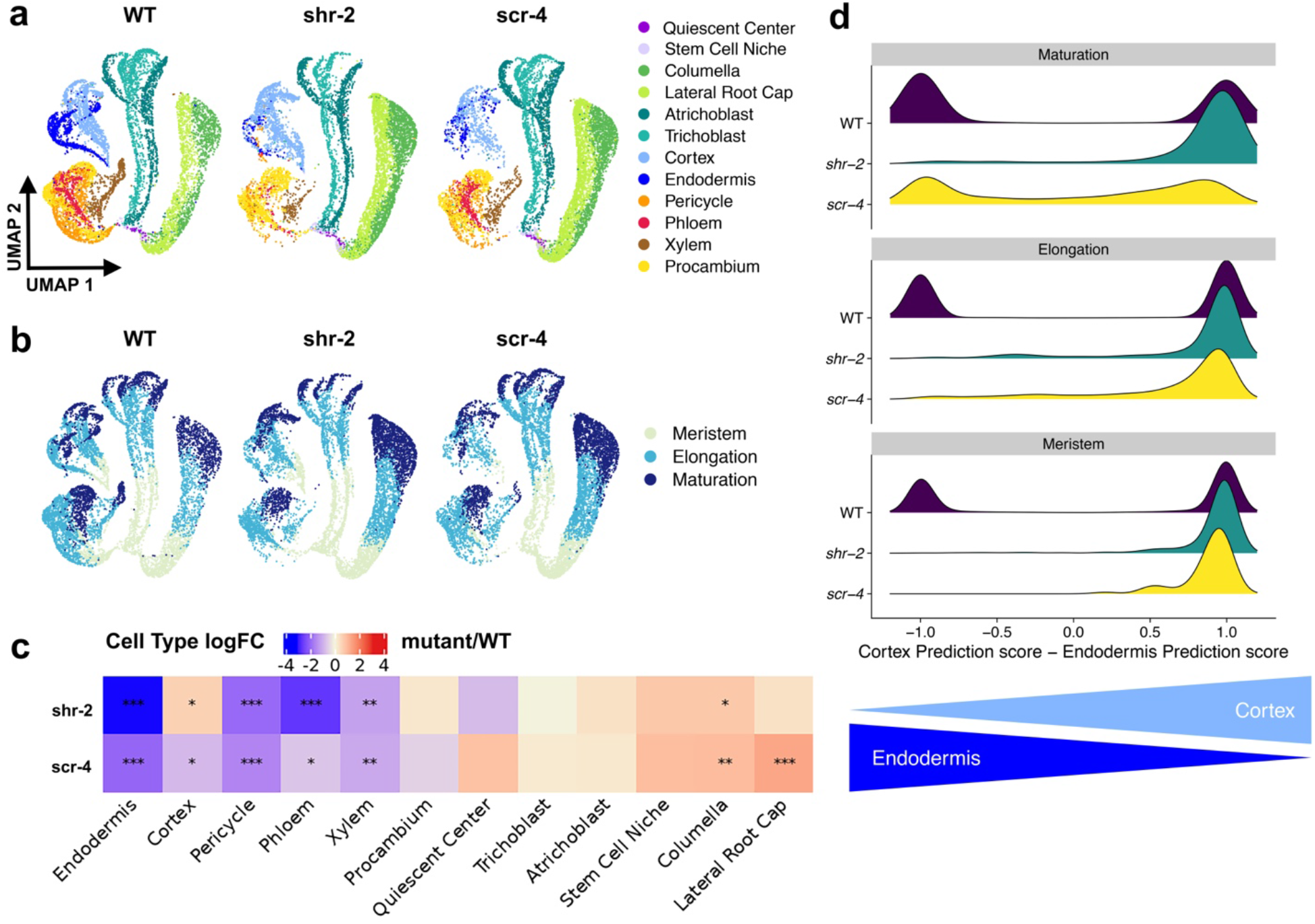
Atlas informs cell type abundance and identity changes in *shr-2* and *scr-4* mutants. **a** and **b)** UMAP projection of WT integrated with *shr-2* and *scr-4* with cell type (a) and developmental stage (b) annotations. Data from each genotype was down-sampled to 10,000 cells to facilitate UMAP comparison. **c)** Differential abundance analysis using full integrated dataset reports significant changes in per-label cell type abundance between mutants and WT. *** False Discovery Rate (FDR) < 0.001; ** FDR < 0.01; * FDR < 0.05. **d)** Difference between cortex and endodermis cell type classification scores for each cell plotted by developmental stage. Classification scores calculated by Seurat during label transfer range from zero (lowest confidence) to one (highest confidence).

We next asked how individual cells contribute to the reported mixed identity of the *scr-4* mutant layer^36^. One hypothesis is that cells acquire an endodermis or cortex identity early in development and the mutant layer is a heterogeneous mixture of the two cell types along the entire cell file. Alternatively, each cell may have a mixture of cortex and endodermis attributes. A third hypothesis is that cells acquire one identity early in development and subsequently change their fate.

To distinguish among these possibilities, we integrated the *scr-4* biological replicates and extracted only cortex and endodermis-annotated cells, which should constitute the mutant layer^36^. We asked: i) how confident is each *scr-4* cell type annotation based on label transfer from the atlas^10^ and ii) if the proportion of cells with each cell type annotation changes according to developmental zone. Using Seurat, the annotation of each *scr-4* cell was assigned using a weighted vote classifier based on reference cell labels from the atlas. This approach gives a quantitative ‘classification score’ for each predicted label^10^. Most meristematic and elongating *scr-4* cells are confidently classified as cortex (Fig 3d). However, differentiating *scr-4* cells are annotated as either cortex or endodermis, though some cells seem to have attributes of both. This result suggests that *scr-4* mutant layer cells are cortex-like in the early stages of development but change their fate to acquire endodermal identity in the maturation zone. A similar analysis for *shr-2* indicates that, unlike *scr-4*, nearly all presumed mutant layer cells are confidently annotated as cortex (Fig 3d).

To explore this possibility that some *scr-4* mutant layer cells acquire cortex identity early in development and subsequently change their fate, we quantified mutant layer developmental progression to infer a trajectory (Fig 4). We first extracted endodermis, cortex, SCN, and QC cells from the two *scr-4* biological replicates. Subsequently, we transferred consensus time labels from the WT ground tissue trajectory (Fig 4g) to the extracted *scr-4* cells (Fig 4 b,e,h). Transferred annotation labels and associated classification scores (Extended Data Fig. 5) indicate that the youngest cells of the *scr-4* mutant layer are confidently cortex-like while endodermis identity is evident only in older cells. By contrast, in a similar trajectory inferred for *shr-2*, cortex identity is predominant in all developmental states after T0 (Fig 4 c,f,i).

**Figure 4.**
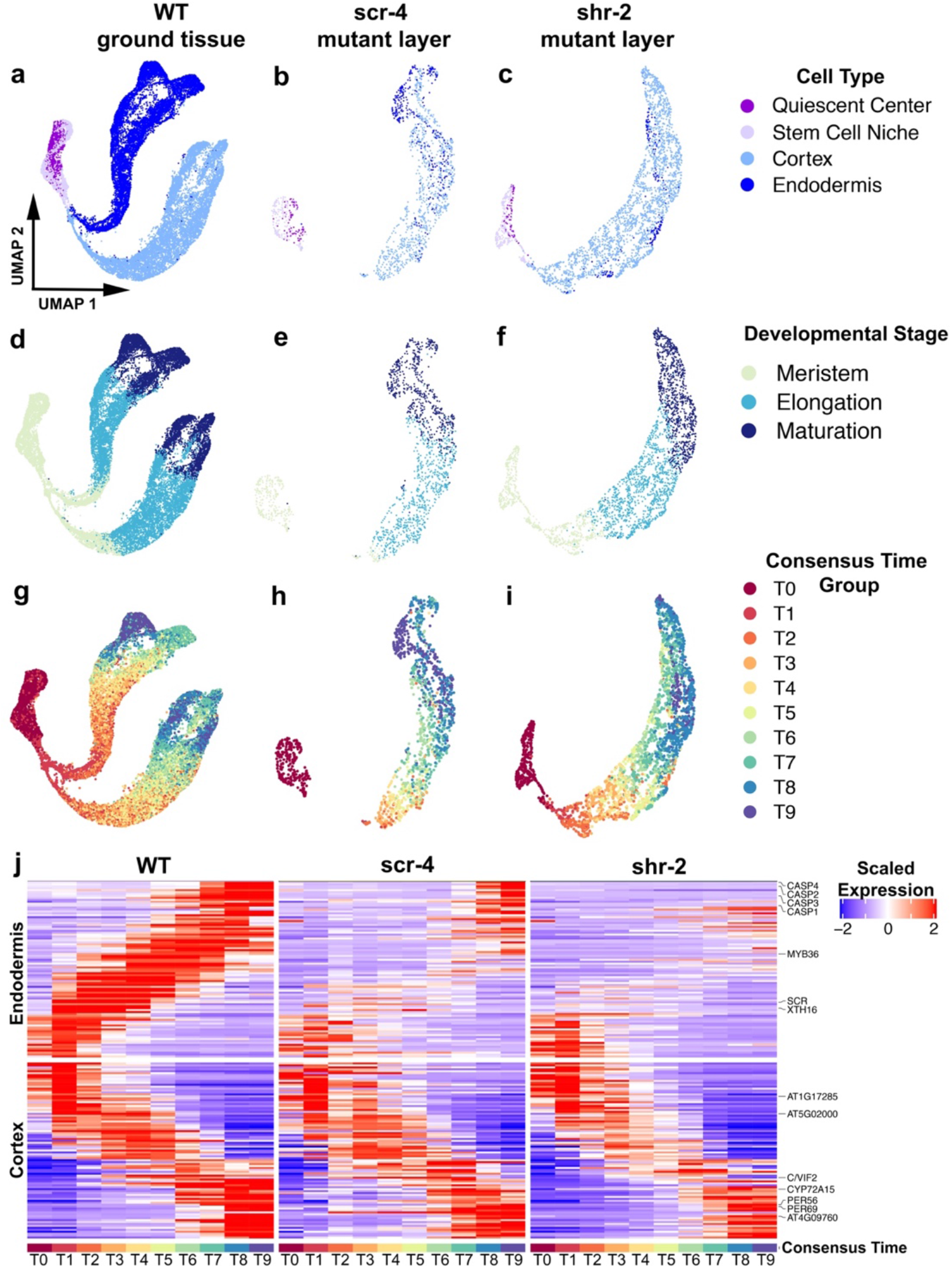
Consensus time trajectory inference suggests trans-differentiation of *scr-4* mutant layer. Cortex, endodermis, QC, and SCN cells were extracted from *scr-4* (**b,e**) and *shr-2* (**c,f**). Consensus time annotation labels were transferred from the WT ground tissue to *scr-4* (**h)** and *shr-2* (**i**). Plots for WT ground tissue are shown for comparison (**a,d,g**). Scaled expression of genes differentially expressed between ten subgroups (T0-T9) along the WT ground tissue trajectory are also shown for *scr-4* and *shr-2* mutant layer trajectories **(j)**.

Given these results, we asked how gene expression dynamics along the *scr-4* and *shr-2* mutant layer trajectories compare to the WT ground tissue trajectory. Genes differentially expressed along the WT cortex trajectory have similar dynamics in *scr-4* and *shr-*2 (Fig 4j). However, only genes expressed at the end of the WT endodermis trajectory, including the *CASPs*, are similarly expressed along the *scr-4* trajectory (group T9; Fig 4j). Conversely, WT endodermis-expressed genes are generally not expressed along the *shr-2* trajectory (Fig 4j). Taken together, our work suggests that *scr-4* mutant layer cells first acquire cortex identity but trans-differentiate in the late stages of development to acquire endodermal identity.

Although the *scr-4* root phenotype was characterized over 20 years ago, we report the first examination of how individual cells at all developmental stages contribute to the mixed identity of the mutant layer. Our results suggest that early specification of endodermis requires both *SHR* and *SCR*. However, in the absence of *SCR*, there exists an alternative pathway able to specify endodermal identity in more mature cells, for which *SHR* is required. Given that root cell lineages are normally transcriptionally distinguishable very early in development (Fig 1b,c), uncovering the architecture of this new pathway could provide insight into novel mechanisms underlying specification and stabilization of the endodermis, an important tissue analogous to the mammalian epithelium^43^.

## Conclusion

The *Arabidopsis* root is a powerful model to investigate the full developmental trajectory from stem cell to differentiated tissue using scRNA-seq. In plants, cell identity is primarily determined by spatial location^44^. The root atlas will facilitate interrogation of how neighboring cell types affect development and what aspects of differentiation are unique or shared between cell types. In addition to mutants, the atlas will guide interpretation of scRNA-seq data from plants treated with hormones or subjected to stress, as well as data from valuable crop species for which comprehensive cell-type markers are unavailable.

## Methods

### Plant material and growth conditions

Seeds from wild type *Arabidopsis thaliana* ecotype Columbia (Col-0), *shortroot-2* (Col-0; ABRC stock number CS2972), and *scarecrow-4* (Landsberg background; ABRC stock number CS6505; we backcrossed to Col-0 > 5 times) were surface sterilized with a 50% (v/v) bleach, 0.05% (v/v) Tween-20 solution for 10 minutes and subsequently stratified for 48 hours at 4°C. Seeds were sown at a density of ~150-300 seeds/row on 1X Linsmaier and Skoog (LSP03-1LT, Caisson Labs; pH 5.7), 1% sucrose media covered by 100 μm nylon mesh. Plates were placed vertically in a Percival chamber programmed to 16h light, 8h dark conditions at 22°C.

### Protoplast Isolation and scRNA-seq

Five days after sowing, 1,000-3,500 primary roots/sample were cut ~0.5 cm from the root tip and placed in a 35 mm-diameter dish containing a 70 μm cell strainer and 4.5 mL enzyme solution (1.25% [w/v] cellulase [ONOZUKA R-10, Yakult], 0.1% Pectolyase [P-3026, Sigma], 0.4 M mannitol, 20 mM MES (pH 5.7), 20 mM KCl, 10 mM CaCl2, 0.1% bovine serum albumin, and 0.000194% (v/v) ß-mercaptoethanol). After digestion at 25°C for 1 hour at 85 rpm on an orbital shaker with occasional stirring, the cell solution was filtered twice through 40 μm cell strainers and centrifuged for 5 minutes at 500 x g in a swinging bucket centrifuge. Subsequently, the pellet was resuspended with 1 mL washing solution (0.4 M mannitol, 20 mM MES (pH 5.7), 20 mM KCl, 10 mM CaCl_2_, 0.1% bovine serum albumin, and 0.000194% (v/v) ß-mercaptoethanol) and centrifuged for 3 minutes at 500 x g. The pellet was resuspended in washing solution to a final concentration of ~1000 cells/ μL. The protoplast suspension was then loaded onto microfluidic chips (10X Genomics) with v3 chemistry to capture either 5,000 or 10,000 cells/ sample. Cells were barcoded with a Chromium Controller (10X Genomics). mRNA was reverse transcribed and Illumina libraries were constructed for sequencing with reagents from a Gene Expression v3 kit (10X Genomics) according to the manufacturer’s instructions. cDNA and final library quality were assessed with a Bioanalyzer High Sensitivity DNA Chip (Agilent). Sequencing was performed with a NovaSeq 6000 (Illumina).

### Read alignment, generation of digital gene expression matrices, and pre-processing

FASTQ files were generated from Illumina BCL files with Cell Ranger (v3.1.0) mkfastq (10X Genomics). Subsequently, gene-by-cell raw count matrices of spliced and un-spliced transcripts were generated using kallisto^24^ (v0.46.2) and bustools^25^ (v0.40.0) as well as R packages BUSpaRse^46^ (v1.1.3) and BSgenome (v1.54.0)^47^. The pipeline is summarized on our scKB GitHub repository (github.com/Hsu-Che-Wei/scKB). Reads were aligned to the Arabidopsis genome BSgenome object (“BSgenome.Athaliana.TAIR.TAIR9”) with TAIR10 gene annotation file. Samples sc_9 and sc_10 (Supplementary Dataset 1) contained a mixture of *Arabidopsis* and rice (*Oryza sativa* X. Kitaake) root protoplasts. Since only the *Arabidopsis* cells were of interest for this study, we mapped the reads to a concatenated version of the *Arabidopsis* TAIR10 and rice MSU7 genomes and retained only the reads which specifically mapped to the *Arabidopsis* genome. The matrices of spliced and un-spliced counts were combined into a total count matrix. Genes with no counts in any cell were removed. Cells were filtered based on the following rationale and procedure. First, putative dying cells were identified based on the enrichment of mitochondrial gene expression (> 5% of the total UMI counts) and the mode of the putative dying cells’ count distribution was treated as the initial boundary to separate cells into two groups representing low and high-quality cells. Second, expression profile references were built for both low and high-quality cells by taking the average of normalized counts. Third, the whole distribution of low-quality cells was recovered by comparing the Pearson correlation coefficient of each high-quality cell to the two references. In other words, if cells in the high-quality group have higher correlation to the low-quality cell profile than the high-quality cell profile, then those cells would be reannotated as low quality. Finally, the low-quality cells and cells enriched in mitochondrial expression were removed along with the top 1% of high-quality cells in order to address any issues associated with outliers. Putative doublets were removed using DoubletFinder^48^ with default parameters according to the estimated doublet rate (10X Genomics Chromium Single Cell 3’ Reagent Kits User Guide (v3 Chemistry)). This pre-processing pipeline is available as an R package, COPILOT (github.com/Hsu-Che-Wei/COPILOT), with a jupyter notebook tutorial. Prior to downstream analyses, protoplasting-induced genes^19^ as well as mitochondrial and chloroplast genes were removed.

### Data normalization and clustering

Using Seurat version 3.1.5, data were normalized using the SCTransform method^49^ followed by principal component analysis (PCA), non-linear dimensionality reduction using UMAP, and clustering. Fifty principal components were calculated using RunPCA function with parameters “approx” set to FALSE. UMAP embedding was generated by RunUMAP function using all 50 principal components with parameters n_neighbors = 30, min_dist = 0.3, umap.method = “umap-learn”, metric = “correlation”. Clustering was done using FindClusters function with parameters res = 0.5, algorithm = 4. Protoplasting-induced genes^19^, mitochondrial genes, and chloroplast genes were excluded during PCA, UMAP dimensionality reduction, and clustering. All steps are incorporated into the COPILOTR package and a jupyter notebook demonstrating the analysis is provi ded (github.com/Hsu-Che-Wei/COPILOT).

### Integration of Seurat objects

Data were integrated following the Seurat reference-based integration pipeline^10,26^. The sample with the highest median UMI/gene per cell and number of genes detected was chosen as the reference (sample name: sc_12; Supplementary Dataset 1). Overall, 16 WT biological replicates were used to build the atlas, including three previously published samples (Supplementary Dataset 1). A jupyter notebook demonstrating the integration process is available on Github (github.com/Hsu-Che-Wei/COPILOT).

### Atlas cell type and developmental stage annotation

The atlas annotation is based on comparison to published whole-transcriptome profiles^17,18^ of root cells isolated from reporter lines as well as known markers (Supplementary Dataset 1) that have been previously validated and show specific local expression on the atlas UMAP. We combined three annotation methods, described below.

### Correlation-based annotation

Prior to scRNA-seq sample integration, Pearson correlation coefficient was calculated between each cell and whole-transcriptome reference expression profiles for cell types and developmental zones. We used bulk RNA-seq data^17^ previously generated for 14 cell types isolated with FACS: phloem and companion cells (cells isolated with a fluorescent reporter for APL), developing cortex (CO2), hair cells (COBL9), mature cortex (CORTEX), mature endodermis (E30), non-hair cells (GL2), columella (PET111), phloem pole pericycle (S17), mature xylem pole (S18), protophloem and metaphloem (S32), developing xylem (S4), endodermis and QC cells (SCR), LRC & non-hair cells (WER), and QC cells (WOX5). We also compared each cell to bulk RNA-seq profiles previously generated for hand-dissected tissue segments corresponding to three morphologically defined developmental zones: meristem, elongation, and maturation^17^. Further, we compared each cell in the atlas to ATH1 microarray data generated for thirteen cell types^18^: QC cells (AGL42), hair cells (COBL9), cortex cells (CORTEX), non-hair cells (GL2), xylem pole pericycle (JO121), lateral root cap (J3411), columella (PET111), phloem pole pericycle (S17), mature xylem (S18), meristematic xylem (S4), phloem (S32), endodermis (SCR5), and phloem (SUC2). Lastly, we compared each cell in the atlas to ATH1 microarray data generated for thirteen hand-dissected tissue segments: columella, meristem-1 (meri-1), meri-2, meri-3, meri-4, meri-5, meri-6, elong-7, elong-8, mat-9, mat-10, mat-11, and mat-12. Each expression profile was built by first aligning the quality-filtered FASTQ reads, which are processed by Trimmomatic^50^ (v.0.39) with default parameters and quality-checked by FastQC^51^ (v.0.11.8), to the TAIR10 genome using STAR^52^ (v.2.7.1a) with default parameters. Then, count normalization was carried out with DESeq and vst function in R package DESeq2^53^ (v1.24.0) with default parameters. 181 genes that are highly variable across cell types in both RNA-seq and microarray data were kept, while 500 highly variable genes across 3 developmental zones and 809 highly variable genes across 13 developmental sections were selected, respectively. The SCTranform-normalized counts in each cell and DEseq2 normalized counts in each expression profile were used to calculate Pearson correlation coefficient. Each cell was labeled with the cell type and developmental zone with which it had the highest correlation coefficient.

### Index of Cell Identity (ICI) calculation

Another method to infer cell identity was an Index of Cell Identity (ICI)-based classification approach^28^. We identified 13 datasets^16–18,54–61^ consisting of cell-type specific gene expression profiles (RNA-seq or ATH1 Microarray) for the 18 cell types considered for this atlas (Extended Data Fig. 2; Supplementary Datasets 4 and 5). RNA-seq data was preprocessed by adapter- and quality-trimming raw FASTAQ reads using the BBDuk tool (BBTools suite^62^; sourceforge.net/projects/bbmap/), using adapter sequences found in the adapters.fa resource within bbtools, and parameters, k=23, mink=11, hdist=1, ktrim=r, and qtrim=10. Trimmed reads were mapped with the STAR^63^ utility (v. 2.7.2b) using default parameters with counts per gene quantified using the quantMode GeneCounts parameter. Read counts were normalized using the DESeq2 R package^53^ (v1.26.0), using a design matrix that treats datasets generated with the same marker:GFP construct as replicates, by running the estimateSizeFactors, estimateDispersions, and the vst functions to normalize and model gene expression. Microarray expression datasets were normalized using the gcrma^64^ R package (v2.58.0). RNA-seq and microarray expression datasets were then normalized together using the FSQN^65^ R package (v0.0.1) to model the RNA-seq gene expression distributions using the microarray data as a reference. Normalized data from both the combined ATH1 and RNASeq datasets, as well as the RNASeq datasets alone, were then used to build two ICI specificity score (spec) tables (using the same methodology as described by Efroni and colleagues^28^, binning expression of each gene into 10 bins, with a minimum background bin set to 3. Markers were identified from this spec table, using a total information level of 50, and normalized, scaled expression of all identified markers was examined in all original datasets. Based on how well correlated each dataset was with its associated datasets of the same cell type, some datasets were filtered out. After dataset filtering, the final spec tables were re-calculated with the same parameters. The spec tables were then used (with an information level of 50) to compute ICI scores, p-values (using the permutation procedure described previously^28^) for all 18 cell types for cells in the atlas, using the log-normalized data values in the SCT slot of each individual dataset’s Seurat object. For each cell, the highest-scoring cell type (from either the combined ATH1/RNASeq or RNASeq only spec tables) was assigned as the ICI-derived annotation.

### Marker-based annotation

The enrichment scores of known cell type-specific markers (Supplementary Dataset 1) were calculated for each cell in the atlas using escoring (github.com/ohlerlab/escoring). Briefly, escoring uses cluster/reference-free, rank based statistics to calculate the significance of local enrichment of gene expression based on the distance between cells. Distance among cells in UMAP space was estimated with radial function over the Euclidean distance (RBF kernel) metric and the size of cell neighborhood was determined by setting the parameter “gamma” to 0.8. A gene is considered significantly enriched with respect to a cell if its enrichment score is more than 5 standard deviations away from the mean of the permutation null distribution. This permutation null distribution is obtained by applying enrichment scoring to 100 times permuted expression vectors. Cells were then annotated with a cell type label according to which significantly enriched marker had the highest enrichment score.

In addition to evaluating known markers, we used escoring to identify new *in silico* markers for regions of the atlas that were annotated with low confidence with the correlation-based and ICI annotation methods. We define a low confidence annotation as correlation coefficient < 0.6 for all known markers and ICI adjusted p-value > 0.01. New markers identified with escoring represent the most enriched genes in the regions/clusters/neighborhood of a specific cell.

### Combination of annotation methods

Final cell type annotations were assigned by combining information from the three annotation approaches. First, considering all confidently annotated cells, if a cell had the same label from at least two annotation methods, then it was annotated as such. Otherwise the cell was treated as not annotated. Labels of some cell types were merged due to lack of consistency among different annotation approaches (e.g. xylem pole pericycle and phloem pole pericycle were merged as pericycle while phloem, companion cells and sieve elements were merged as phloem). Subsequently, we built new reference expression profiles for each cell type by taking the average of the expression values for annotated cells. All cells were then re-annotated using the correlation-based approach by comparison to the newly built references. Finally, at the base of the UMAP, we identified cells from the quiescent center (QC). We labelled cells with support from multiple annotation approaches as “Putative Quiescent Center” in our Seurat objects. For simplicity, we refer to these cells as ‘QC’ here. We also found cells at the base of the UMAP for which a QC annotation was supported by only the escoring marker annotation method. We labelled these cells as “Stem Cell Niche.” In addition, for regions of the UMAP that could not be confidently annotated with correlation-based and ICI methods (e.g., columella and lateral root cap, for which known markers are scarce), the annotation was based solely on the escoring marker annotation method.

We found that putative markers for specific *Arabidopsis* root stem cells^55^ are not specifically expressed on the UMAP and, in general, are highly transcriptionally correlated with the putative QC cells. Thus, we were unable to distinguish specific types of stem cells and chose to use a single annotation label.

To assign a developmental stage annotation to each cell, we used an approach similar to that described for cell type annotation. We built new references for three developmental zones (Meristem, Elongation and Maturation) based on cells that are confidently annotated with the same label according to correlation with RNA-seq^17^ and microarray-based^18^ whole-transcriptome profiles. In addition, we also leveraged information from CytoTRACE (see section “Trajectory inference”) to divide cells into finer groups, which were used to build reference expression profiles. We performed this step separately for each cell type. Finally, all cells were re-annotated using the correlation-based approach by comparing to the newly built references.

In practice, cell type and developmental stage annotations were performed simultaneously, meaning that the newly built references described in previous sections refer to the combination of developmental stage and cell type. A jupyter notebook demonstrating the annotation process is available from Github (github.com/Hsu-Che-Wei/COPILOT).

### Identification of cell-type and developmental stage-specific markers

To identify genes enriched in each radial cell type, we used the FindMarkers function implemented in Seurat. We first prefiltered features using a log2 fold-change threshold of 2 and a minimum percentage difference in expression of 0.25. We then performed differential expression testing using the ROC method that implements a standard AUC classifier. Each cell identity of interest was down-sampled to a maximum of 10,000 cells to speed computation. Classification power (AUC) in this analysis ranges from 0 (random) to 1 (perfect). Only markers with an AUC greater than 0.75 were retained for downstream analysis. We rank ordered markers based on AUC, percentage difference and fold-change. A similar analysis was carried out to identify cell type and developmental stage specific markers. The top 20 markers from each cell type and developmental stage combination were manually examined and one marker for each category was plotted using Seurat’s DotPlot function. Additional plot decorations were constructed using ggplot2.

### Trajectory inference

Trajectories were inferred with R package CytoTRACE^29^ (v0.1.0) and Python-based scVelo^30^ (v.0.1.25), both of which are able to identify the root of the trajectory in an unsupervised manner and are not affected by UMAP embeddings. The batch-corrected (‘integrated’ assay in Seurat object) expression value was manually made non-negative before being fed to CytoTRACE and scVelo. The ratio of spliced and un-spliced transcripts of each gene and cell was calculated using raw counts. The ratio was then multiplied by the batch-corrected non-negative expression count matrix to generate the corresponding spliced and unspliced count matrices, which serve as input for scVelo. Latent time was then estimated by running pp.moments function with parameter, n_pcs = 50, n_neighbors = 100 and tl.velocity function with mode set to “dynamical” in scVelo, while CytoTRACE was implemented with default parameters. A consensus trajectory was derived by taking the average of CytoTRACE and scVelo-inferred latent time. Consensus time was estimated for each tissue/lineage independently to address differences in maturation rates. The consensus time for QC and SCN were then averaged and all the cells in the trajectory were divided into 10 evenly sized groups, each containing the same number of cells. The groups were labeled T0 to T9 based on consensus time order. A jupyter notebook demonstrating how results from the two tools were combined is provided under the GitHub repository for COPILOT (github.com/Hsu-Che-Wei/COPILOT).

### Identification of differentially expressed genes along ground tissue trajectory

We also applied the approach described under ‘Identification of Marker Genes’ to identify genes that vary along the developmental progression of the cortex and endodermis within WT ground tissue. We used the combination of cell type (cortex or endodermis) and consensus time group (10 groups ranging from T0 to T9) as identity of interest among which differential expression analysis was performed. Spearman’s correlation of each marker with consensus time was considered as an additional metric to aid in selecting genes that vary along the gradient of differentiation. Ten genes were selected for each cell type and consensus time group combination resulting in a total of 91 and 92 non-redundant genes to plot for cortex and endodermis, respectively. Genes were arranged according to their highest rank along consensus time. Pseudo-bulk expression profiles within each consensus time group were calculated for each gene and row normalized expression values were then displayed using ComplexHeatmap^66^ in R.

### Mutant analysis

Annotations were transferred from the atlas to the mutant samples as well as to the wild type samples that were grown and processed together with the mutants. Label transfer was performed following Seurat’s label transfer pipeline. A jupyter notebook demonstrating label transfer from the atlas to mutant samples is available on Github (github.com/Hsu-Che-Wei/COPILOT).

### Differential abundance of cell identity labels in *shr-2* and *scr-4* mutants

We used differential abundance analysis to examine which cell types were enriched or depleted in *shr-2* or *scr-4* compared to WT^67^. First, we quantified the number of cells assigned to each label on a per sample basis. We then used the EdgeR^68,69^ package to fit a negative binomial generalized linear model in which the counts represent cells per label. Normalization was conducted according to the number of cells per sample. Separate contrasts were used to compare *shr-2* versus WT or *scr-4* versus WT, each with a blocking factor to account for any potential batch effects between different experimental runs. Differences in abundance were tested using the function glmQLFTest. P-values were adjusted for multiple testing according to Benjamini and Hochberg^70^ and cell type labels with a false discovery rate less than 0.05 were considered significantly altered. We then used ComplexHeatmap^66^ in R to plot the log2 fold-change estimates (mutant/WT) from EdgeR.

## Data Availability

scRNA-seq data and Seurat objects from this study have been deposited in the National Center for Biotechnology Information (NCBI) Gene Expression Omnibus (GEO) with the accession number GSE152766. A web browser for the atlas is soon to be made available on Phytozome.

## Code Availability

All code from this study, as well as detailed tutorials for COPILOT, annotation, and trajectory inference, are available on Github (github.com/Hsu-Che-Wei/COPILOT).

## Acknowledgements

This work was funded by the Howard Hughes Medical Institute to PNB as an Investigator; the US National Institutes of Health (MIRA 1R35GM131725 and NRSA postdoctoral fellowship 1F32GM136030-01) to PNB and RS, respectively; the US National Science Foundation (Postdoctoral Research Fellowships in Biology Program Grant No. IOS-2010686) to TMN; and Deutsche Forschungsgemeinschaft (International Research Training Group 2403) to C-WH and UO. Work was performed by BJC at the U.S. Department of Energy Joint Genome Institute, a DOE Office of Science User Facility, supported under Contract No. DE-AC02-05CH11231. The authors thank Drs. Kook Hui Ryu and John Schiefelbein for advice on protoplast preparation, Dr. Nicolas Devos and Duke GCB for sequencing services, Megan Perkins Jacobs for experimental assistance, Sarah Van Dierdonck for computing assistance, José Maria Muino Acuna for scRNA-seq analysis support, Abdull J. Massri and Drs. Pablo Szekely, Cara Winter, Jazz Dickinson, David McClay, Zhongchi Liu, Jan Philipp Junker, and Gregory Wray for critical reading of the manuscript.

## Author Contributions

RS, C-WH, TMN, BJC, PNB, and UO conceptualized the experiments. RS, TMN, and IWT generated the scRNA-seq data. C-WH, TMN, BJC, AHCV, and RS analyzed the data. RS wrote the manuscript with input from all authors. PNB and UO supervised the experiments and analyses.

## Additional Information

Supplementary Information is available for this paper.

**Extended Data Figure 1.**
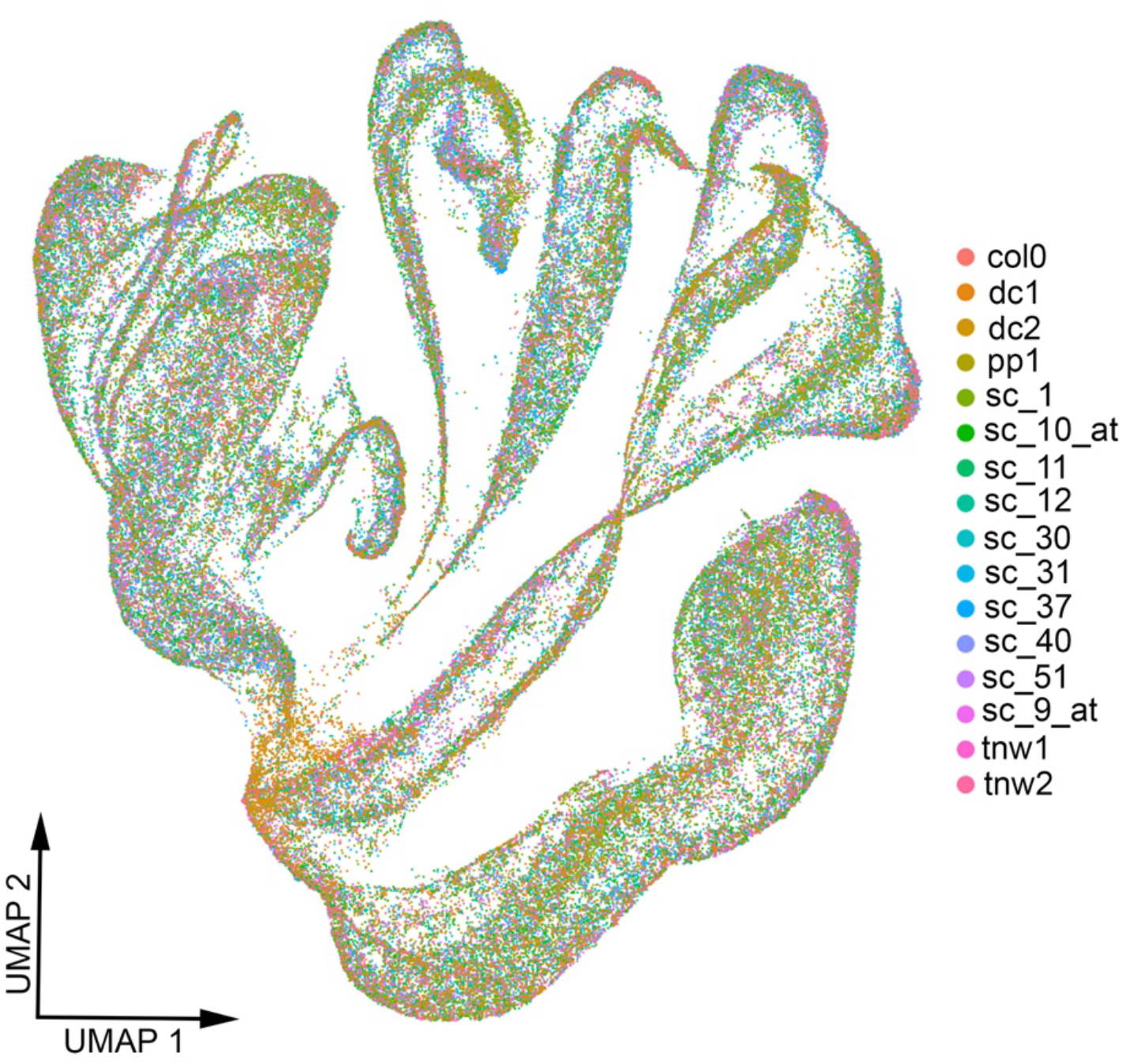
Cells from sixteen WT biological replicates are well mixed in integrated atlas. Each cell on the UMAP is colored based on the sample of origin.

**Extended Data Figure 2.**
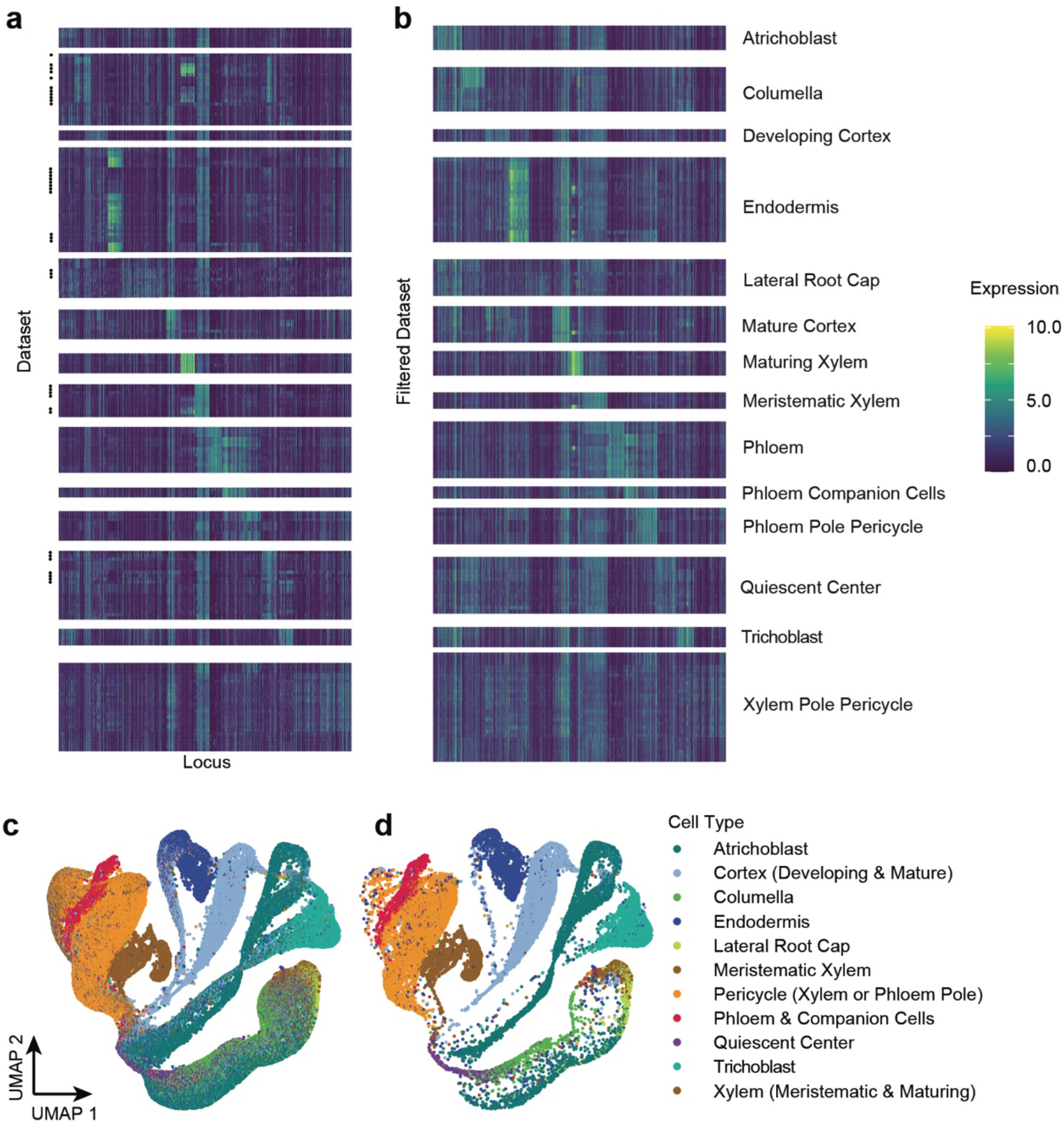
Filtering datasets for ICI computation. Datasets corresponding to FACS-sorted root protoplasts using a variety of cell-type specific GFP markers (278) were downloaded, normalized together (see Methods), and used to make an ICI specification score table^28^. Top markers were identified from this specification score table (corresponding to an information level of 50). **a)** Expression levels of identified markers for each cell-type specific dataset. Dots on the left indicate whether that dataset was subsequently filtered out. **b)** Expression of newly identified markers after filtering, then re-computing the specification table (using both affymetrix-based and RNASeq-based datasets together). A pair of specification tables were then generated using either RNASeq-derived data alone, or both RNASeq- and Affymetrix-derived datasets together. ICI scores were then computed using both methods, and the top-scoring cell type is indicated in **c,** plotted on the Root Cell Atlas UMAP. **d)** The same ICI-based cell identities as shown in **c**, but with non-significant (adjusted P < 0.05) cell type assignments removed.

**Extended Data Figure 3.**
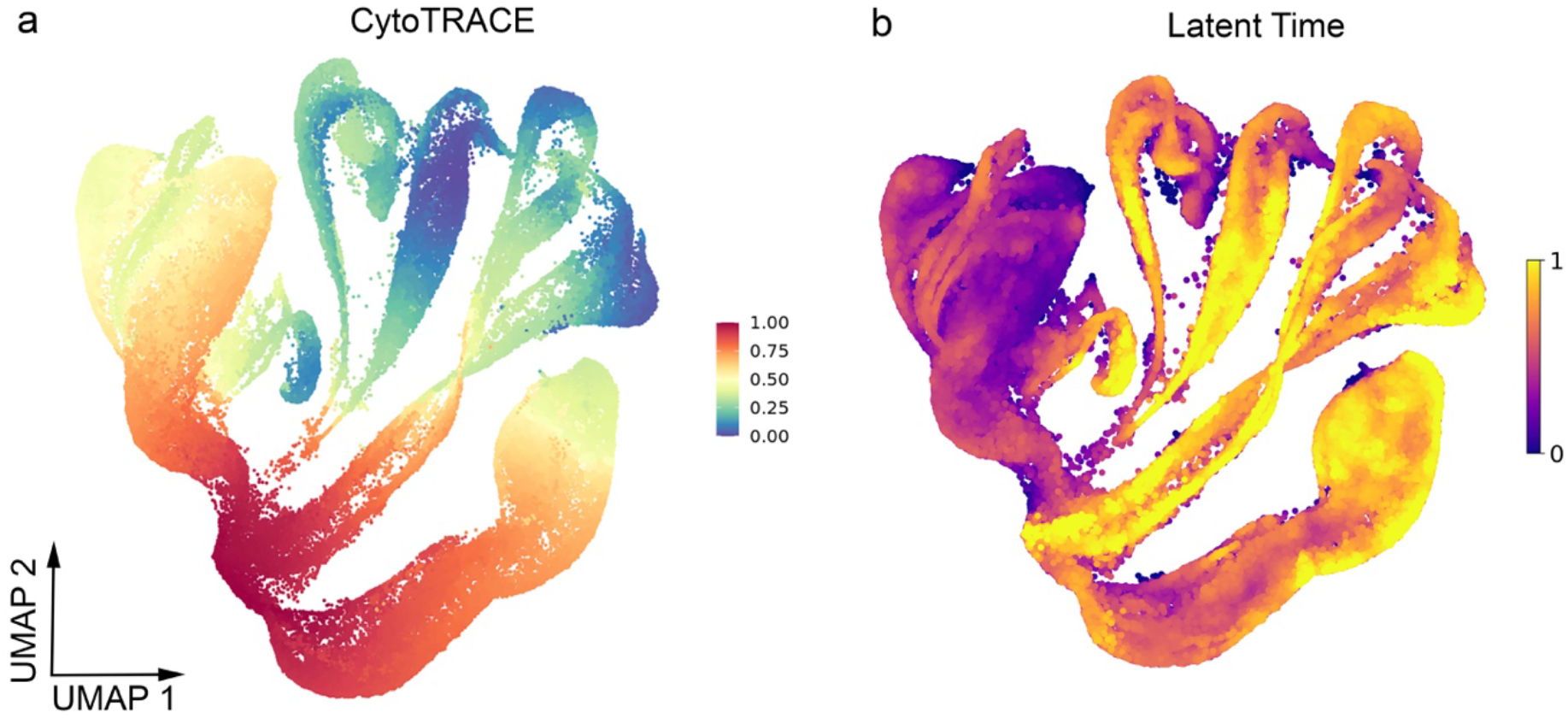
Trajectories inferred for the full atlas do not reflect existing biological knowledge. **a)** CytoTRACE was used to infer the developmental state of each cell in the atlas. Warmer colors denote younger cells while cooler colors denote older cells. **b)** scVelo was used to calculate latent time. Cooler colors denote younger cells while warmer colors denote older cells.

**Extended Data Figure 4.**
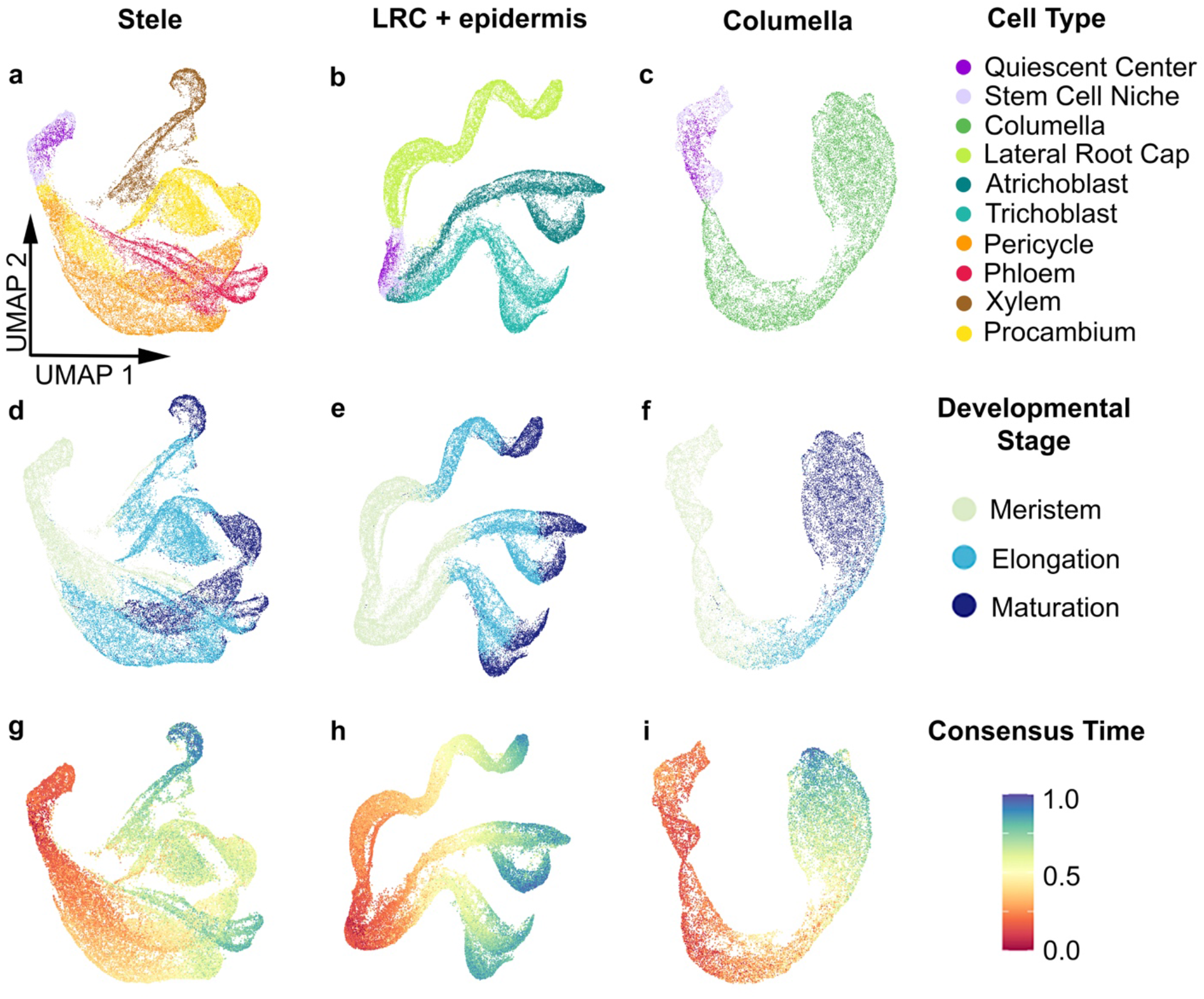
Consensus trajectories inferred for the atlas subdivided into tissues/lineages. The direction of trajectories inferred for **a)** stele (phloem, xylem, procambium, and pericycle), **b)** epidermis (atrichoblast and trichoblast) + lateral root cap, and **c)** columella root cap agree with developmental stage annotations (**d-i**). Warm colors on consensus time plots indicate earlier developmental states while cool colors indicate late developmental states (**g-i**).

**Extended Data Figure 5.**
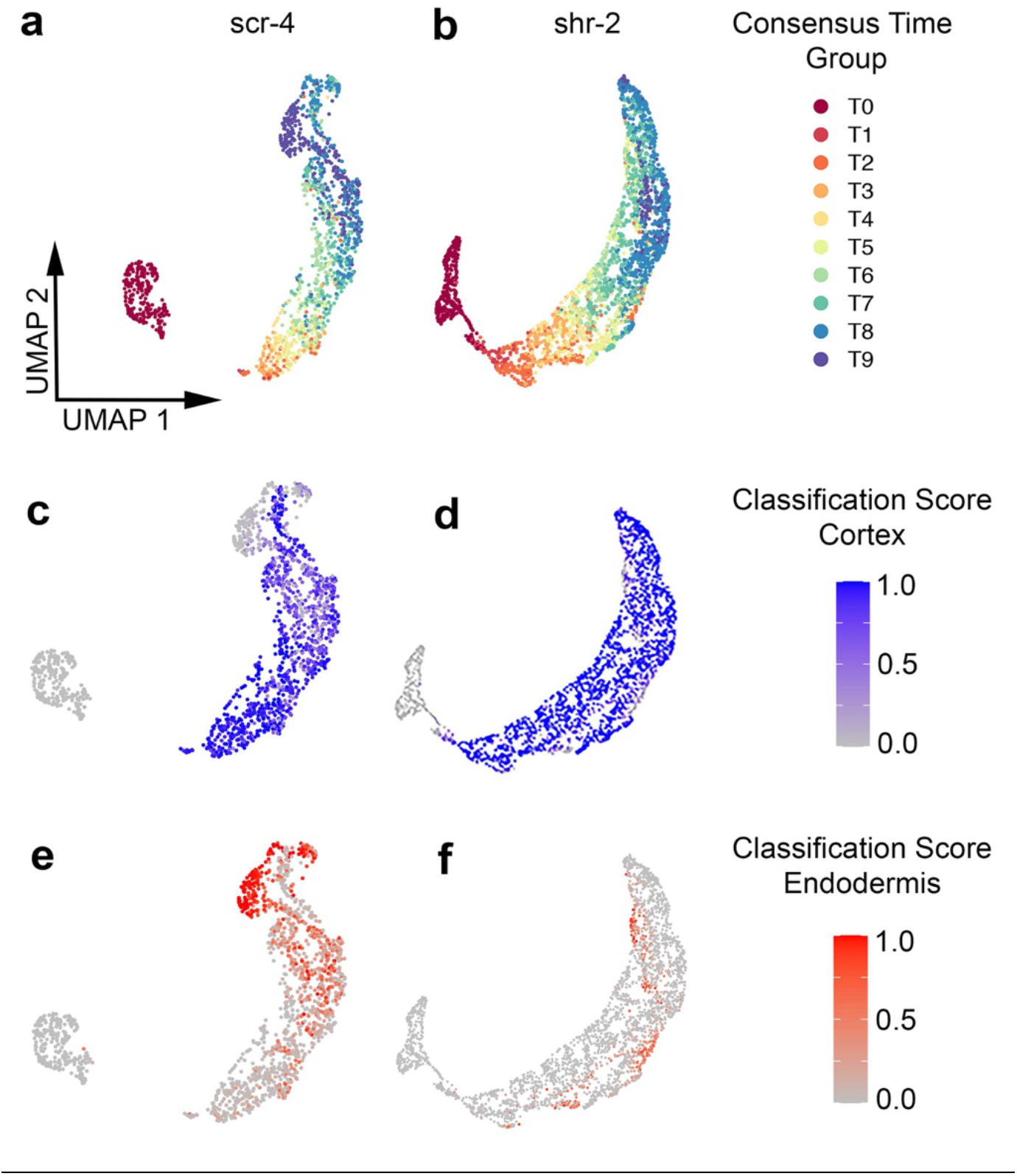
Late stage *scr-4* mutant layer cells are most confidently assigned endodermis identity. The *scr-4* (**a**) and *shr-2* (**b**) mutant layer trajectories are shown with consensus time group labels. Cortex (**c,d**) and endodermis (**e,f**) label classification scores are shown for the two mutant layer trajectories. Classification scores were calculated by Seurat for each cell type label transferred from the atlas. Classification scores range from zero (lowest confidence) to one (highest confidence).

## SI Guide

**Supplementary Dataset 1:**

Excel file with sample information for each scRNA-seq dataset in the atlas (sheet 1); number of cells per cell type and developmental stage label (sheet 2); and curated list of cell type-specific marker genes used for annotation (sheet 3).

**Supplementary Dataset 2:**

PDF with COPILOT summaries for the 16 scRNA-seq datasets in the root atlas.

**Supplementary Dataset 3:**

Excel file with metadata for each cell within the atlas, including final developmental stage, cell type, and consensus time group labels.

**Supplementary Dataset 4:**

Excel file with RNA-seq counts used to build Index of Cell Identity (ICI) specification table.

**Supplementary Dataset 5:**

Excel file with metadata for ICI-based classification method used to infer cell type identity for atlas cells.

**Supplementary Dataset 6:**

Excel file with cell type-specific and developmental stage-specific markers identified from the atlas.

**Supplementary Dataset 7:**

Excel file with Pearson correlation coefficient values calculated for CytoTRACE, scVelo latent time, and consensus time trajectories for each tissue/lineage.

**Supplementary Dataset 8:**

Excel file with genes enriched in cortex and endodermis cells of each consensus time group along the WT ground tissue trajectory.

**Supplementary Movie 1:**

Animation showing the 3D atlas UMAP with cell type annotation.

## References

1. Mereu, E. et al. Benchmarking single-cell RNA-sequencing protocols for cell atlas projects. Nat. Biotechnol. 38, 747–755 (2020).

2. Karaiskos, N. et al. The Drosophila embryo at single-cell transcriptome resolution. Science 358, 194–199 (2017).

3. Farrell, J. A. et al. Single-cell reconstruction of developmental trajectories during zebrafish embryogenesis. Science 360, (2018).

4. Consortium, T. T. M. et al. A Single Cell Transcriptomic Atlas Characterizes Aging Tissues in the Mouse. Preprint at https://www.biorxiv.org/content/10.1101/661728v3 (2020).

5. Asp, M. et al. A Spatiotemporal Organ-Wide Gene Expression and Cell Atlas of the Developing Human Heart. Cell 179, 1647–1660.e19 (2019).

6. Travaglini, K. J. et al. A molecular cell atlas of the human lung from single cell RNA sequencing. Preprint at https://www.biorxiv.org/content/10.1101/742320v2 (2020).

7. Cao, J. et al. Comprehensive single-cell transcriptional profiling of a multicellular organism. Science 357, 661–667 (2017).

8. Han, X. et al. Construction of a human cell landscape at single-cell level. Nature 581 303–309 (2020).

9. Wagner, D. E. & Klein, A. M. Lineage tracing meets single-cell omics: opportunities and challenges. Nat. Rev. Genet. 21, 410–427 (2020).

10. Stuart, T. et al. Comprehensive Integration of Single-Cell Data. Cell 177, 1888–1902.e21 (2019).

11. Luecken, M. D. et al. Benchmarking atlas-level data integration in single-cell genomics. Preprint at https://www.biorxiv.org/content/10.1101/2020.05.22.111161v2 (2020).

12. Drapek, C., Sparks, E. E. & Benfey, P. N. Uncovering Gene Regulatory Networks Controlling Plant Cell Differentiation. Trends Genet. 33, 529–539 (2017).

13. Dolan, L. et al. Cellular organisation of the Arabidopsis thaliana root. Dev. Camb. Engl. 119, 71–84 (1993).

14. Efroni, I. & Birnbaum, K. D. The potential of single-cell profiling in plants. Genome Biol. 17, 65 (2016).

15. McFaline-Figueroa, J. L., Trapnell, C. & Cuperus, J. T. The promise of single-cell genomics in plants. Curr. Opin. Plant Biol. 54, 114–121 (2020).

16. Birnbaum, K. et al. A gene expression map of the Arabidopsis root. Science 302, 1956–1960 (2003).

17. Li, S., Yamada, M., Han, X., Ohler, U. & Benfey, P. N. High-Resolution Expression Map of the Arabidopsis Root Reveals Alternative Splicing and lincRNA Regulation. Dev. Cell 39, 508–522 (2016).

18. Brady, S. M. et al. A high-resolution root spatiotemporal map reveals dominant expression patterns. Science 318, 801–806 (2007).

19. Denyer, T. et al. Spatiotemporal Developmental Trajectories in the Arabidopsis Root Revealed Using High-Throughput Single-Cell RNA Sequencing. Dev. Cell 48, 840–852.e5 (2019).

20. Jean-Baptiste, K. et al. Dynamics of Gene Expression in Single Root Cells of Arabidopsis thaliana. Plant Cell 31, 993–1011 (2019).

21. Ryu, K. H., Huang, L., Kang, H. M. & Schiefelbein, J. Single-Cell RNA Sequencing Resolves Molecular Relationships Among Individual Plant Cells. Plant Physiol. 179, 1444–1456 (2019).

22. Shulse, C. N. et al. High-Throughput Single-Cell Transcriptome Profiling of Plant Cell Types. Cell Rep. 27, 2241–2247.e4 (2019).

23. Zhang, T.-Q., Xu, Z.-G., Shang, G.-D. & Wang, J.-W. A Single-Cell RNA Sequencing Profiles the Developmental Landscape of Arabidopsis Root. Mol. Plant 12, 648–660 (2019).

24. Bray, N. L., Pimentel, H., Melsted, P. & Pachter, L. Near-optimal probabilistic RNA-seq quantification. Nat. Biotechnol. 34, 525–527 (2016).

25. Melsted, P., Ntranos, V. & Pachter, L. The barcode, UMI, set format and BUStools. Bioinformatics 35, 4472–4473 (2019).

26. Butler, A., Hoffman, P., Smibert, P., Papalexi, E. & Satija, R. Integrating single-cell transcriptomic data across different conditions, technologies, and species. Nat. Biotechnol. 36, 411–420 (2018).

27. Birnbaum, K. D. & Kussell, E. Measuring cell identity in noisy biological systems. Nucleic Acids Res. 39, 9093–9107 (2011).

28. Efroni, I., Ip, P.-L., Nawy, T., Mello, A. & Birnbaum, K. D. Quantification of cell identity from single-cell gene expression profiles. Genome Biol. 16, 9 (2015).

29. Gulati, G. S. et al. Single-cell transcriptional diversity is a hallmark of developmental potential. Science 367, 405–411 (2020).

30. Bergen, V., Lange, M., Peidli, S., Wolf, F. A. & Theis, F. J. Generalizing RNA velocity to transient cell states through dynamical modeling. Preprint at https://www.biorxiv.org/content/10.1101/820936v1 (2019).

31. Schiefelbein, J., Zheng, X. & Huang, L. Regulation of epidermal cell fate in Arabidopsis roots: the importance of multiple feedback loops. Front. Plant Sci. 5, (2014).

32. Jouannet, V., Brackmann, K. & Greb, T. (Pro)cambium formation and proliferation: two sides of the same coin? Curr. Opin. Plant Biol. 0, 54–60 (2015).

33. Beeckman, T. & De Smet, I. Pericycle. Curr. Biol. CB 24, R378–379 (2014).

34. Benfey, P. N. et al. Root development in Arabidopsis: four mutants with dramatically altered root morphogenesis. Dev. Camb. Engl. 119, 57–70 (1993).

35. Scheres, B. et al. Mutations affecting the radial organisation of the Arabidopsis root display specific defects throughout the embryonic axis. Development 121, 53–62 (1995).

36. Di Laurenzio, L. et al. The SCARECROW gene regulates an asymmetric cell division that is essential for generating the radial organization of the Arabidopsis root. Cell 86, 423–433 (1996).

37. Helariutta, Y. et al. The SHORT-ROOT gene controls radial patterning of the Arabidopsis root through radial signaling. Cell 101, 555–567 (2000).

38. Levesque, M. P. et al. Whole-genome analysis of the SHORT-ROOT developmental pathway in Arabidopsis. PLoS Biol. 4, e143 (2006).

39. Carlsbecker, A. et al. Cell signalling by microRNA165/6 directs gene dose-dependent root cell fate. Nature 465, 316–321 (2010).

40. Yu, N.-I. et al. Characterization of SHORT-ROOT function in the Arabidopsis root vascular system. Mol. Cells 30, 113–119 (2010).

41. Cui, H. et al. Genome-Wide Direct Target Analysis Reveals a Role for SHORT-ROOT in Root Vascular Patterning through Cytokinin Homeostasis1[W][OA]. Plant Physiol. 157, 1221–1231 (2011).

42. Kim, H. et al. SHORTROOT-Mediated Intercellular Signals Coordinate Phloem Development in Arabidopsis Roots. Plant Cell 32, 1519–1535 (2020).

43. Geldner, N. The Endodermis. Annu. Rev. Plant Biol. 64, 531–558 (2013).

44. Pierre-Jerome, E., Drapek, C. & Benfey, P. N. Regulation of Division and Differentiation of Plant Stem Cells. Annu. Rev. Cell Dev. Biol. 34, 289–310 (2018).

45. Bouché, F. Arabidopsis - Root cell types. (2017) doi:10.6084/m9.figshare.4688752.v1.

## Methods References

46. Moses L & Pachter L. BUSpaRse: kallisto | bustools R utilities. R package version 1.2.2, https://github.com/BUStools/BUSpaRse (2020).

47. Pagès, H. BSgenome: Software infrastructure for efficient representation of full genomes and their SNPs. (Bioconductor version: Release (3.11)). doi:10.18129/B9.bioc.BSgenome (2020).

48. McGinnis, C. S., Murrow, L. M. & Gartner, Z. J. DoubletFinder: Doublet Detection in Single-Cell RNA Sequencing Data Using Artificial Nearest Neighbors. Cell Syst. 8, 329–337.e4 (2019).

49. Hafemeister, C. & Satija, R. Normalization and variance stabilization of single-cell RNA-seq data using regularized negative binomial regression. Genome Biol. 20, 296 (2019).

50. Bolger, A. M., Lohse, M. & Usadel, B. Trimmomatic: a flexible trimmer for Illumina sequence data. Bioinforma. Oxf. Engl. 30, 2114–2120 (2014).

51. Andrews S. FastQC: A quality control tool for high throughput sequence data. https://www.bioinformatics.babraham.ac.uk/projects/fastqc/ (2010).

52. Dobin, A. & Gingeras, T. R. Optimizing RNA-Seq Mapping with STAR. Methods Mol. Biol. Clifton NJ 1415, 245–262 (2016).

53. Love, M. I., Huber, W. & Anders, S. Moderated estimation of fold change and dispersion for RNA-seq data with DESeq2. Genome Biol. 15, 550 (2014).

54. Nawy, T. et al. Transcriptional Profile of the Arabidopsis Root Quiescent Center. Plant Cell 17, 1908–1925 (2005).

55. Clark, N. M. et al. Stem-cell-ubiquitous genes spatiotemporally coordinate division through regulation of stem-cell-specific gene networks. Nat. Commun. 10, 5574 (2019).

56. Lee, J.-Y. et al. Transcriptional and posttranscriptional regulation of transcription factor expression in Arabidopsis roots. Proc. Natl. Acad. Sci. 103, 6055–6060 (2006).

57. Dinneny, J. R. et al. Cell Identity Mediates the Response of Arabidopsis Roots to Abiotic Stress. Science 320, 942–945 (2008).

58. Gifford, M. L., Dean, A., Gutierrez, R. A., Coruzzi, G. M. & Birnbaum, K. D. Cell-specific nitrogen responses mediate developmental plasticity. Proc. Natl. Acad. Sci. 105, 803–808 (2008).

59. Bargmann, B. O. R. et al. A map of cell type-specific auxin responses. Mol. Syst. Biol. 9, 688 (2013).

60. Yadav, R. K., Tavakkoli, M., Xie, M., Girke, T. & Reddy, G. V. A high-resolution gene expression map of the Arabidopsis shoot meristem stem cell niche. Development 141, 27352744 (2014).

61. Birnbaum, K & Yuan, S. Auxin induced endodermal to QC transdifferentiation time series and downstream of JKD analysis. GEO Accession viewer. https://www.ncbi.nlm.nih.gov/geo/query/acc.cgi?acc=GSE61408. (2015).

62. BBMap download | SourceForge.net. https://sourceforge.net/projects/bbmap/.

63. Dobin, A. et al. STAR: ultrafast universal RNA-seq aligner. Bioinformatics 29, 15–21 (2013).

64. Gentry, J. W., Irizarry, R., MacDonald, J. gcrma. (Bioconductor). doi:10.18129/B9.BIOC.GCRMA (2017).

65. Franks, J. M., Cai, G. & Whitfield, M. L. Feature specific quantile normalization enables crossplatform classification of molecular subtypes using gene expression data. Bioinformatics 34, 1868–1874 (2018).

66. Gu, Z., Eils, R. & Schlesner, M. Complex heatmaps reveal patterns and correlations in multidimensional genomic data. Bioinforma. Oxf. Engl. 32, 2847–2849 (2016).

67. Amezquita, R. A. et al. Orchestrating single-cell analysis with Bioconductor. Nat. Methods 17, 137–145 (2020).

68. Robinson, M. D., McCarthy, D. J. & Smyth, G. K. edgeR: a Bioconductor package for differential expression analysis of digital gene expression data. Bioinformatics 26, 139–140 (2010).

69. McCarthy, D. J., Chen, Y. & Smyth, G. K. Differential expression analysis of multifactor RNA-Seq experiments with respect to biological variation. Nucleic Acids Res. 40, 4288–4297 (2012).

70. Benjamini, Y. & Hochberg, Y. Controlling the False Discovery Rate: A Practical and Powerful Approach to Multiple Testing. J. R. Stat. Soc. Ser. B Methodol. 57, 289–300 (1995).

