## Supplementary Dataset 2 for "A single cell *Arabidopsis* root atlas reveals developmental trajectories in wild type and cell identity mutants"

### col0 Summary

Processed by COPILOT

Summary Analysis

#### Parameters

| | |
|---|---|
| Iteration of Filtering | 1 |
| Mitochondrial Expression Threshold | 5 % |
| Top High Quality Cell Filtered | 1 % |
| Doublet Removed | Yes |

#### Cell Stats

| | |
|---|---|
| Estimated Number of High Quality Cell | 6,779 |
| High Quality Cell | 9.28 % |
| Total UMI Counts in High Quality Cell | 136,642,155 |
| UMI Counts in High Quality Cell | 66.94 % |
| Median UMI Counts per High Quality Cell | 11,648 |
| Median Genes per High Quality Cell | 2,994 |
| Total Genes Detected in High Quality Cell | 24,854 |
| Cell above Mitochondrial Expression Threshold | 8.13 % |
| Estimated Doublet Rate in High Quality Cell | 5.11 % |

#### Sequencing Stats

| | |
|---|---|
| Number of Reads Processed | 385,741,789 |
| Reads Pseudoaligned | 88.9 % |
| Reads on Whitelist | 94.66 % |
| Total UMI Counts | 204,120,352 |
| Sequencing Technology | 10xv3 |
| Species | Arabidopsis thaliana |
| Transcriptome | TAIR10 |

#### Sample Stats

| | |
|---|---|
| Sample | col0 |
| Name | WT Col-0 |
| Source | Benfey lab |
| Genotype | WT Col-0 |
| Transgene | NA |
| Treatment | untreated |
| Age | 5_day |
| Timepoint | NA |
| Rep | NA |
| Target Cells | 5,000 |
| Date | NA |
| Seq Run | NA |

#### UMI Counts Histogram

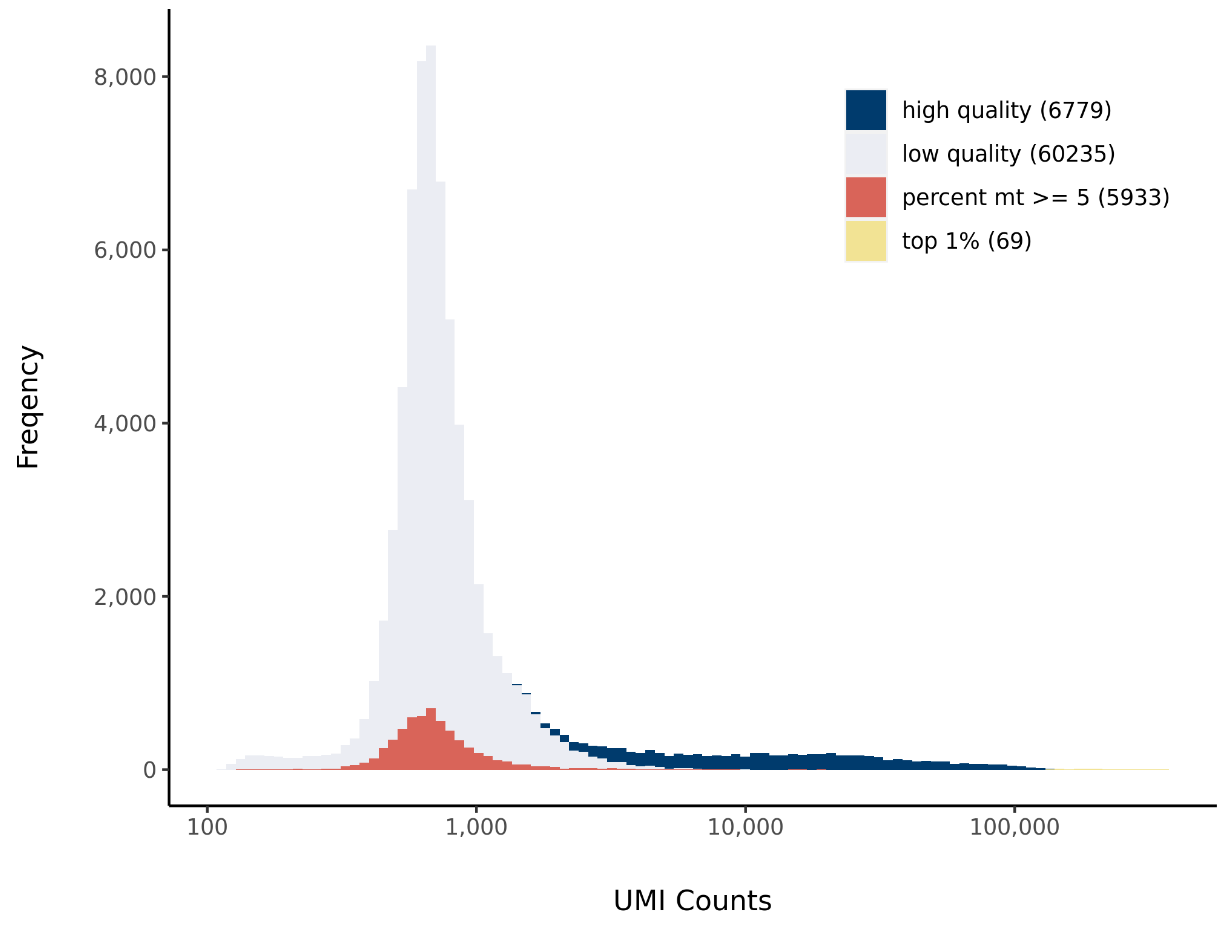


#### Number of Genes Histogram

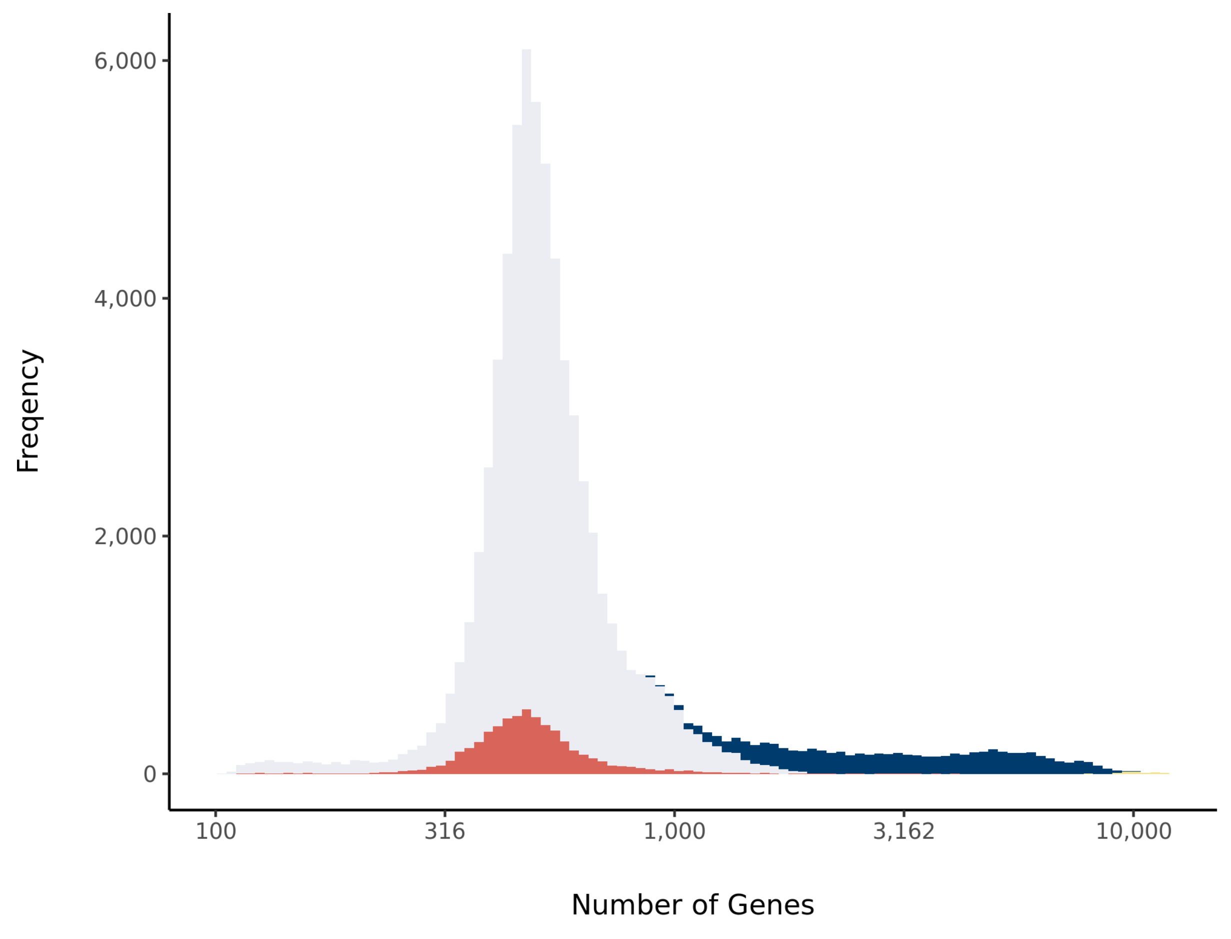


#### Barcode Rank Plot

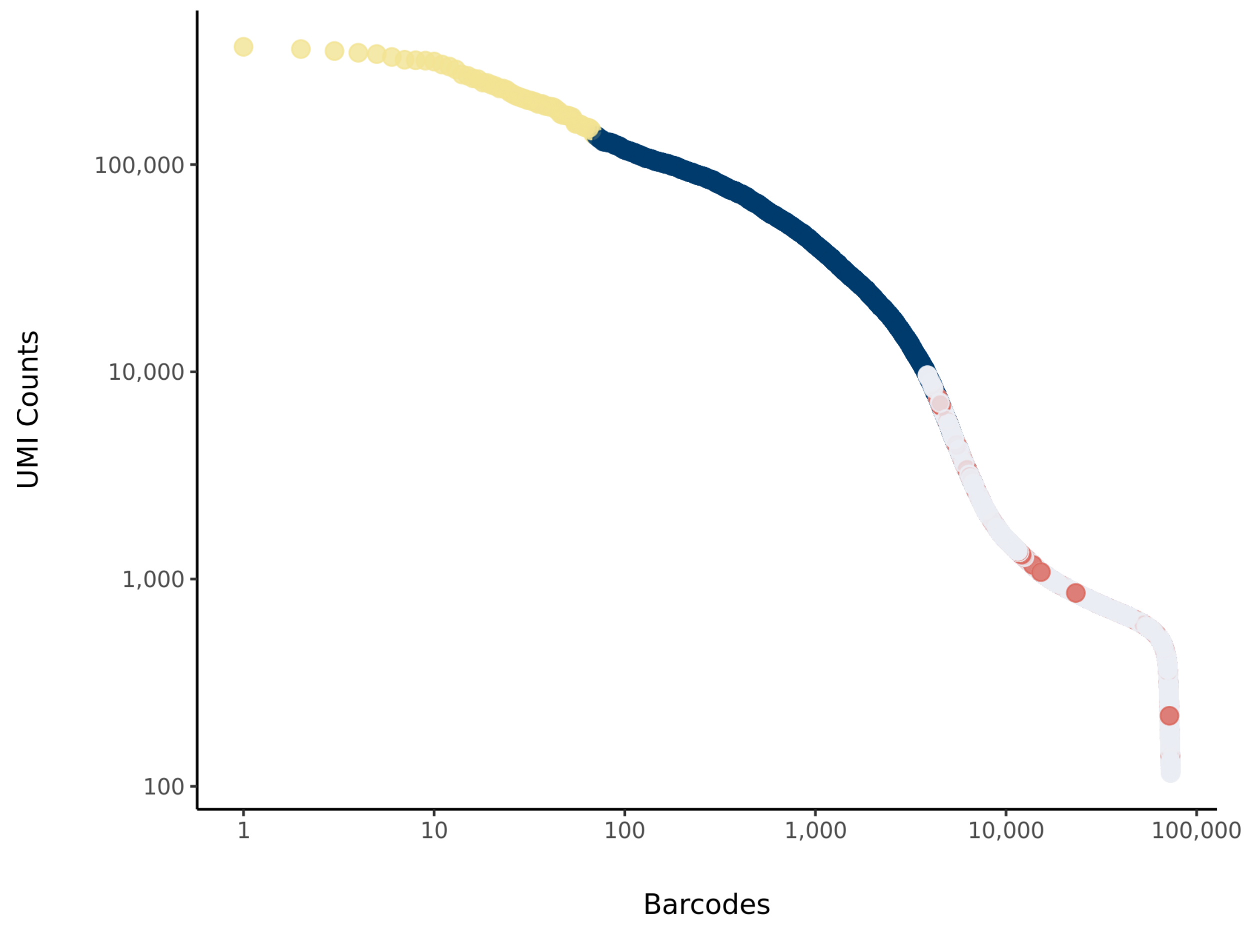

### dc1 Summary

Processed by COPILOT

Summary Analysis

#### Parameters

| | |
|---|---|
| Iteration of Filtering | 1 |
| Mitochondrial Expression Threshold | 5 % |
| Top High Quality Cell Filtered | 1 % |
| Doublet Removed | Yes |

#### Cell Stats

| | |
|---|---|
| Estimated Number of High Quality Cell | 3,471 |
| High Quality Cell | 4.65 % |
| Total UMI Counts in High Quality Cell | 68,731,526 |
| UMI Counts in High Quality Cell | 48.3 % |
| Median UMI Counts per High Quality Cell | 10,673 |
| Median Genes per High Quality Cell | 3,482 |
| Total Genes Detected in High Quality Cell | 25,036 |
| Cell above Mitochondrial Expression Threshold | 0.07 % |
| Estimated Doublet Rate in High Quality Cell | 2.68 % |

#### Sequencing Stats

| | |
|---|---|
| Number of Reads Processed | 261,013,034 |
| Reads Pseudoaligned | 90.1 % |
| Reads on Whitelist | 95.61 % |
| Total UMI Counts | 142,313,355 |
| Sequencing Technology | 10xv2 |
| Species | Arabidopsis thaliana |
| Transcriptome | TAIR10 |

#### Sample Stats

| | |
|---|---|
| Sample | dc1 |
| Name | WT Developmental Cell 1 |
| Source | Denyer et al. 2019, Developmental Cell |
| Genotype | WT Col-0 |
| Transgene | NA |
| Treatment | Untreated |
| Age | 6_day |
| Timepoint | NA |
| Rep | 1 |
| Target Cells | NA |
| Date | NA |
| Seq Run | NA |

#### UMI Counts Histogram

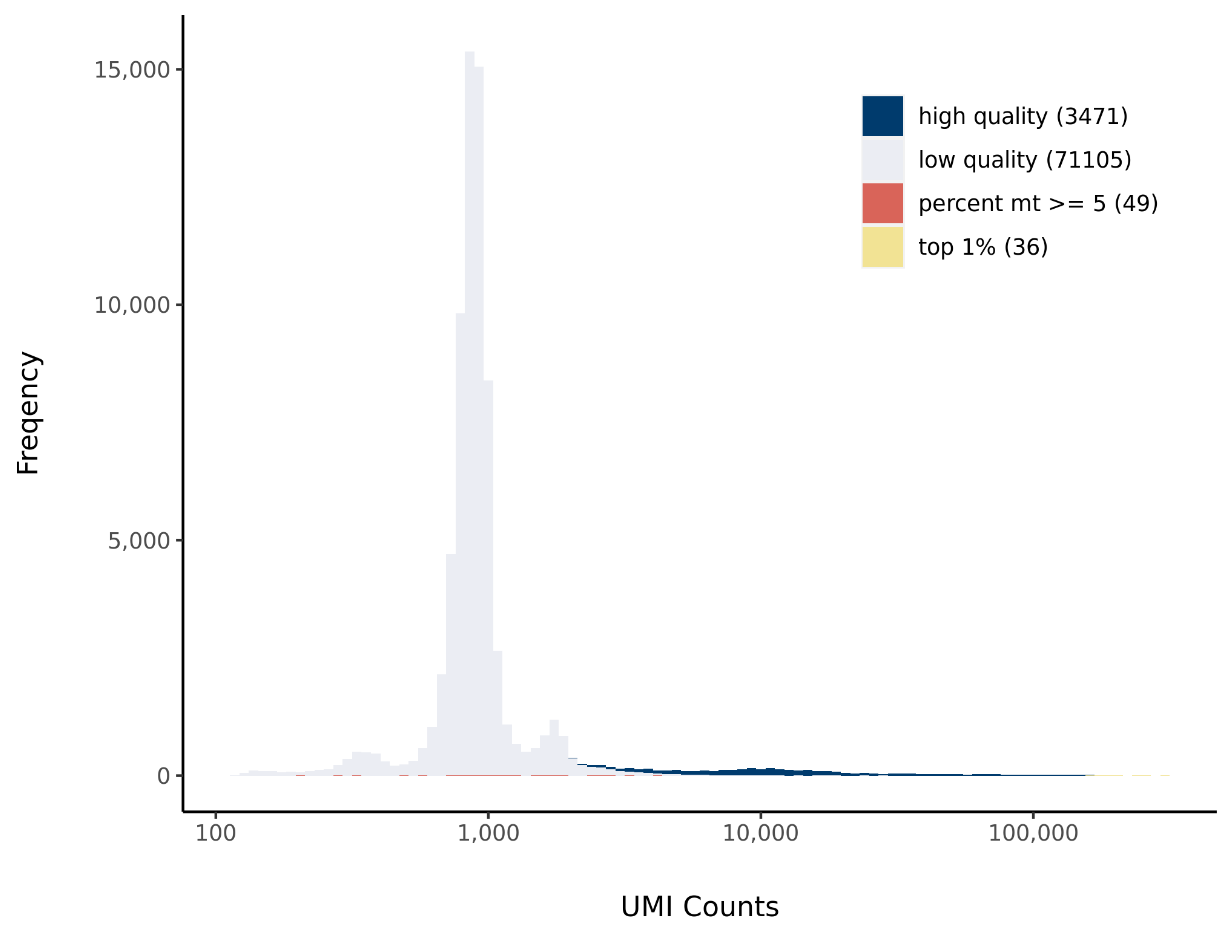


#### Number of Genes Histogram

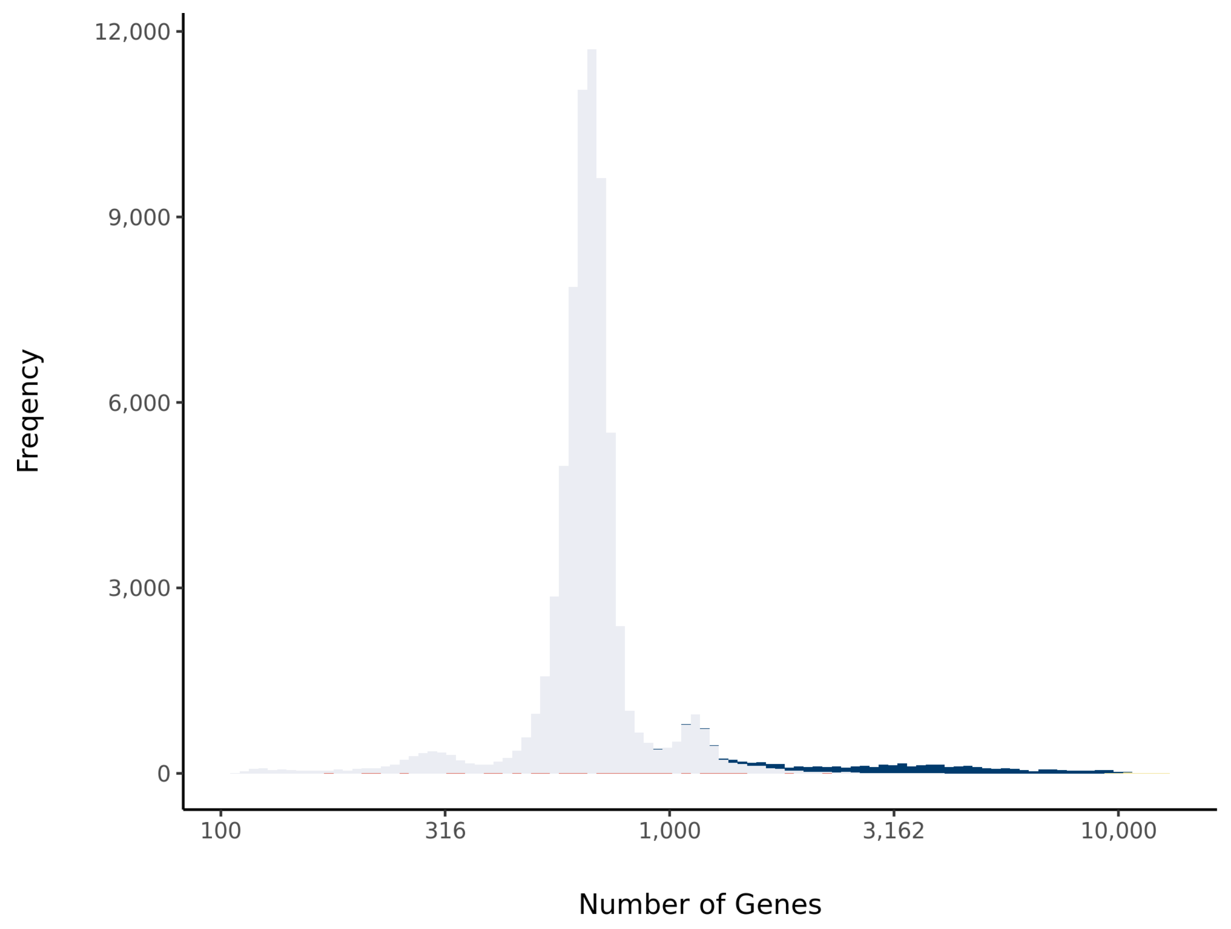


#### Barcode Rank Plot

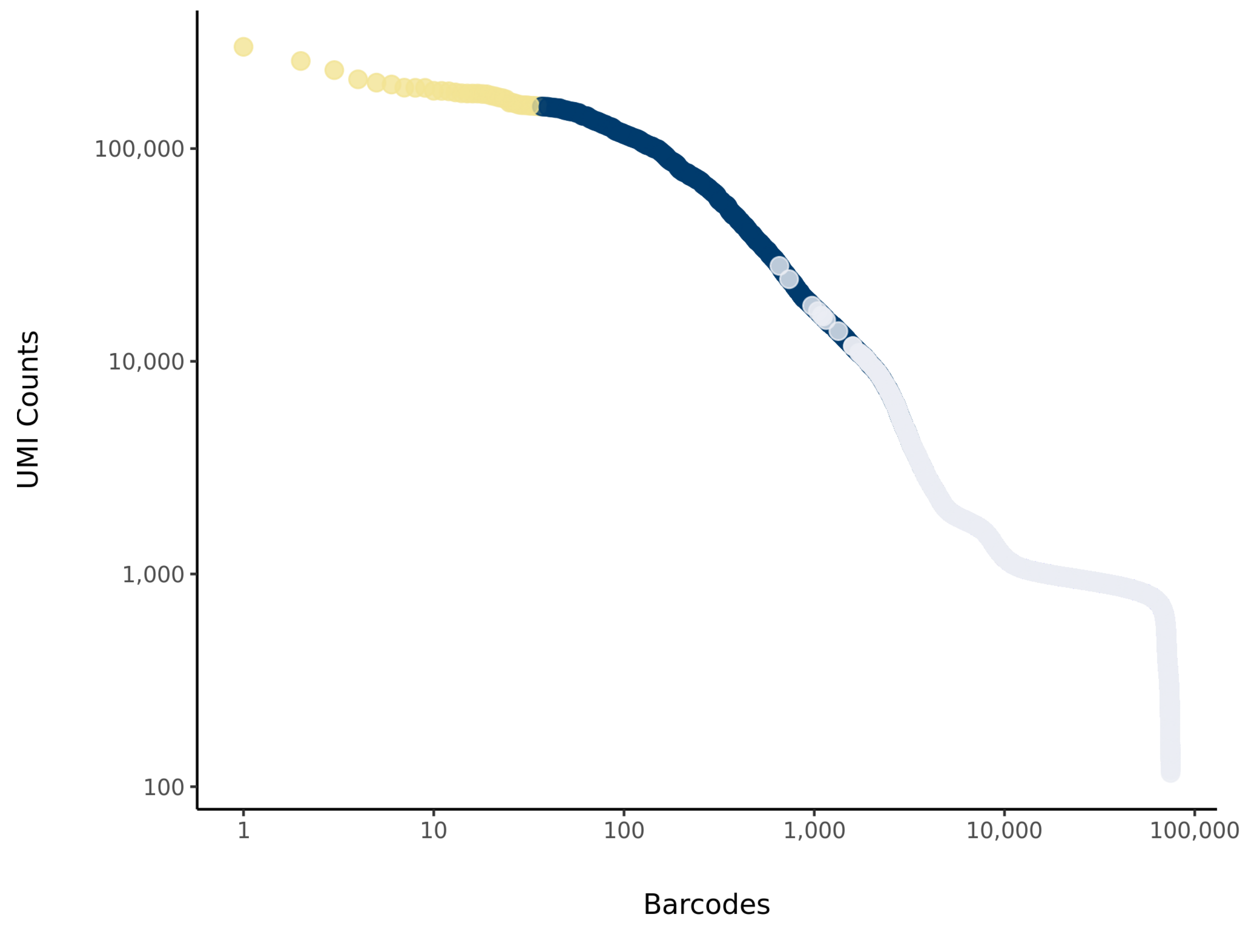

### dc2 Summary

Processed by COPILOT

Summary Analysis

#### Parameters

| | |
|---|---|
| Iteration of Filtering | 1 |
| Mitochondrial Expression Threshold | 5 % |
| Top High Quality Cell Filtered | 1 % |
| Doublet Removed | Yes |

#### Cell Stats

| | |
|---|---|
| Estimated Number of High Quality Cell | 3,371 |
| High Quality Cell | 4.56 % |
| Total UMI Counts in High Quality Cell | 68,781,391 |
| UMI Counts in High Quality Cell | 63.21 % |
| Median UMI Counts per High Quality Cell | 11,439 |
| Median Genes per High Quality Cell | 3,508 |
| Total Genes Detected in High Quality Cell | 25,100 |
| Cell above Mitochondrial Expression Threshold | 0.15 % |
| Estimated Doublet Rate in High Quality Cell | 2.61 % |

#### Sequencing Stats

| | |
|---|---|
| Number of Reads Processed | 235,863,595 |
| Reads Pseudoaligned | 89.6 % |
| Reads on Whitelist | 95.56 % |
| Total UMI Counts | 108,819,006 |
| Sequencing Technology | 10xv2 |
| Species | Arabidopsis thaliana |
| Transcriptome | TAIR10 |

#### Sample Stats

| | |
|---|---|
| Sample | dc2 |
| Name | WT Developmental Cell 2 |
| Source | Denyer et al. 2019, Developmental Cell |
| Genotype | WT Col-0 |
| Transgene | NA |
| Treatment | Untreated |
| Age | 6_day |
| Timepoint | NA |
| Rep | 2 |
| Target Cells | NA |
| Date | NA |
| Seq Run | NA |

#### UMI Counts Histogram

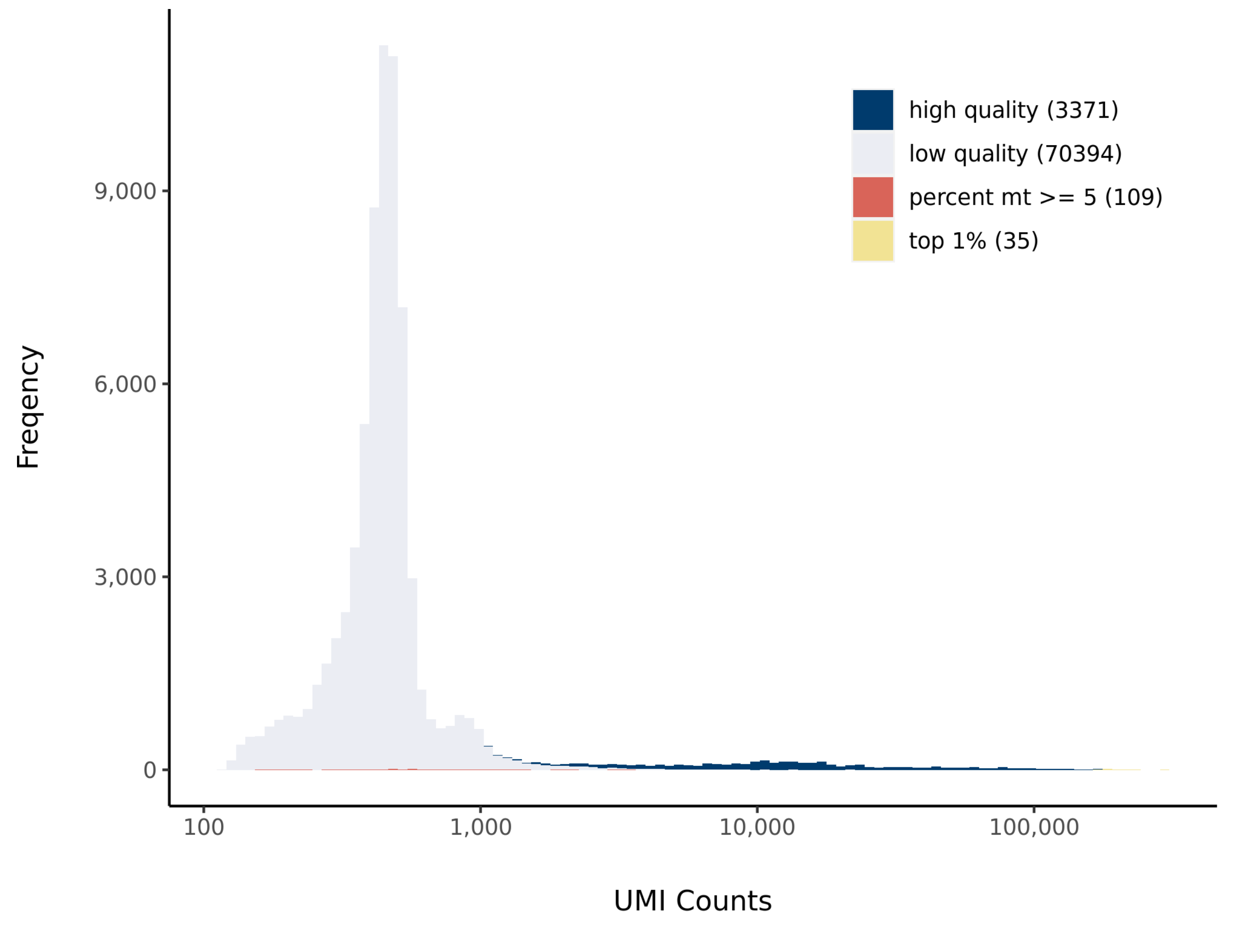


#### Number of Genes Histogram

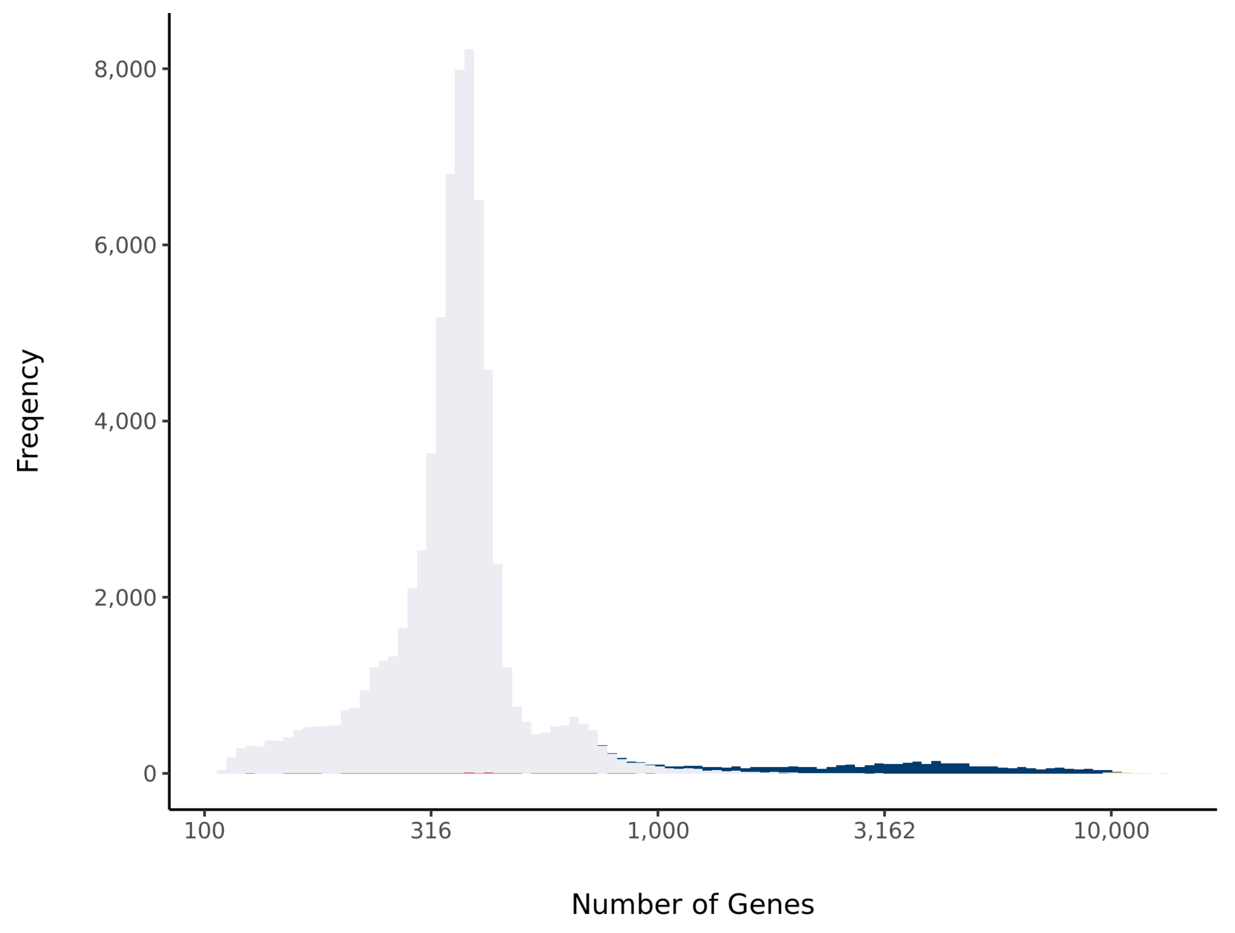


#### Barcode Rank Plot

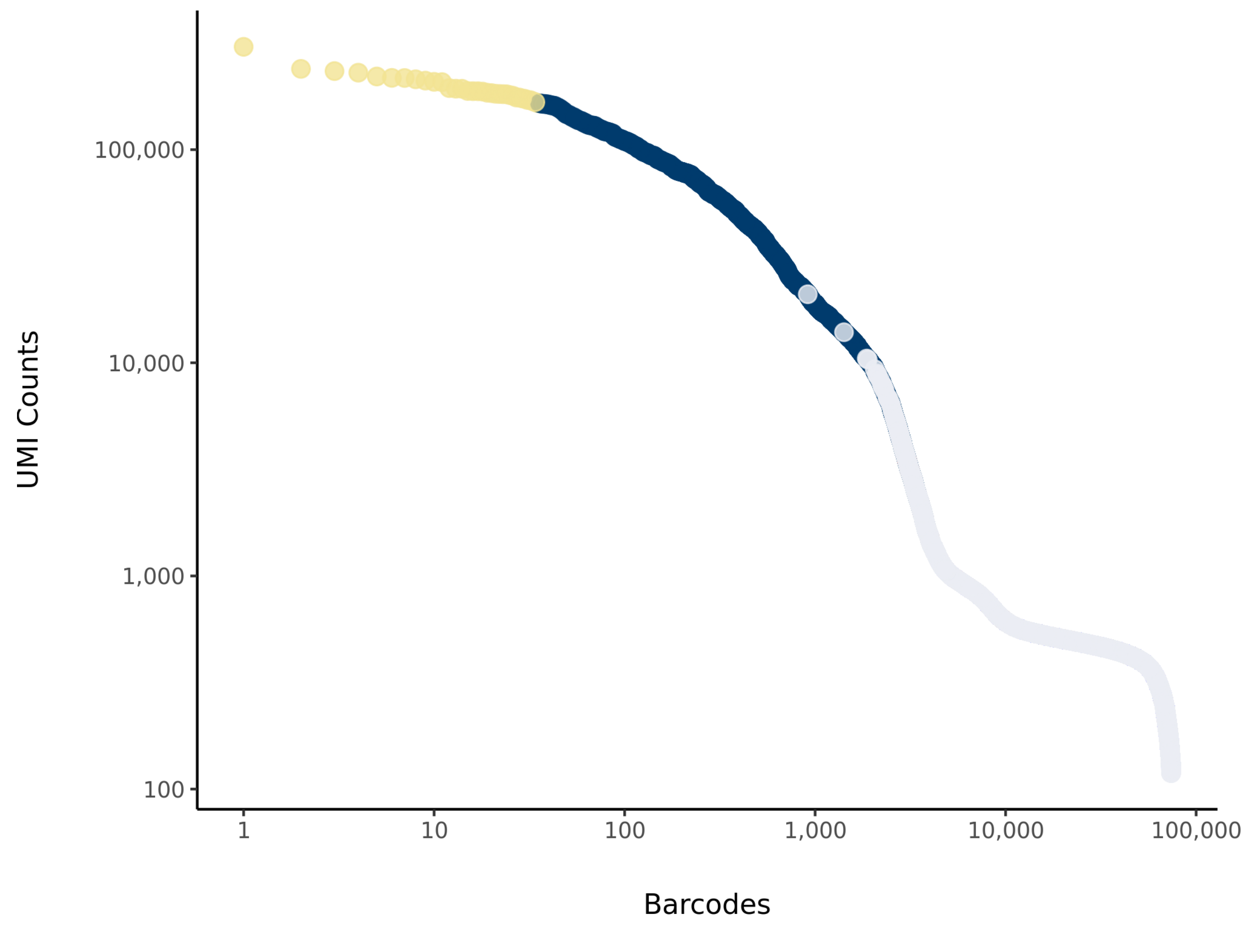

### pp1 Summary

Processed by COPILOT

Summary Analysis

#### Parameters

| | |
|---|---|
| Iteration of Filtering | 1 |
| Mitochondrial Expression Threshold | 5 % |
| Top High Quality Cell Filtered | 1 % |
| Doublet Removed | Yes |

#### Cell Stats

| | |
|---|---|
| Estimated Number of High Quality Cell | 8,168 |
| High Quality Cell | 11.82 % |
| Total UMI Counts in High Quality Cell | 165,283,937 |
| UMI Counts in High Quality Cell | 72.54 % |
| Median UMI Counts per High Quality Cell | 13,299.5 |
| Median Genes per High Quality Cell | 3,777.5 |
| Total Genes Detected in High Quality Cell | 25,625 |
| Cell above Mitochondrial Expression Threshold | 0.05 % |
| Estimated Doublet Rate in High Quality Cell | 6.13 % |

#### Sequencing Stats

| | |
|---|---|
| Number of Reads Processed | 377,721,321 |
| Reads Pseudoaligned | 93.6 % |
| Reads on Whitelist | 96.53 % |
| Total UMI Counts | 227,847,045 |
| Sequencing Technology | 10xv2 |
| Species | Arabidopsis thaliana |
| Transcriptome | TAIR10 |

#### Sample Stats

| | |
|---|---|
| Sample | pp1 |
| Name | WT Plant Physiology 1 |
| Source | Ryu et al. 2019, Plant Physiology |
| Genotype | WT Col-0 |
| Transgene | NA |
| Treatment | Untreated |
| Age | 5_day |
| Timepoint | NA |
| Rep | 1 |
| Target Cells | NA |
| Date | NA |
| Seq Run | NA |

#### UMI Counts Histogram

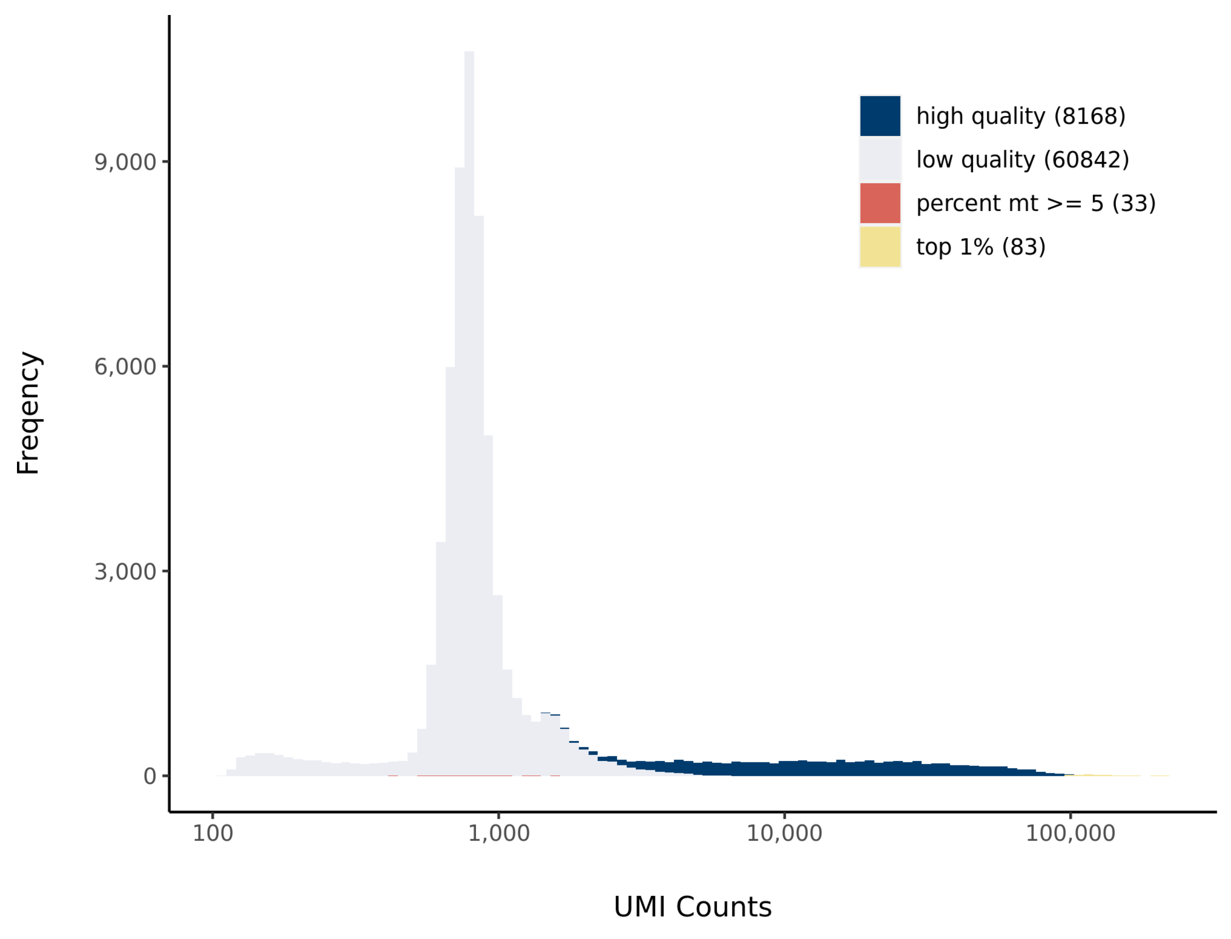


#### Number of Genes Histogram

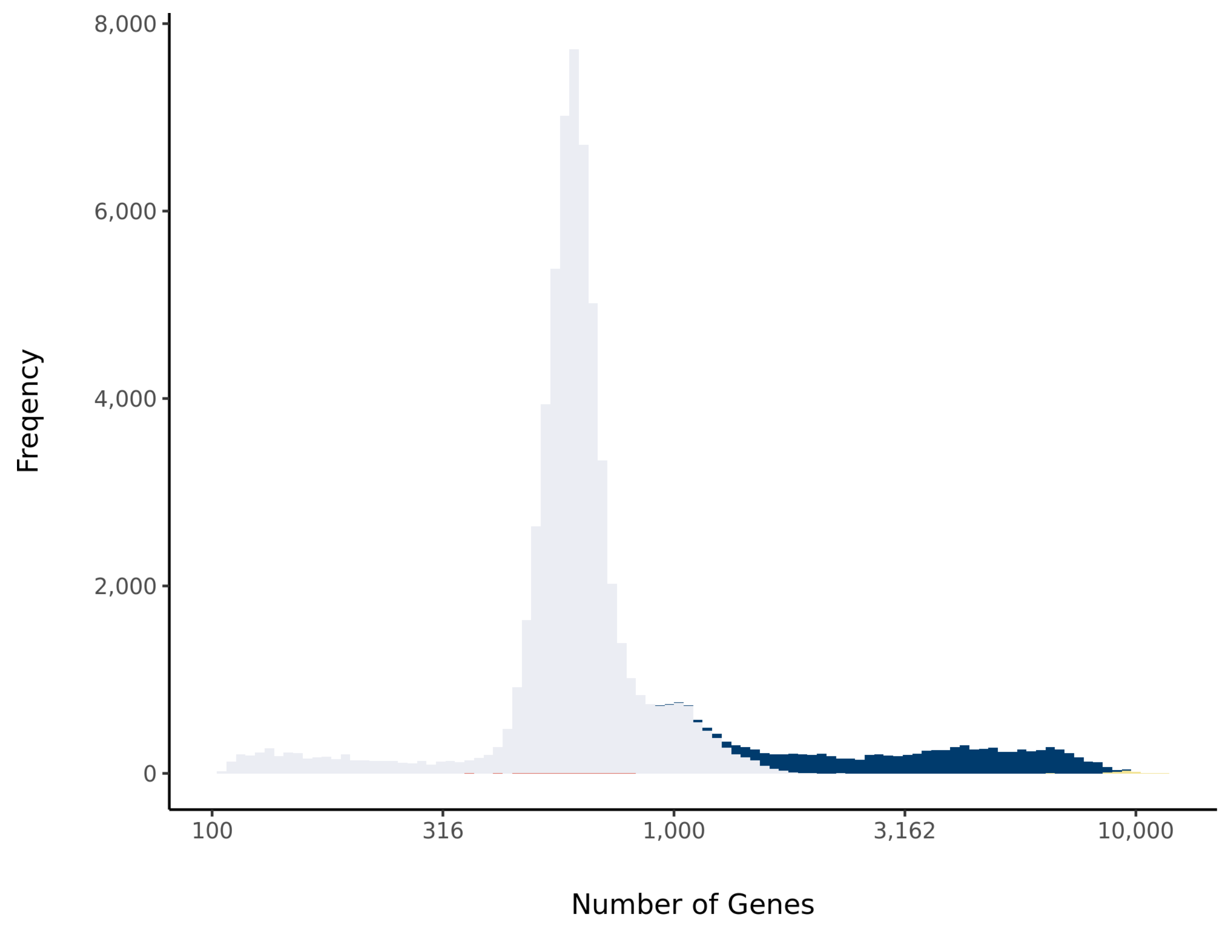


#### Barcode Rank Plot

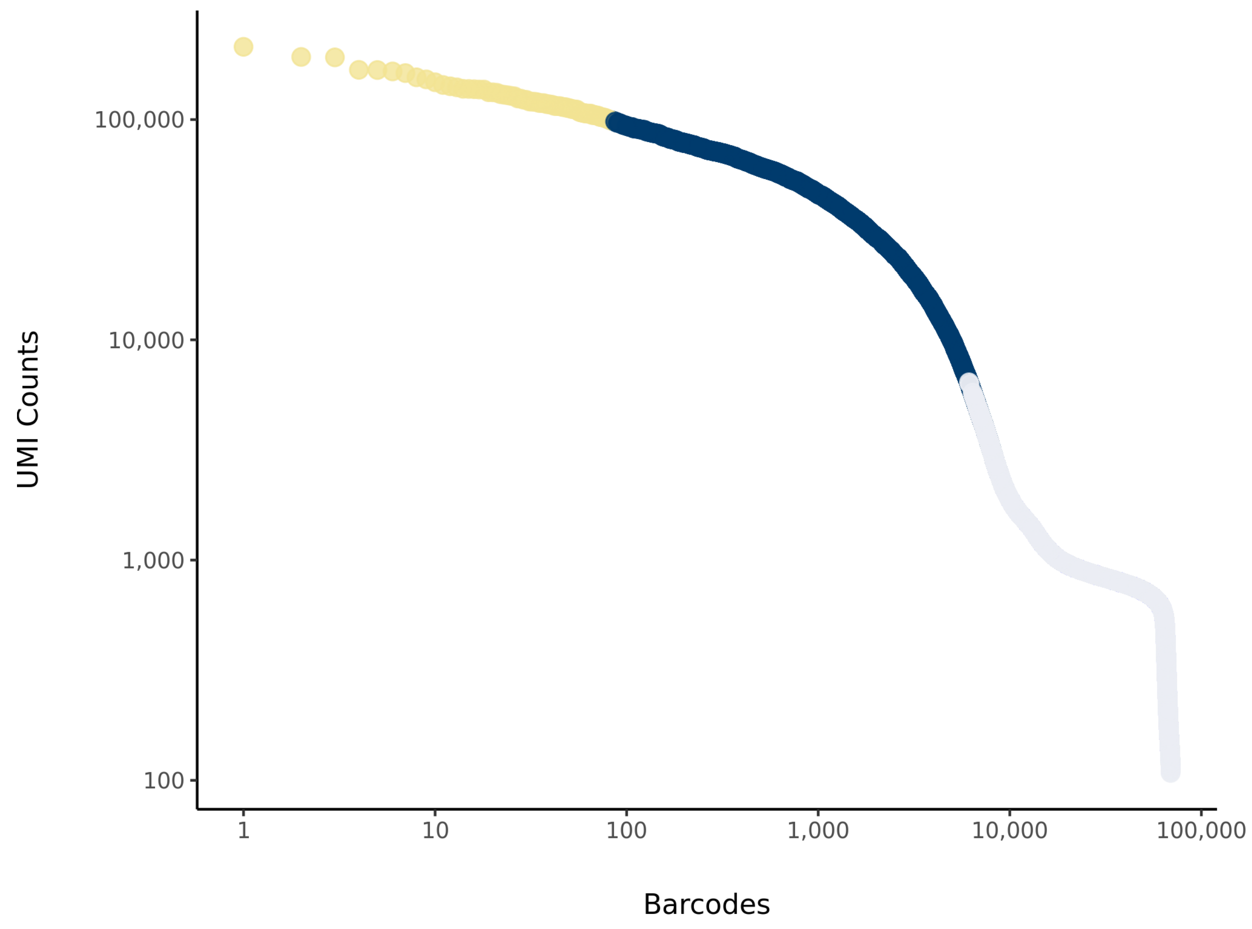

### sc_1 Summary

Processed by COPILOT

Summary    Analysis

#### Parameters

| | |
|---|---|
| Iteration of Filtering | 1 |
| Mitochondrial Expression Threshold | 5 % |
| Top High Quality Cell Filtered | 1 % |
| Doublet Removed | Yes |

#### Cell Stats

| | |
|---|---|
| Estimated Number of High Quality Cell | 10,598 |
| High Quality Cell | 14.68 % |
| Total UMI Counts in High Quality Cell | 79,958,429 |
| UMI Counts in High Quality Cell | 67.1 % |
| Median UMI Counts per High Quality Cell | 3,079 |
| Median Genes per High Quality Cell | 1,526 |
| Total Genes Detected in High Quality Cell | 24,624 |
| Cell above Mitochondrial Expression Threshold | 7.47 % |
| Estimated Doublet Rate in High Quality Cell | 7.92 % |

#### Sequencing Stats

| | |
|---|---|
| Number of Reads Processed | 255,953,705 |
| Reads Pseudoaligned | 92.4 % |
| Reads on Whitelist | 95.01 % |
| Total UMI Counts | 119,154,816 |
| Sequencing Technology | 10xv3 |
| Species | Arabidopsis thaliana |
| Transcriptome | TAIR10 |

#### Sample Stats

| | |
|---|---|
| Sample | sc_1 |
| Name | WT control |
| Source | Benfey lab |
| Genotype | WT Col-0 |
| Transgene | NA |
| Treatment | Untreated |
| Age | 7_day |
| Timepoint | 0 |
| Rep | 1 |
| Target Cells | 10,000 |
| Date | 2019-12-14 |
| Seq Run | Nolan_6131 |

#### UMI Counts Histogram

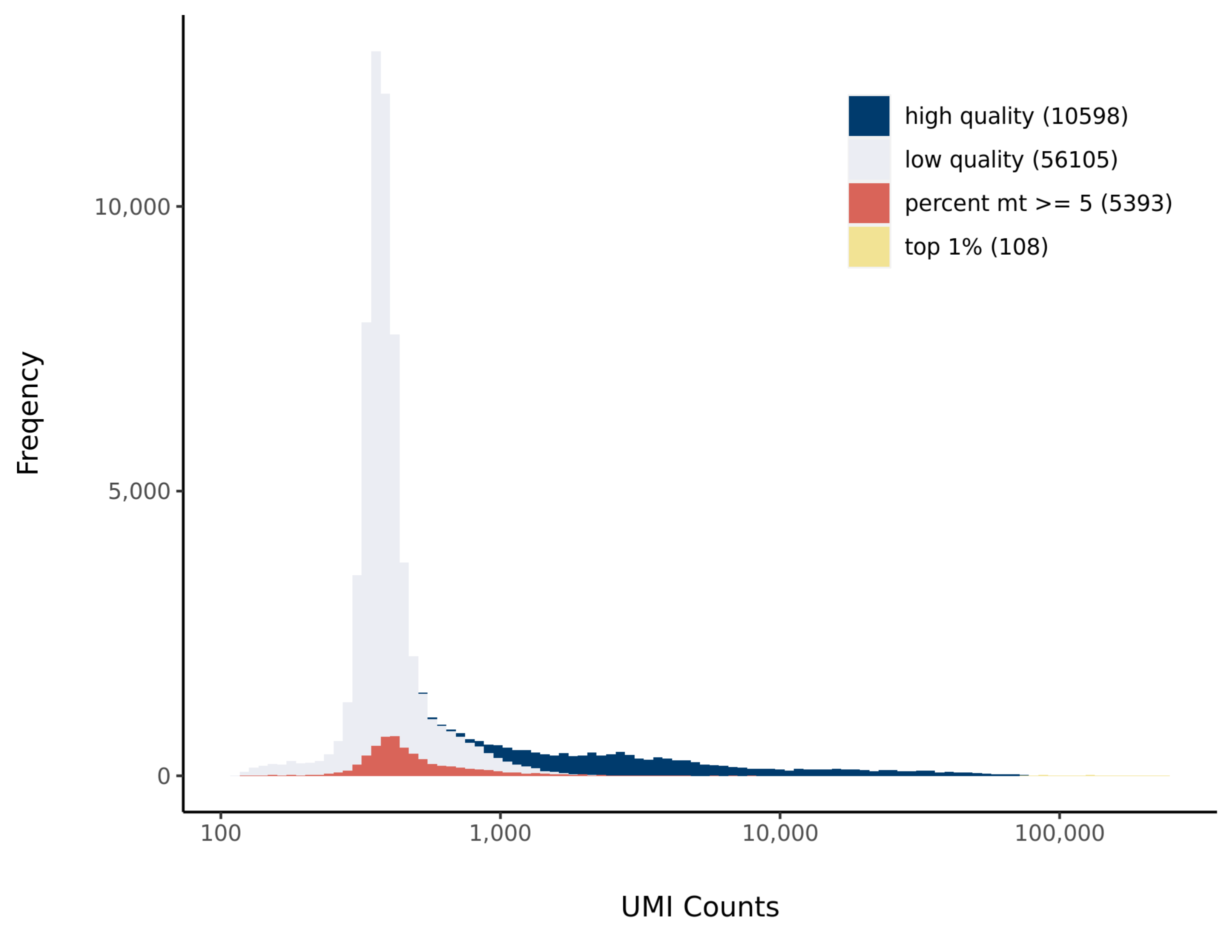


#### Number of Genes Histogram

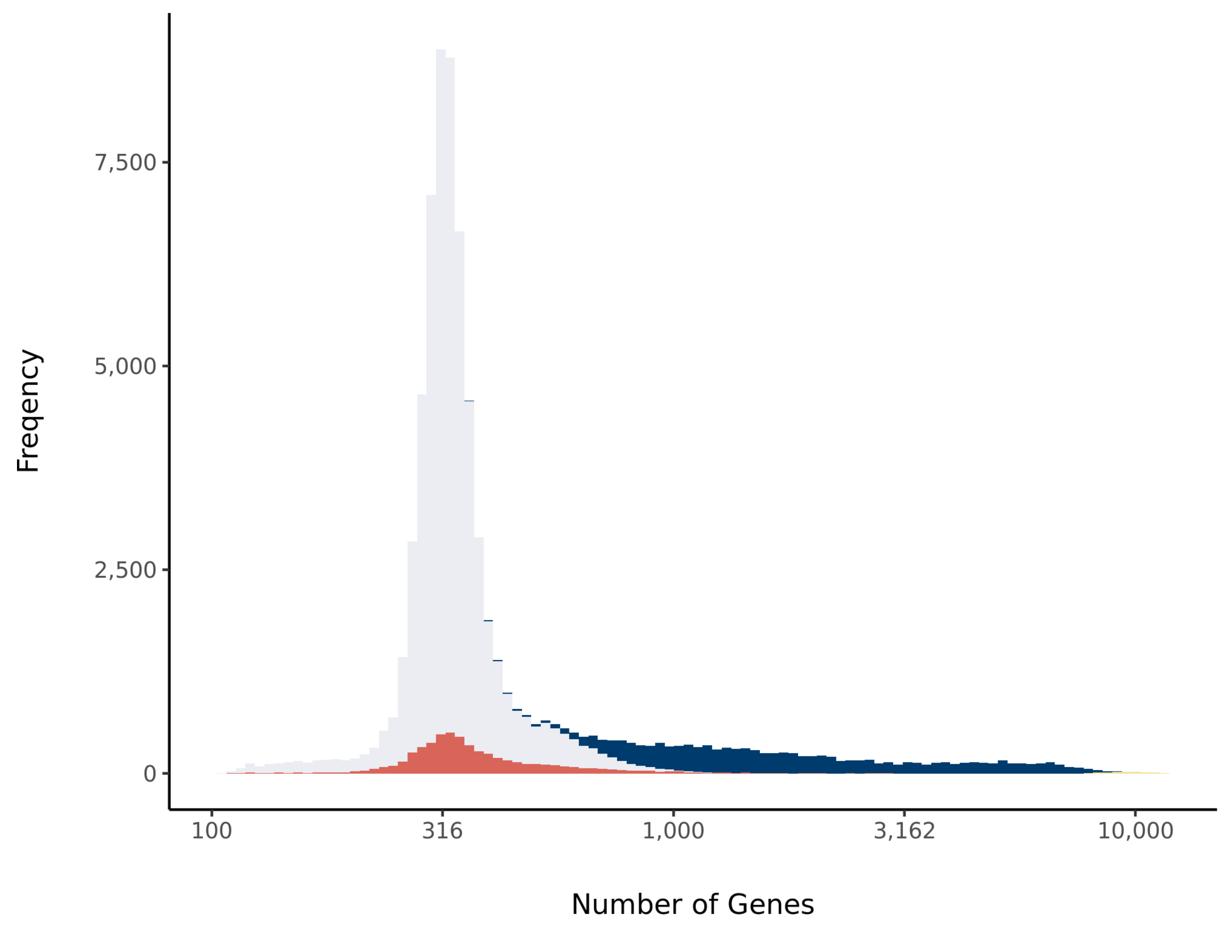


#### Barcode Rank Plot

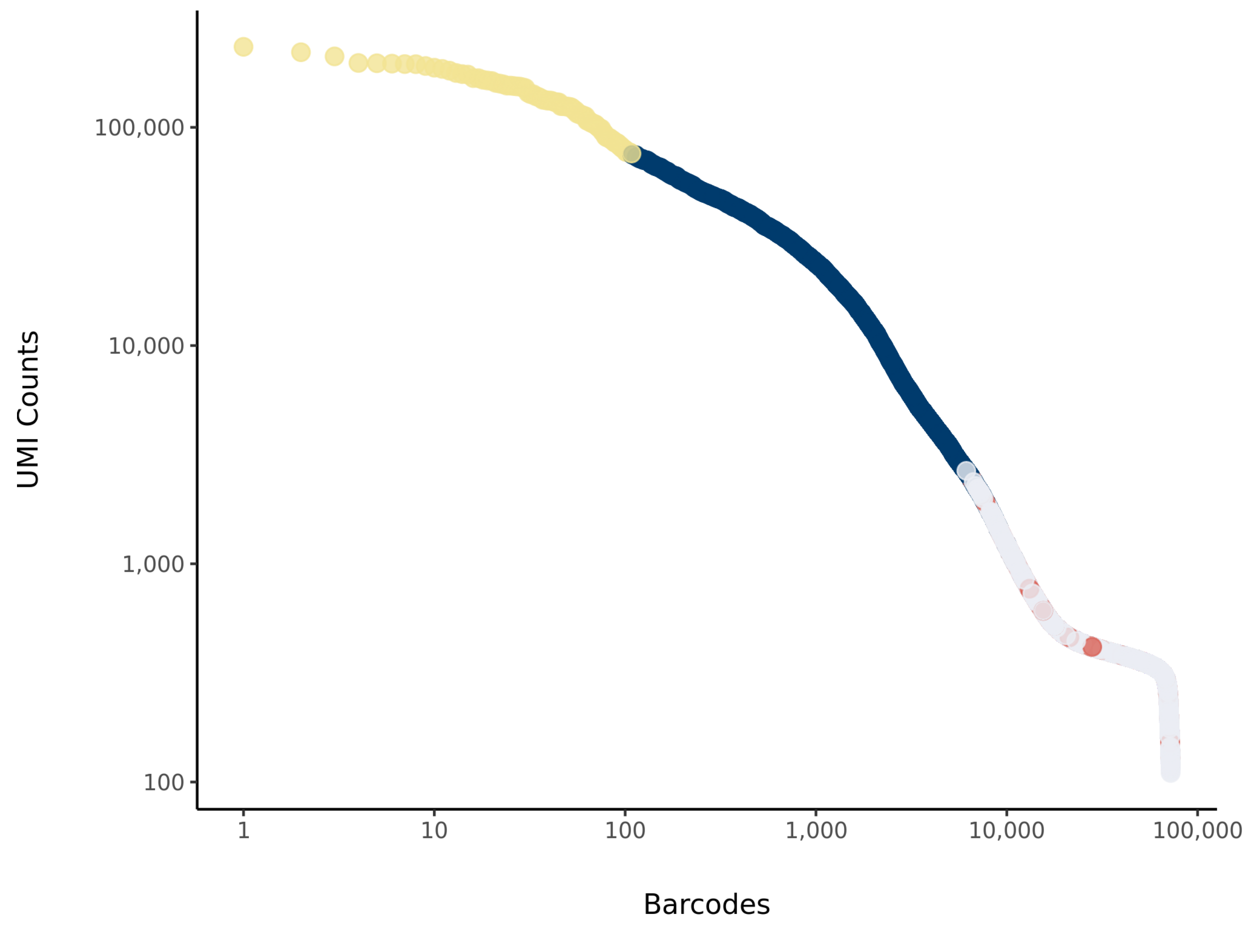

### sc_9_at Summary

Processed by COPILOT

Summary Analysis

#### Parameters

| | |
|---|---|
| Iteration of Filtering | 1 |
| Mitochondrial Expression Threshold | 5 % |
| Top High Quality Cell Filtered | 1 % |
| Doublet Removed | Yes |

#### Cell Stats

| | |
|---|---|
| Estimated Number of High Quality Cell | 3,926 |
| High Quality Cell | 5.9 % |
| Total UMI Counts in High Quality Cell | 67,227,229 |
| UMI Counts in High Quality Cell | 54.25 % |
| Median UMI Counts per High Quality Cell | 9,539 |
| Median Genes per High Quality Cell | 3,046 |
| Total Genes Detected in High Quality Cell | 24,999 |
| Cell above Mitochondrial Expression Threshold | 2.82 % |
| Estimated Doublet Rate in High Quality Cell | 3.01 % |

#### Sequencing Stats

| | |
|---|---|
| Number of Reads Processed | 271,063,644 |
| Reads Pseudoaligned | 91.8 % |
| Reads on Whitelist | 93.83 % |
| Total UMI Counts | 123,926,323 |
| Sequencing Technology | 10xv3 |
| Species | Arabidopsis thaliana |
| Transcriptome | TAIR10 |

#### Sample Stats

| | |
|---|---|
| Sample | sc_9 |
| Name | Rice 1 cm and Arabidopsis |
| Source | Benfey lab |
| Genotype | X.Kitaake; WT Col-0 |
| Transgene | NA |
| Treatment | Untreated |
| Age | 2_day |
| Timepoint | NA |
| Rep | NA |
| Target Cells | 10,000 |
| Date | 2019-12-18 |
| Seq Run | Nolan_6131 |

#### UMI Counts Histogram

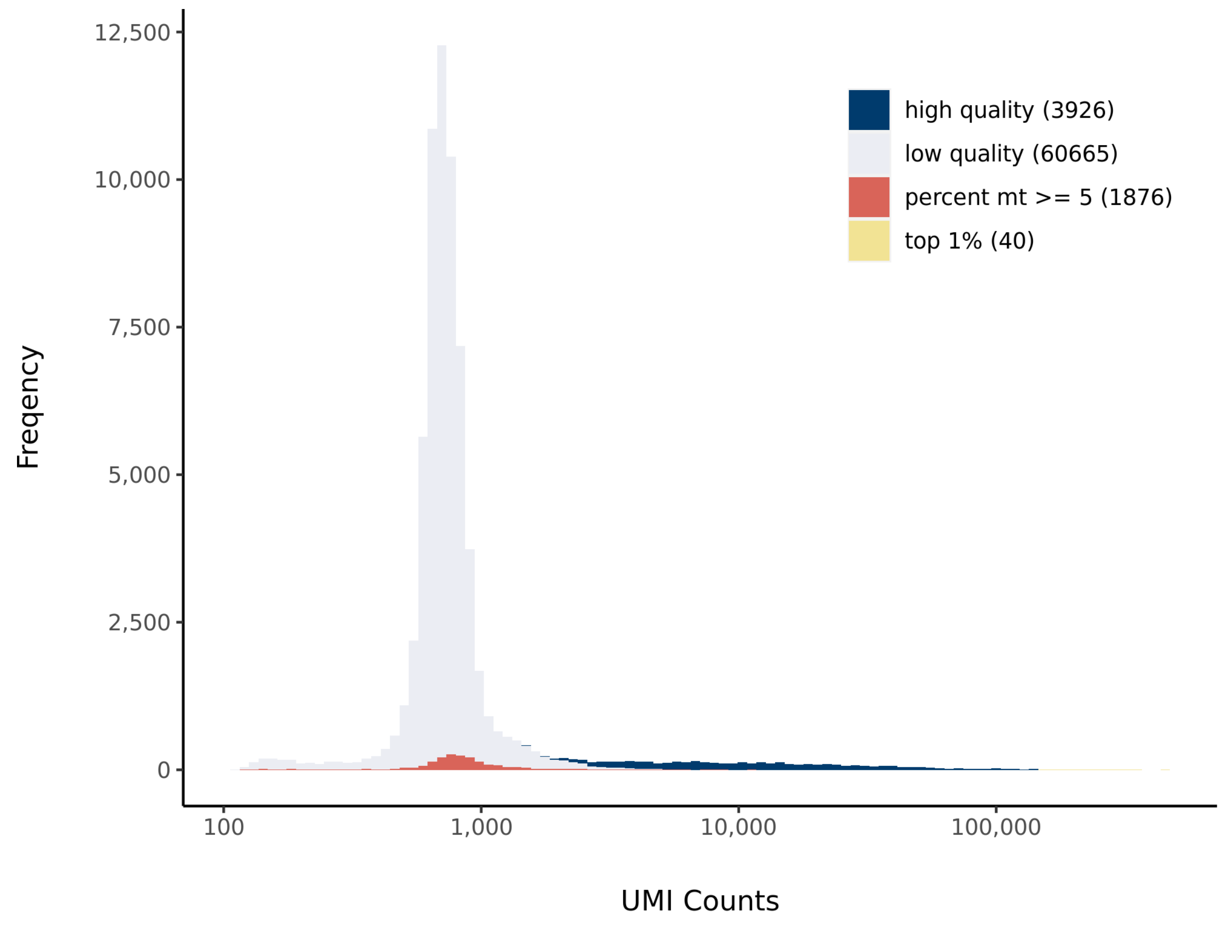


#### Number of Genes Histogram

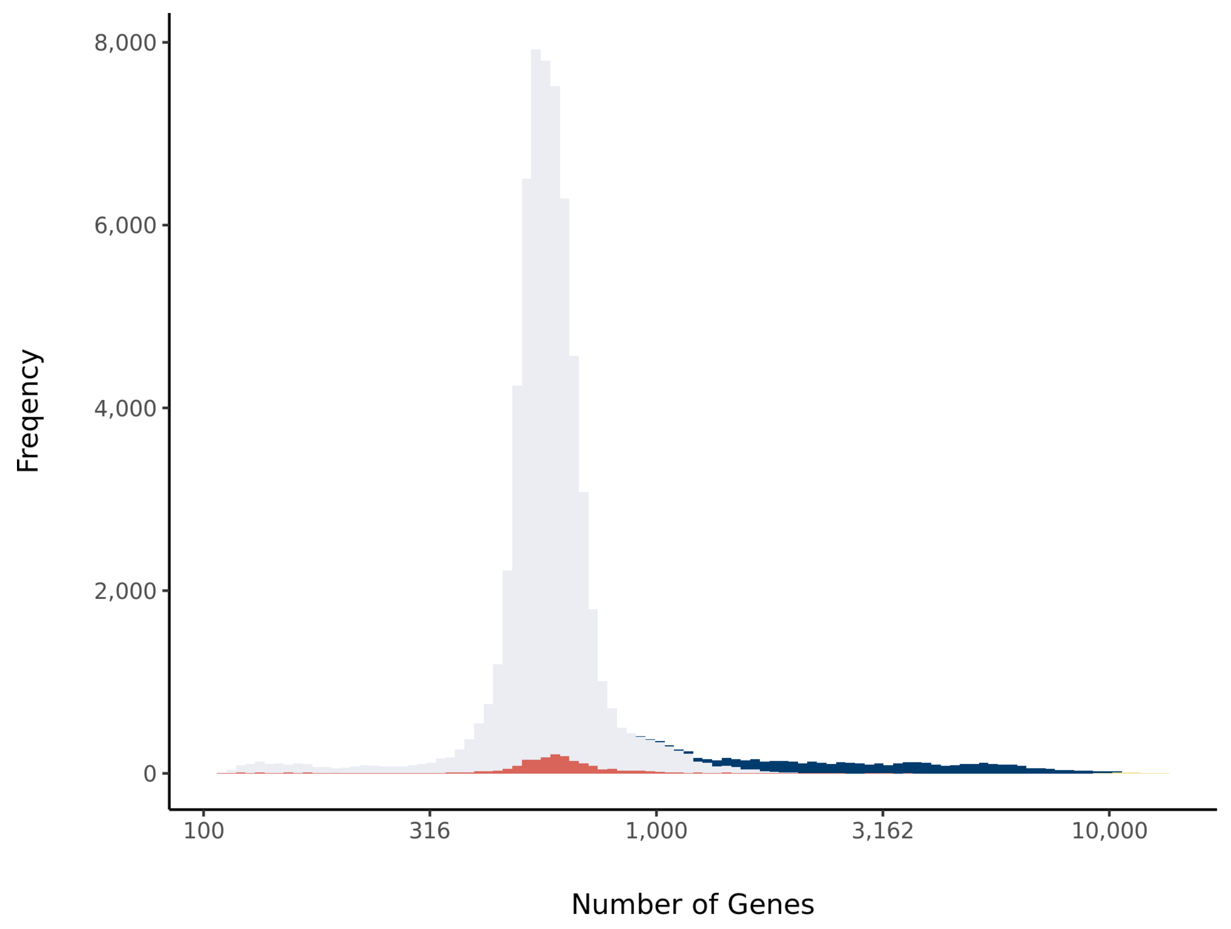


#### Barcode Rank Plot

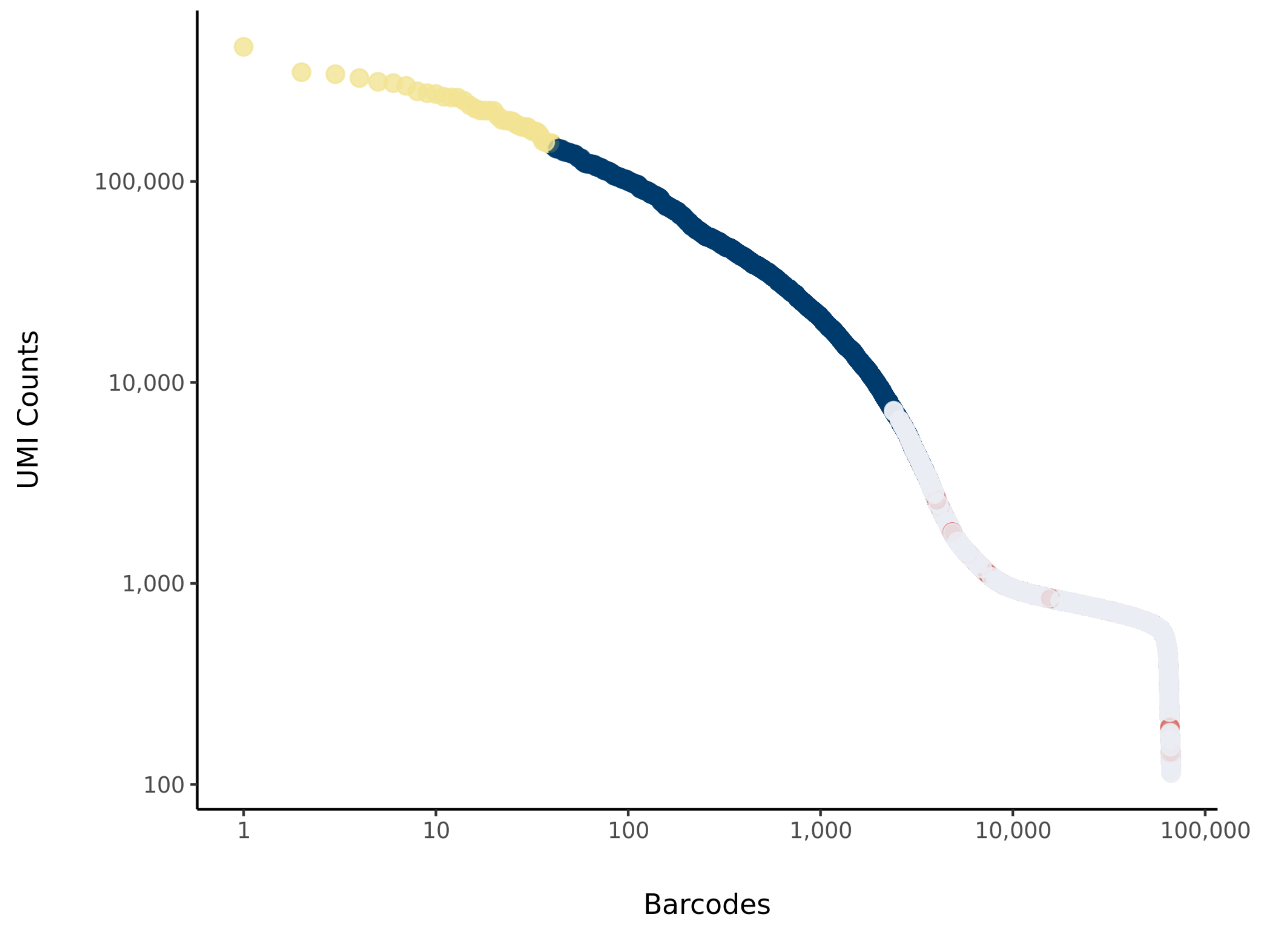

### sc_10_at Summary

Processed by COPILOT

Summary | Analysis

#### Parameters

| | |
|---|---|
| Iteration of Filtering | 1 |
| Mitochondrial Expression Threshold | 5 % |
| Top High Quality Cell Filtered | 1 % |
| Doublet Removed | Yes |

#### Cell Stats

| | |
|---|---|
| Estimated Number of High Quality Cell | 10,040 |
| High Quality Cell | 13.38 % |
| Total UMI Counts in High Quality Cell | 100,751,671 |
| UMI Counts in High Quality Cell | 75.06 % |
| Median UMI Counts per High Quality Cell | 5,553 |
| Median Genes per High Quality Cell | 2,279 |
| Total Genes Detected in High Quality Cell | 25,502 |
| Cell above Mitochondrial Expression Threshold | 69.11 % |
| Estimated Doublet Rate in High Quality Cell | 7.51 % |

#### Sequencing Stats

| | |
|---|---|
| Number of Reads Processed | 300,927,223 |
| Reads Pseudoaligned | 90.4 % |
| Reads on Whitelist | 92.36 % |
| Total UMI Counts | 134,231,998 |
| Sequencing Technology | 10xv3 |
| Species | Arabidopsis thaliana |
| Transcriptome | TAIR10 |

#### Sample Stats

| | |
|---|---|
| Sample | sc_10 |
| Name | Rice 1 cm and Arabidopsis |
| Source | Benfey lab |
| Genotype | X.Kitaake; WT Col-0 |
| Transgene | NA |
| Treatment | Untreated |
| Age | 2_day |
| Timepoint | NA |
| Rep | NA |
| Target Cells | 20,000 |
| Date | 2019-12-18 |
| Seq Run | Nolan_6131 |

#### UMI Counts Histogram

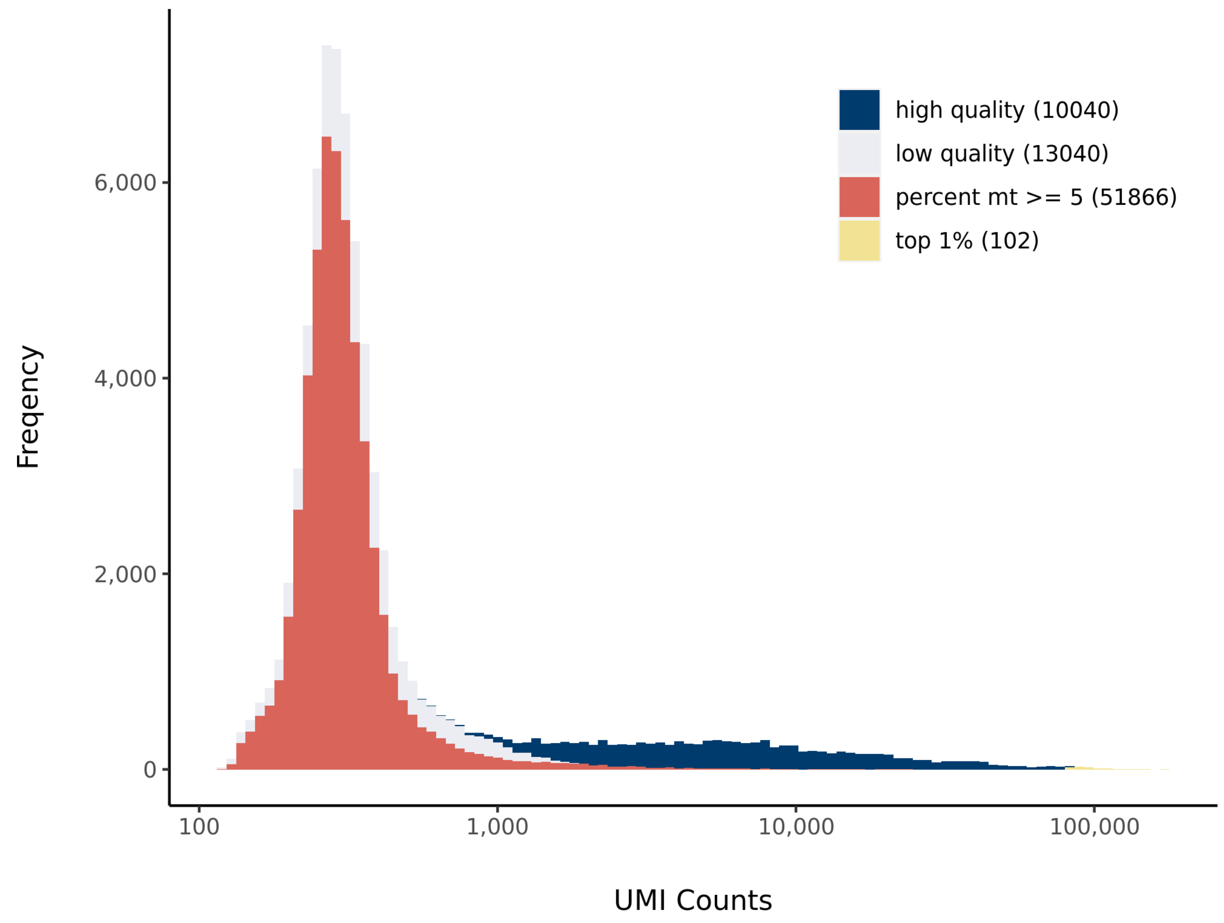


#### Number of Genes Histogram

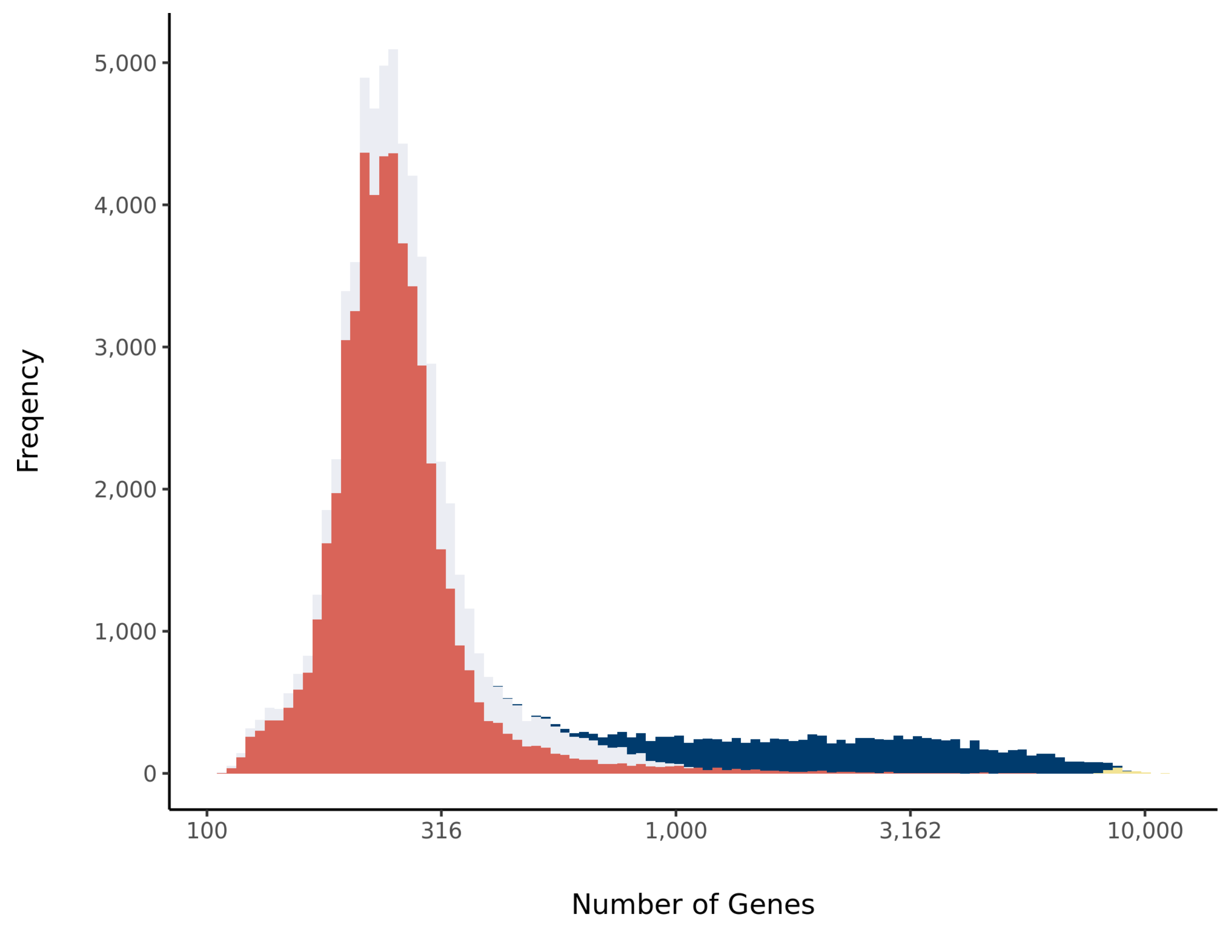


#### Barcode Rank Plot

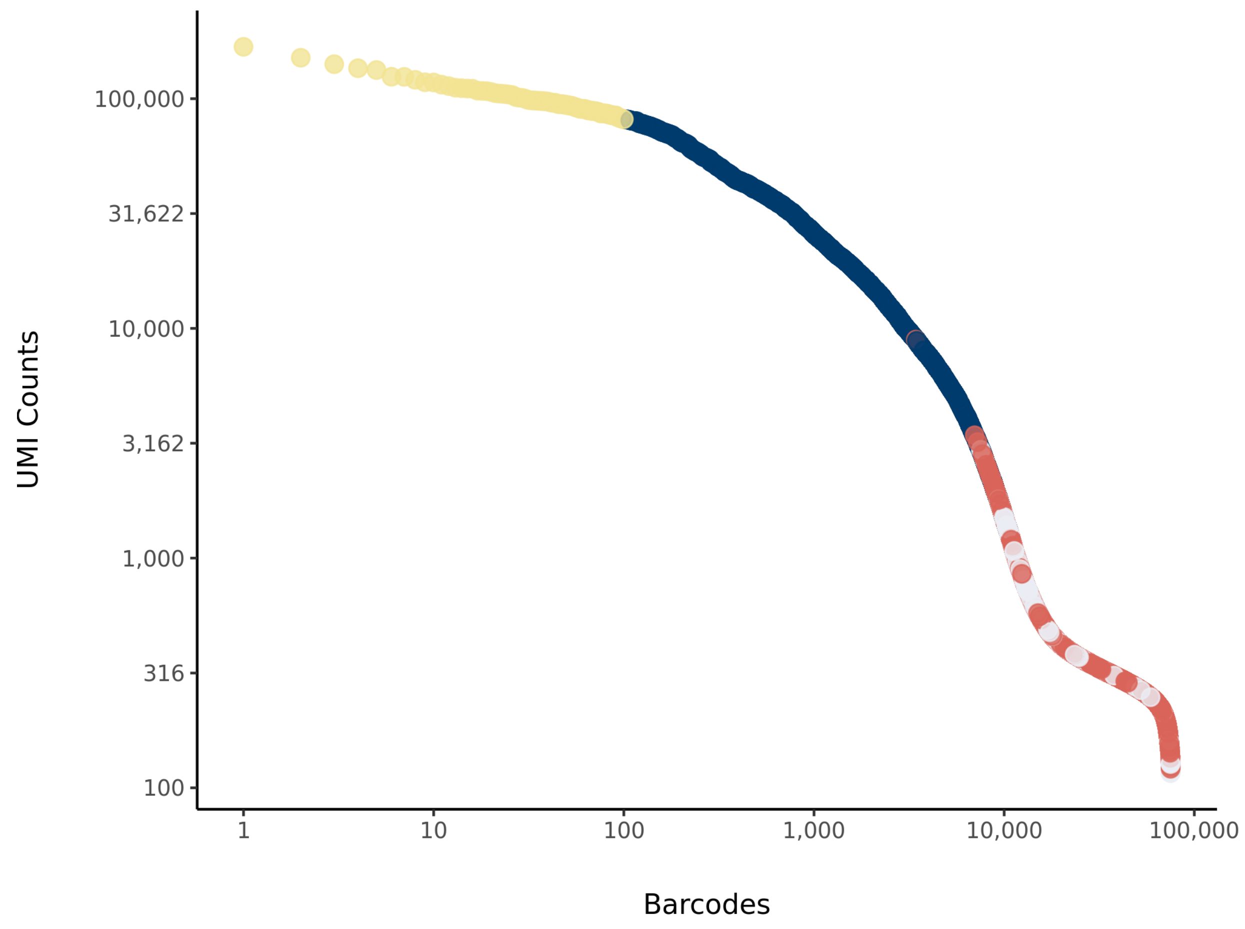

### sc_11 Summary

Processed by COPILOT

Summary | Analysis

#### Parameters

| | |
|---|---|
| Iteration of Filtering | 1 |
| Mitochondrial Expression Threshold | 5 % |
| Top High Quality Cell Filtered | 1 % |
| Doublet Removed | Yes |

#### Cell Stats

| | |
|---|---|
| Estimated Number of High Quality Cell | 9,913 |
| High Quality Cell | 13.96 % |
| Total UMI Counts in High Quality Cell | 168,126,295 |
| UMI Counts in High Quality Cell | 82.76 % |
| Median UMI Counts per High Quality Cell | 9,146 |
| Median Genes per High Quality Cell | 3,050 |
| Total Genes Detected in High Quality Cell | 25,040 |
| Cell above Mitochondrial Expression Threshold | 8.81 % |
| Estimated Doublet Rate in High Quality Cell | 7.42 % |

#### Sequencing Stats

| | |
|---|---|
| Number of Reads Processed | 329,949,998 |
| Reads Pseudoaligned | 93.6 % |
| Reads on Whitelist | 95.21 % |
| Total UMI Counts | 203,138,562 |
| Sequencing Technology | 10xv3 |
| Species | Arabidopsis thaliana |
| Transcriptome | TAIR10 |

#### Sample Stats

| | |
|---|---|
| Sample | sc_11 |
| Name | WT Col-0 untreated |
| Source | Benfey lab |
| Genotype | WT Col-0 |
| Transgene | NA |
| Treatment | Untreated |
| Age | 5_day |
| Timepoint | NA |
| Rep | NA |
| Target Cells | 10,000 |
| Date | 2019-12-20 |
| Seq Run | Nolan_6131 |

#### UMI Counts Histogram

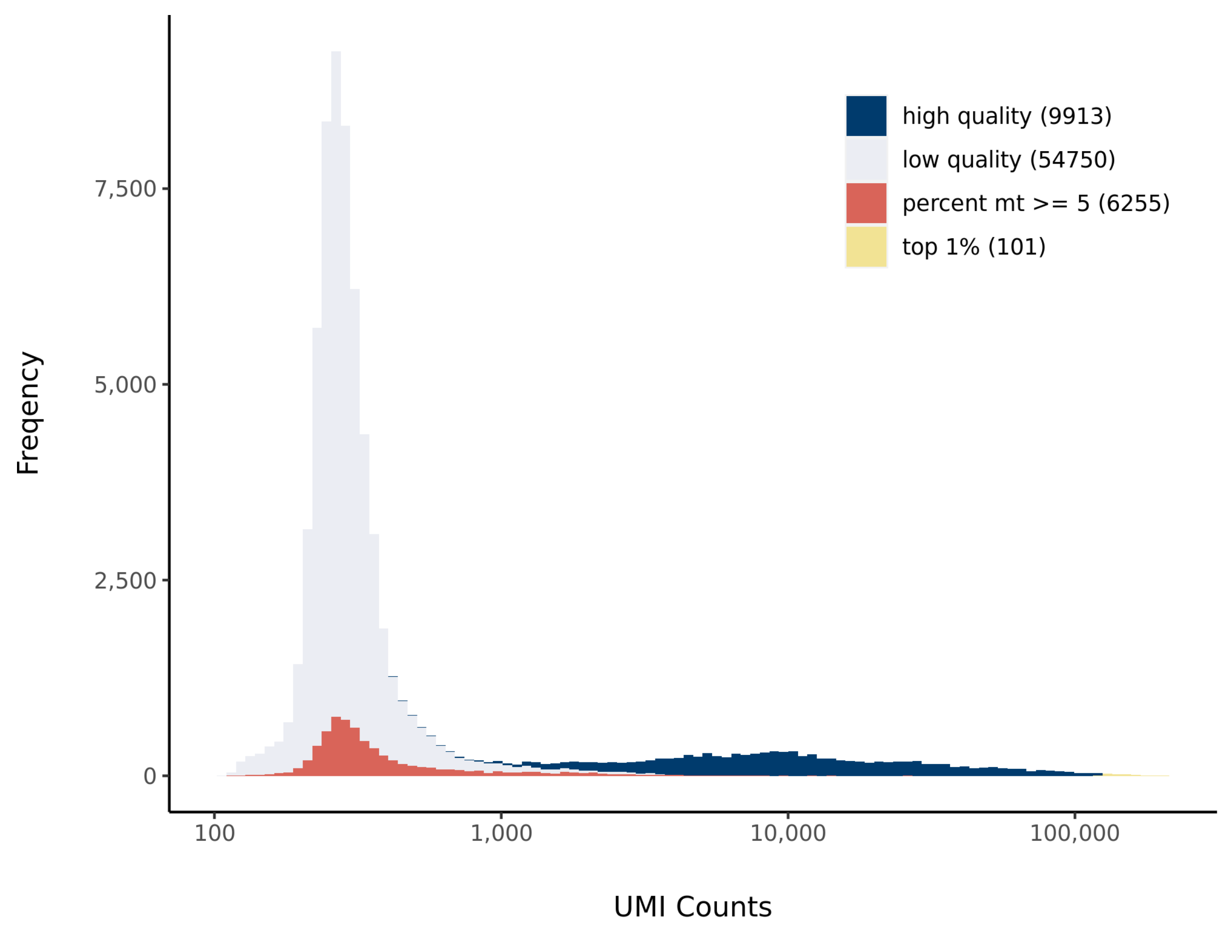


#### Number of Genes Histogram

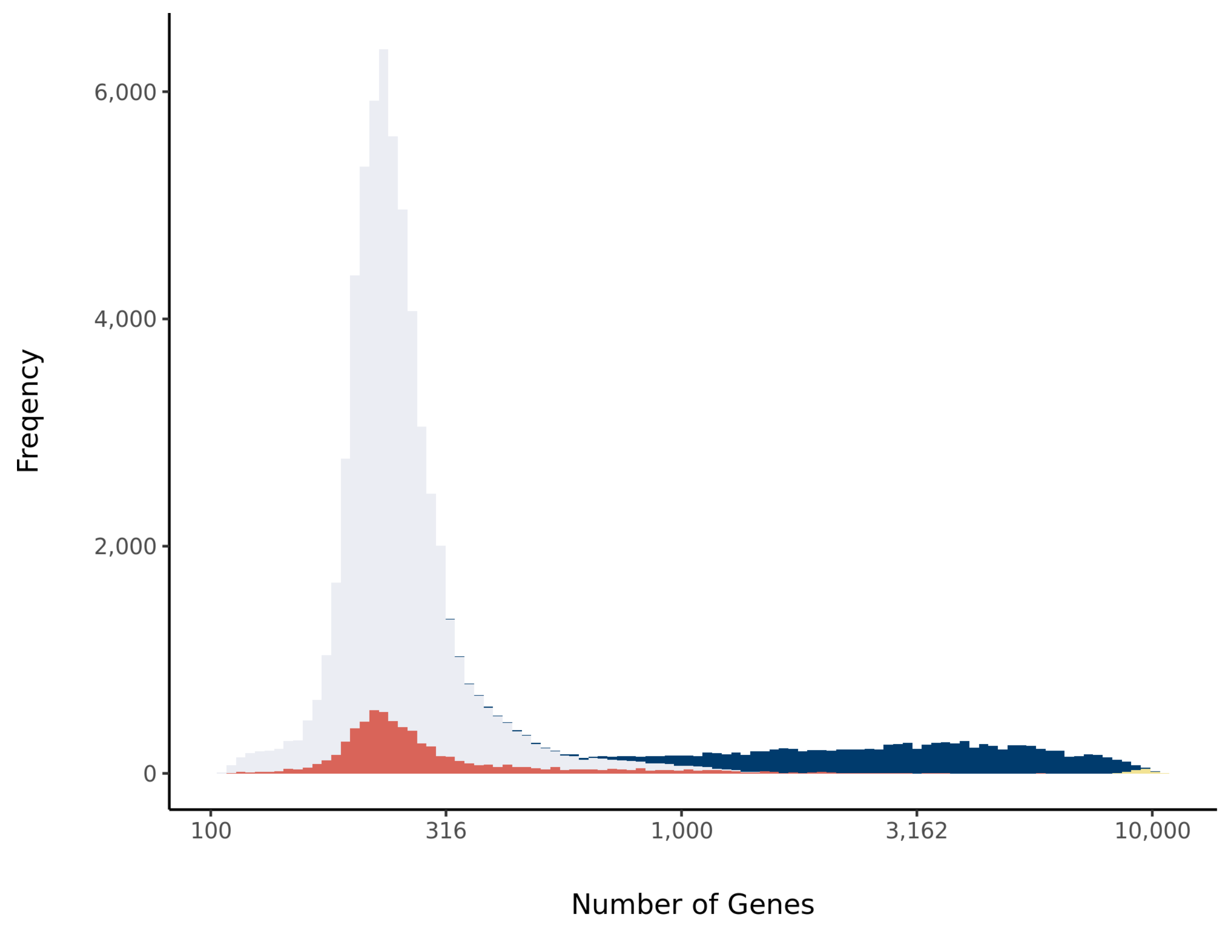


#### Barcode Rank Plot

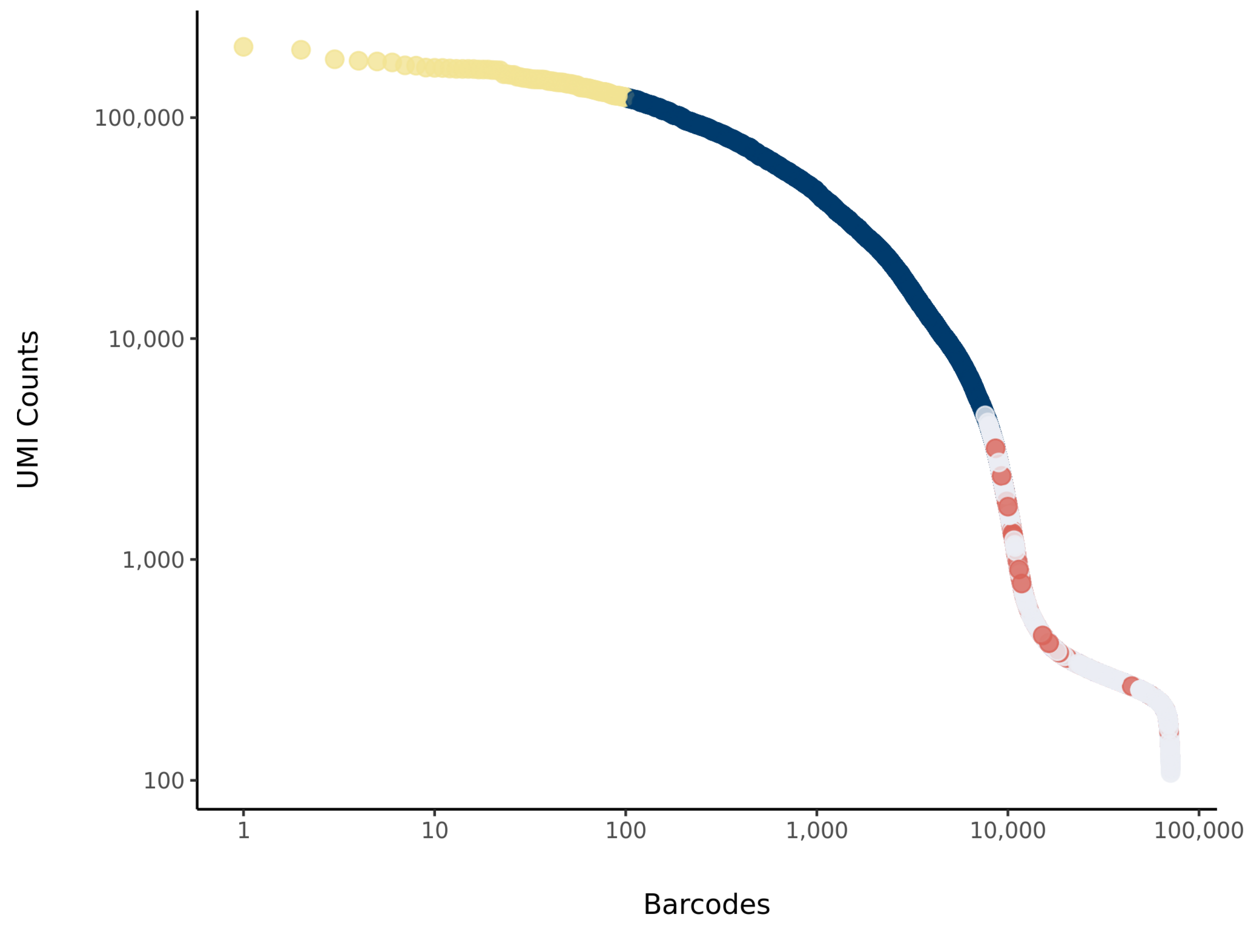

### sc_12 Summary

Processed by COPILOT

Summary Analysis

#### Parameters

| | |
|---|---|
| Iteration of Filtering | 1 |
| Mitochondrial Expression Threshold | 5 % |
| Top High Quality Cell Filtered | 1 % |
| Doublet Removed | Yes |

#### Cell Stats

| | |
|---|---|
| Estimated Number of High Quality Cell | 11,236 |
| High Quality Cell | 13.59 % |
| Total UMI Counts in High Quality Cell | 346,882,014 |
| UMI Counts in High Quality Cell | 83.04 % |
| Median UMI Counts per High Quality Cell | 20,788.5 |
| Median Genes per High Quality Cell | 4,774 |
| Total Genes Detected in High Quality Cell | 26,018 |
| Cell above Mitochondrial Expression Threshold | 6.77 % |
| Estimated Doublet Rate in High Quality Cell | 8.39 % |

#### Sequencing Stats

| | |
|---|---|
| Number of Reads Processed | 841,129,261 |
| Reads Pseudoaligned | 93.7 % |
| Reads on Whitelist | 96.04 % |
| Total UMI Counts | 417,731,033 |
| Sequencing Technology | 10xv3 |
| Species | Arabidopsis thaliana |
| Transcriptome | TAIR10 |

#### Sample Stats

| | |
|---|---|
| Sample | sc_12 |
| Name | WT Col-0 untreated |
| Source | Benfey lab |
| Genotype | WT Col-0 |
| Transgene | NA |
| Treatment | Untreated |
| Age | 5_day |
| Timepoint | NA |
| Rep | NA |
| Target Cells | 10,000 |
| Date | 2019-12-20 |
| Seq Run | Nolan_6131;Shahan_6158 |

#### UMI Counts Histogram

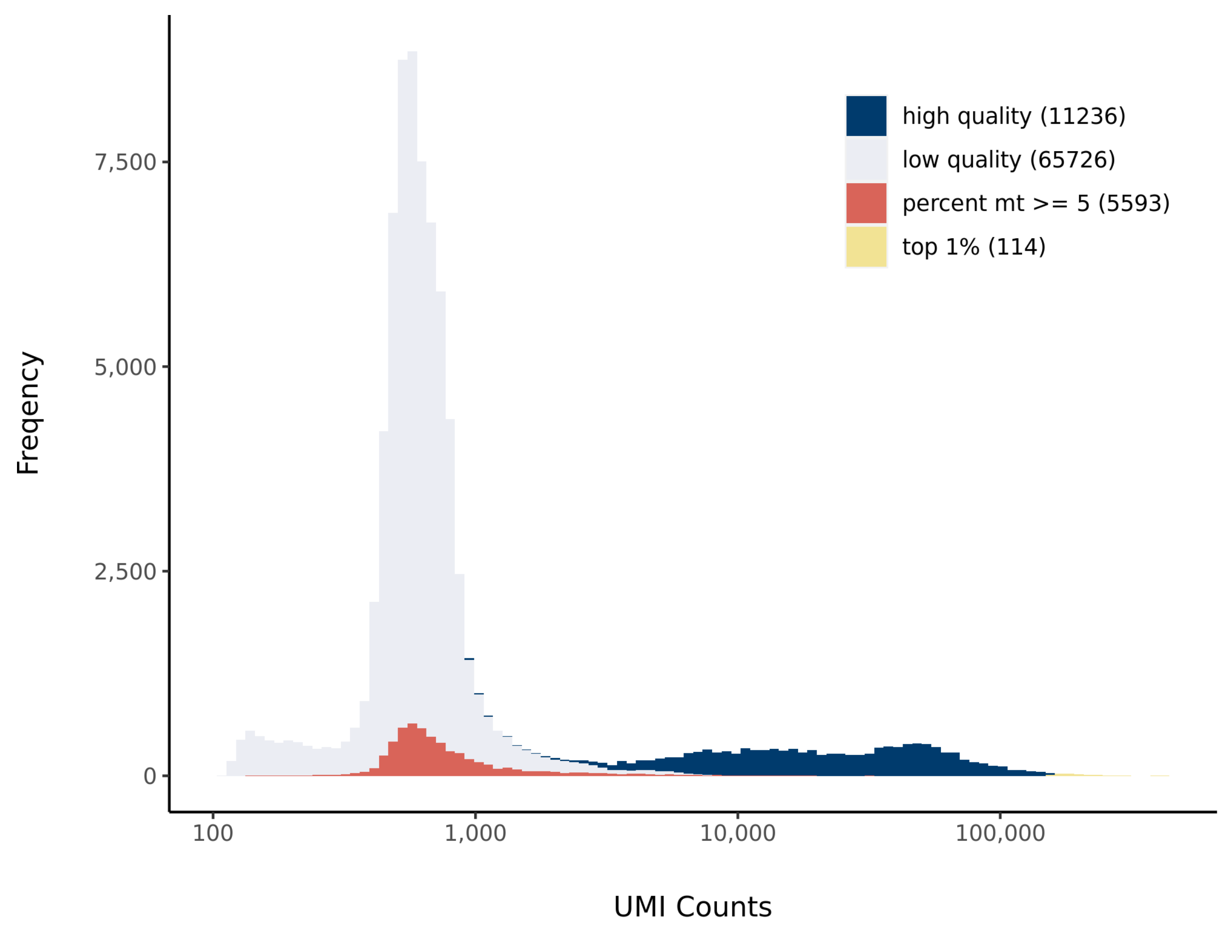


#### Number of Genes Histogram

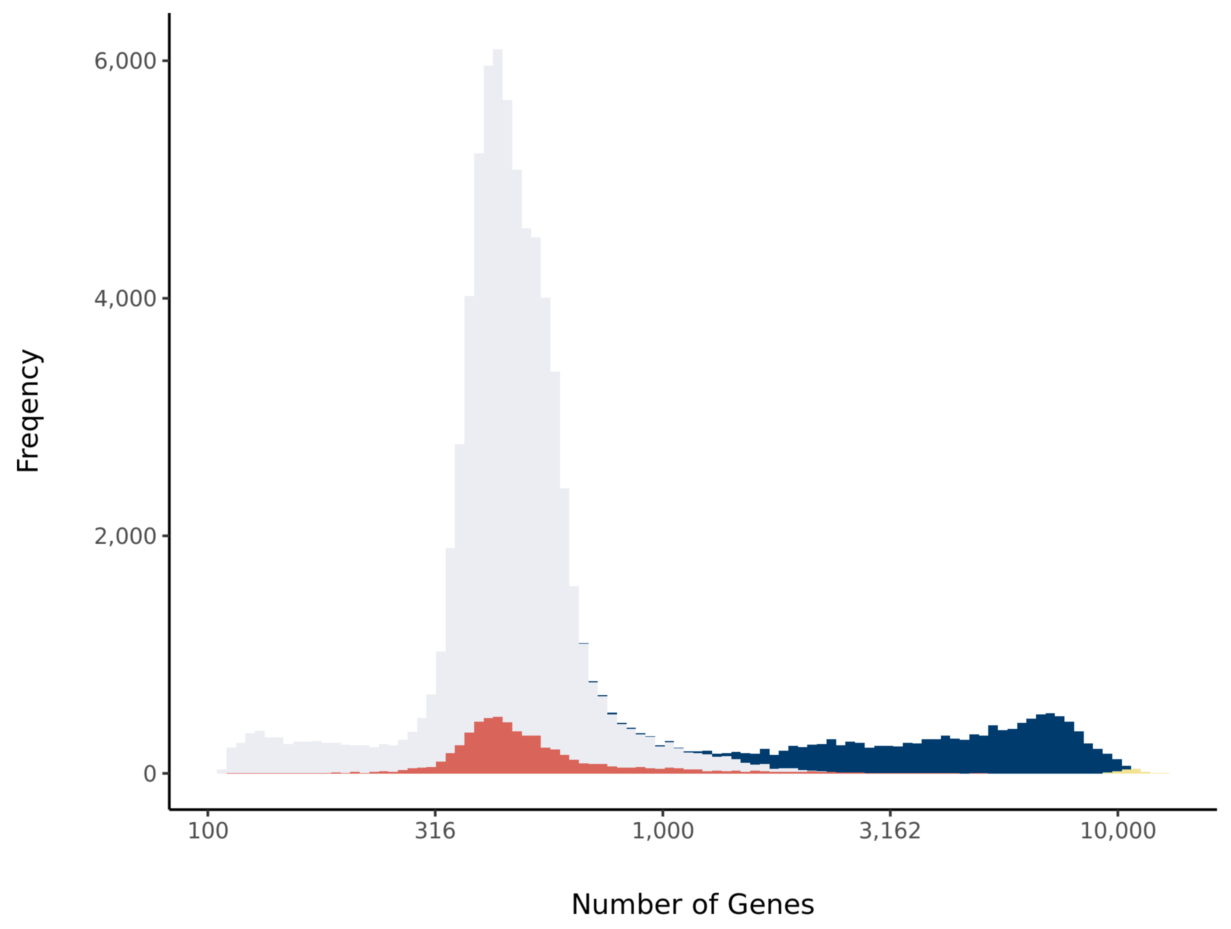


#### Barcode Rank Plot

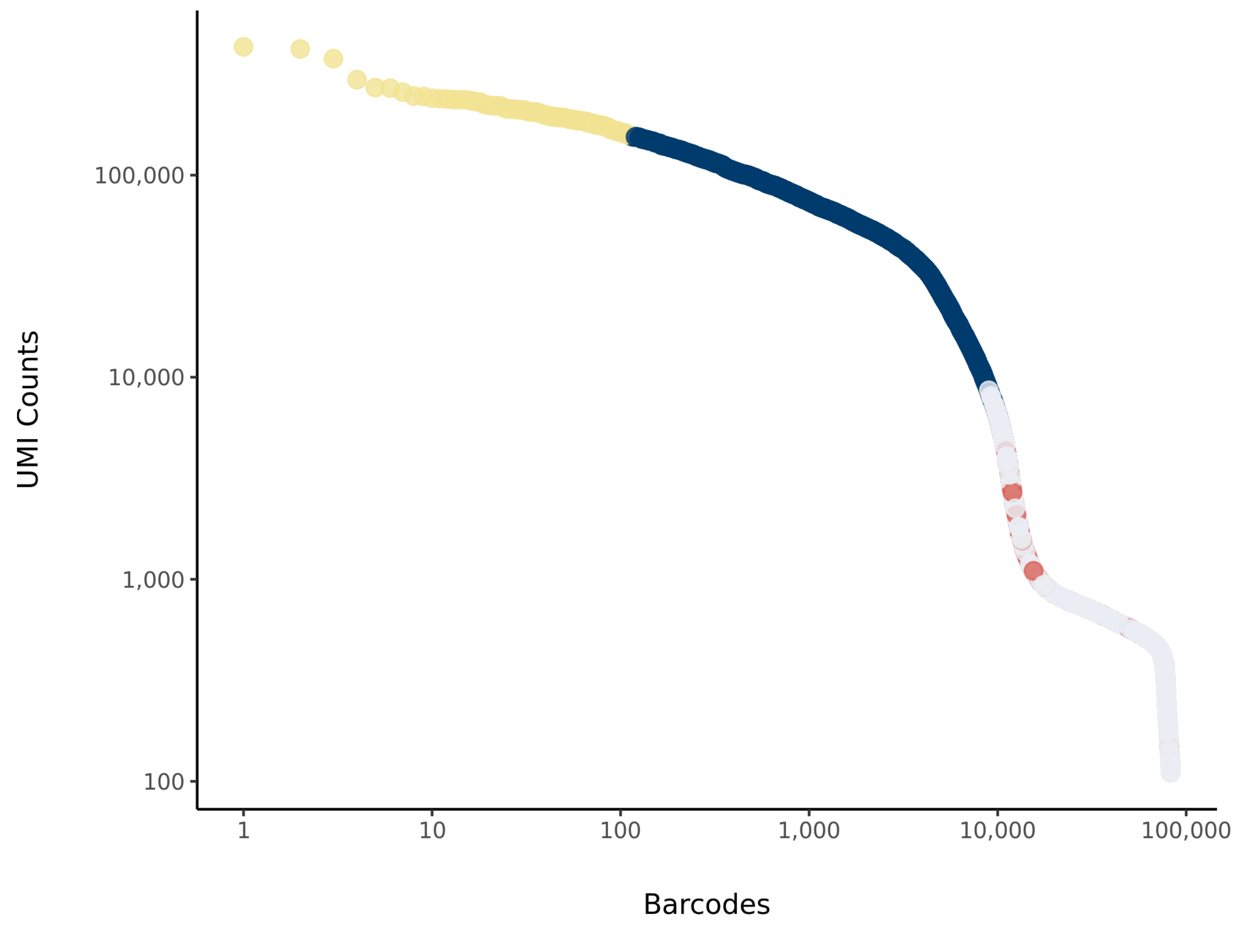

### sc_20 Summary

Processed by COPILOT

Summary Analysis

#### Parameters

| | |
|---|---|
| Iteration of Filtering | 1 |
| Mitochondrial Expression Threshold | 5 % |
| Top High Quality Cell Filtered | 1 % |
| Doublet Removed | Yes |

#### Cell Stats

| | |
|---|---|
| Estimated Number of High Quality Cell | 14,064 |
| High Quality Cell | 19.46 % |
| Total UMI Counts in High Quality Cell | 135,176,770 |
| UMI Counts in High Quality Cell | 54.36 % |
| Median UMI Counts per High Quality Cell | 6,497 |
| Median Genes per High Quality Cell | 2,711 |
| Total Genes Detected in High Quality Cell | 25,325 |
| Cell above Mitochondrial Expression Threshold | 0.93 % |
| Estimated Doublet Rate in High Quality Cell | 10.47 % |

#### Sequencing Stats

| | |
|---|---|
| Number of Reads Processed | 327,662,124 |
| Reads Pseudoaligned | 94.7 % |
| Reads on Whitelist | 96.89 % |
| Total UMI Counts | 248,691,560 |
| Sequencing Technology | 10xv3 |
| Species | Arabidopsis thaliana |
| Transcriptome | TAIR10 |

#### Sample Stats

| | |
|---|---|
| Sample | sc_20 |
| Name | WT Col-0_RS1 |
| Source | Benfey lab |
| Genotype | WT Col-0 |
| Transgene | NA |
| Treatment | Untreated |
| Age | 5_day |
| Timepoint | NA |
| Rep | NA |
| Target Cells | 10,000 |
| Date | NA |
| Seq Run | Shahan_6177 |

#### UMI Counts Histogram

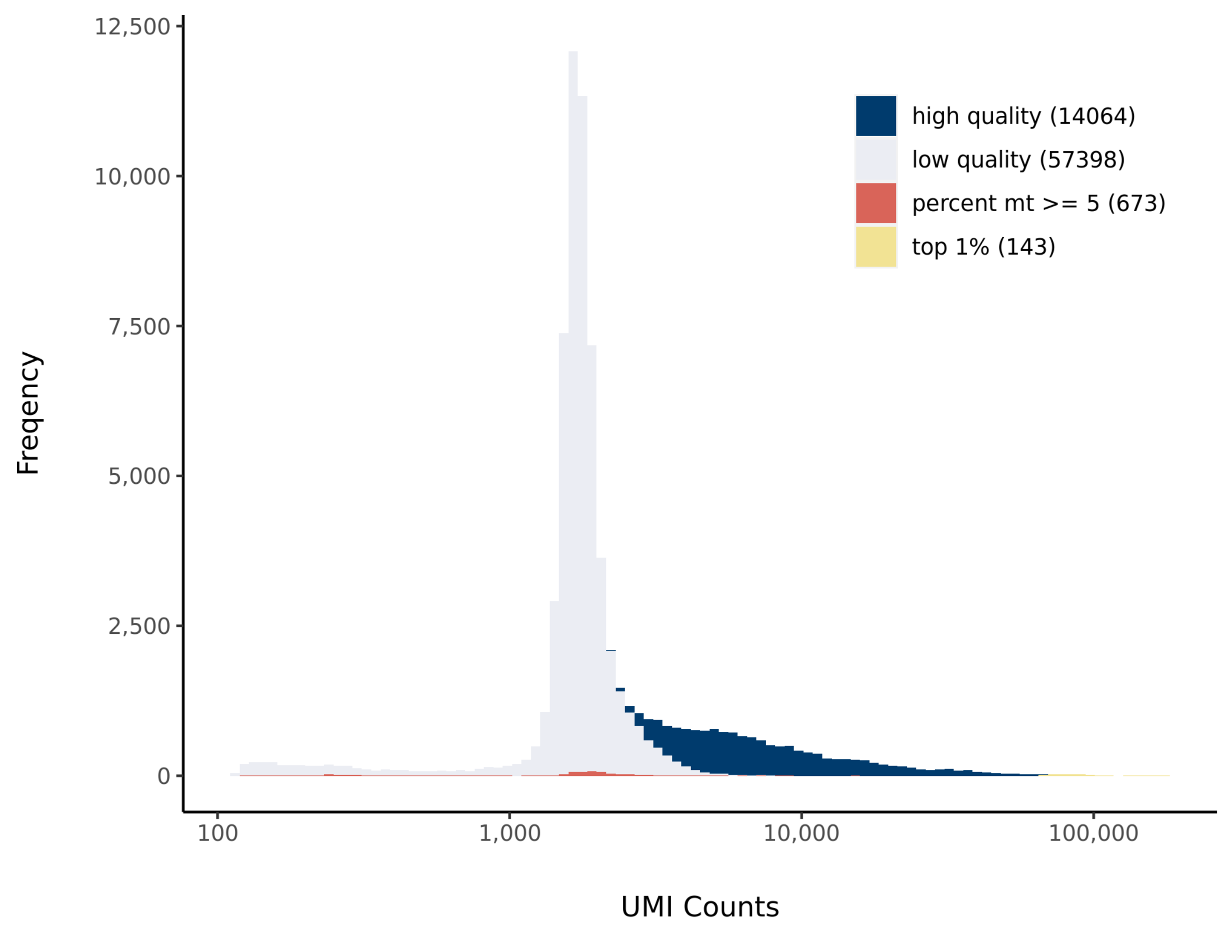


#### Number of Genes Histogram

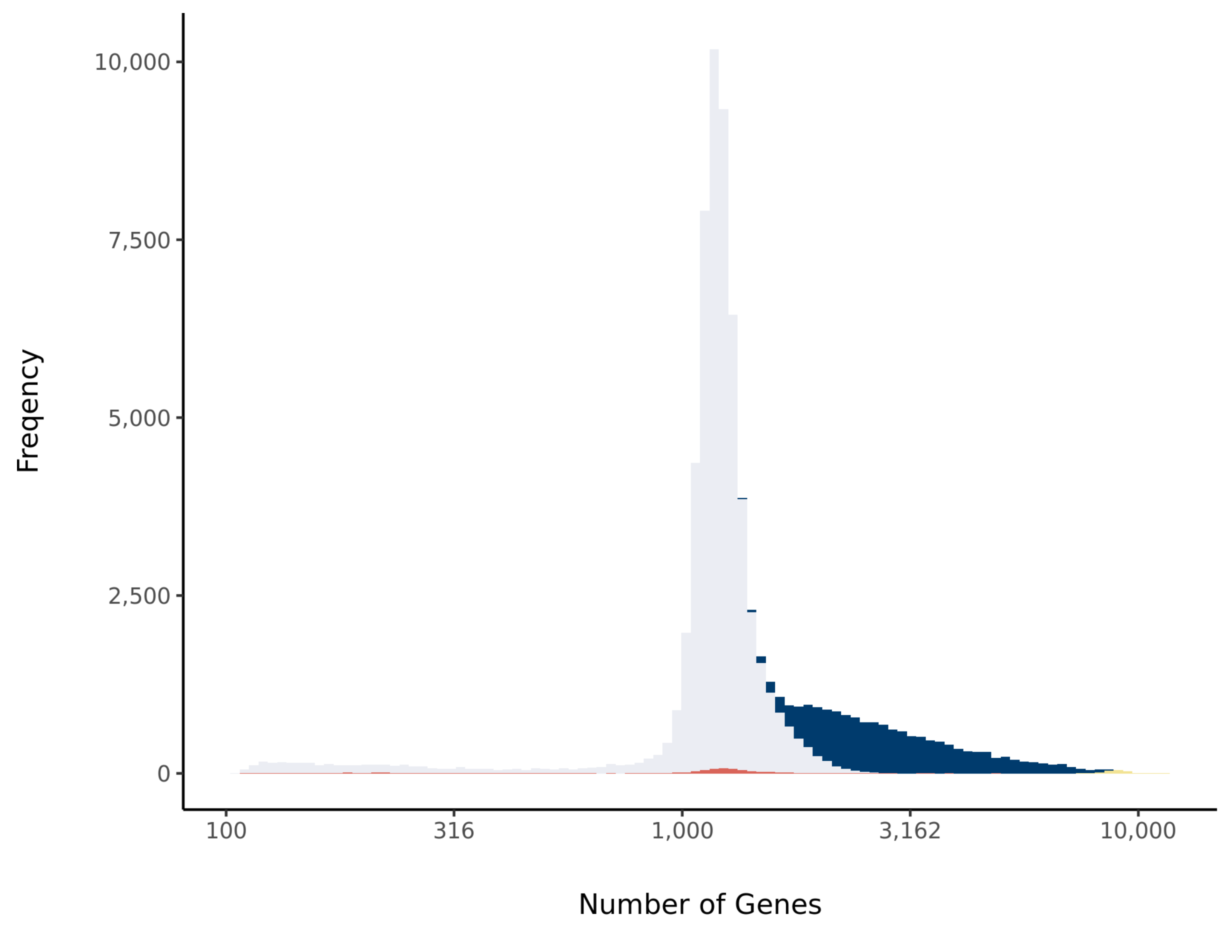


#### Barcode Rank Plot

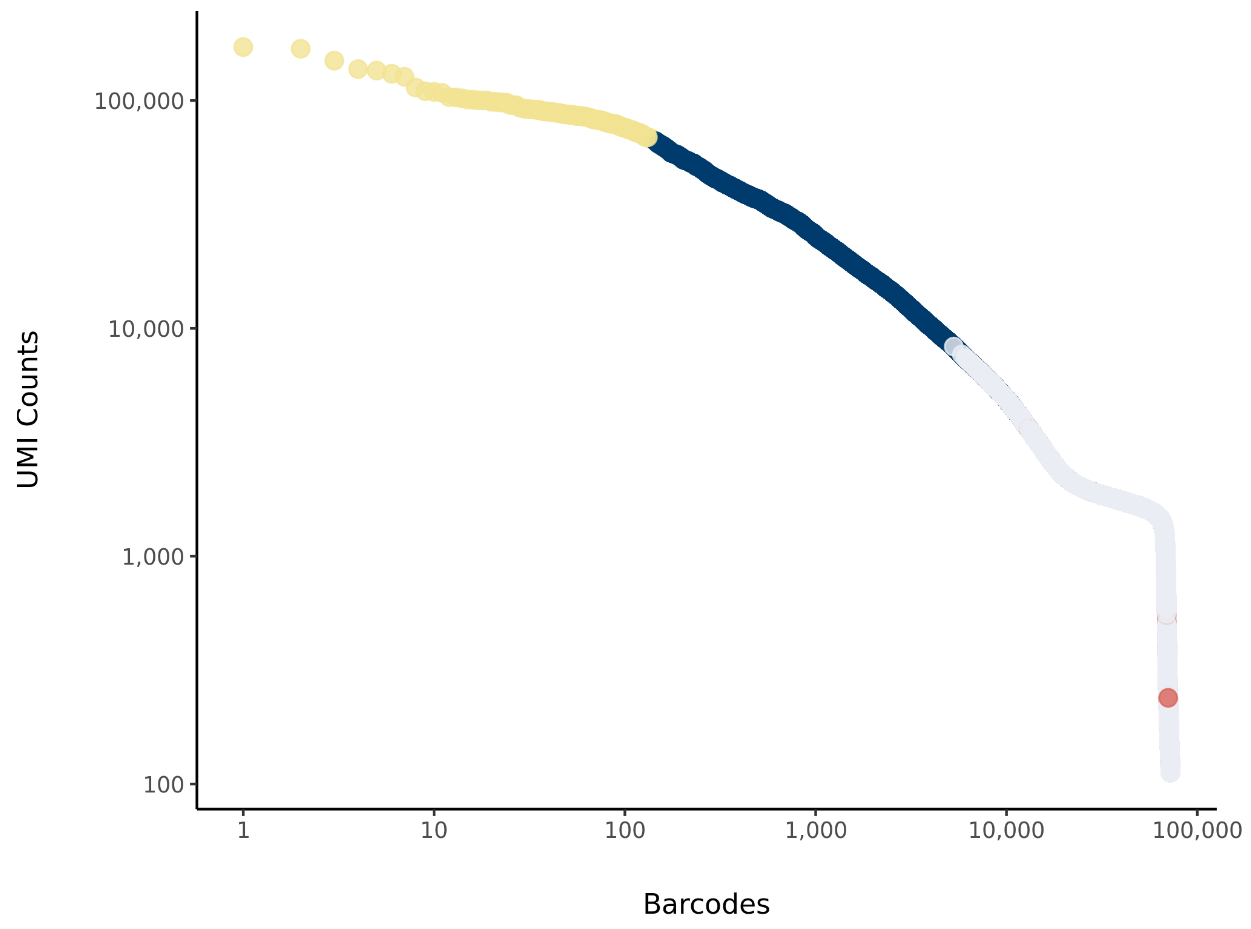

### sc_21 Summary

Processed by COPILOT

Summary Analysis

#### Parameters

| | |
|---|---|
| Iteration of Filtering | 1 |
| Mitochondrial Expression Threshold | 5 % |
| Top High Quality Cell Filtered | 1 % |
| Doublet Removed | Yes |

#### Cell Stats

| | |
|---|---|
| Estimated Number of High Quality Cell | 10,478 |
| High Quality Cell | 14.65 % |
| Total UMI Counts in High Quality Cell | 128,289,748 |
| UMI Counts in High Quality Cell | 53.13 % |
| Median UMI Counts per High Quality Cell | 7,833.5 |
| Median Genes per High Quality Cell | 2,899 |
| Total Genes Detected in High Quality Cell | 25,296 |
| Cell above Mitochondrial Expression Threshold | 0.53 % |
| Estimated Doublet Rate in High Quality Cell | 7.83 % |

#### Sequencing Stats

| | |
|---|---|
| Number of Reads Processed | 318,494,791 |
| Reads Pseudoaligned | 95.2 % |
| Reads on Whitelist | 96.95 % |
| Total UMI Counts | 241,458,602 |
| Sequencing Technology | 10xv3 |
| Species | Arabidopsis thaliana |
| Transcriptome | TAIR10 |

#### Sample Stats

| | |
|---|---|
| Sample | sc_21 |
| Name | WT Col-0_RS2 |
| Source | Benfey lab |
| Genotype | WT Col-0 |
| Transgene | NA |
| Treatment | Untreated |
| Age | 5_day |
| Timepoint | NA |
| Rep | NA |
| Target Cells | 10,000 |
| Date | NA |
| Seq Run | Shahan_6177 |

#### UMI Counts Histogram

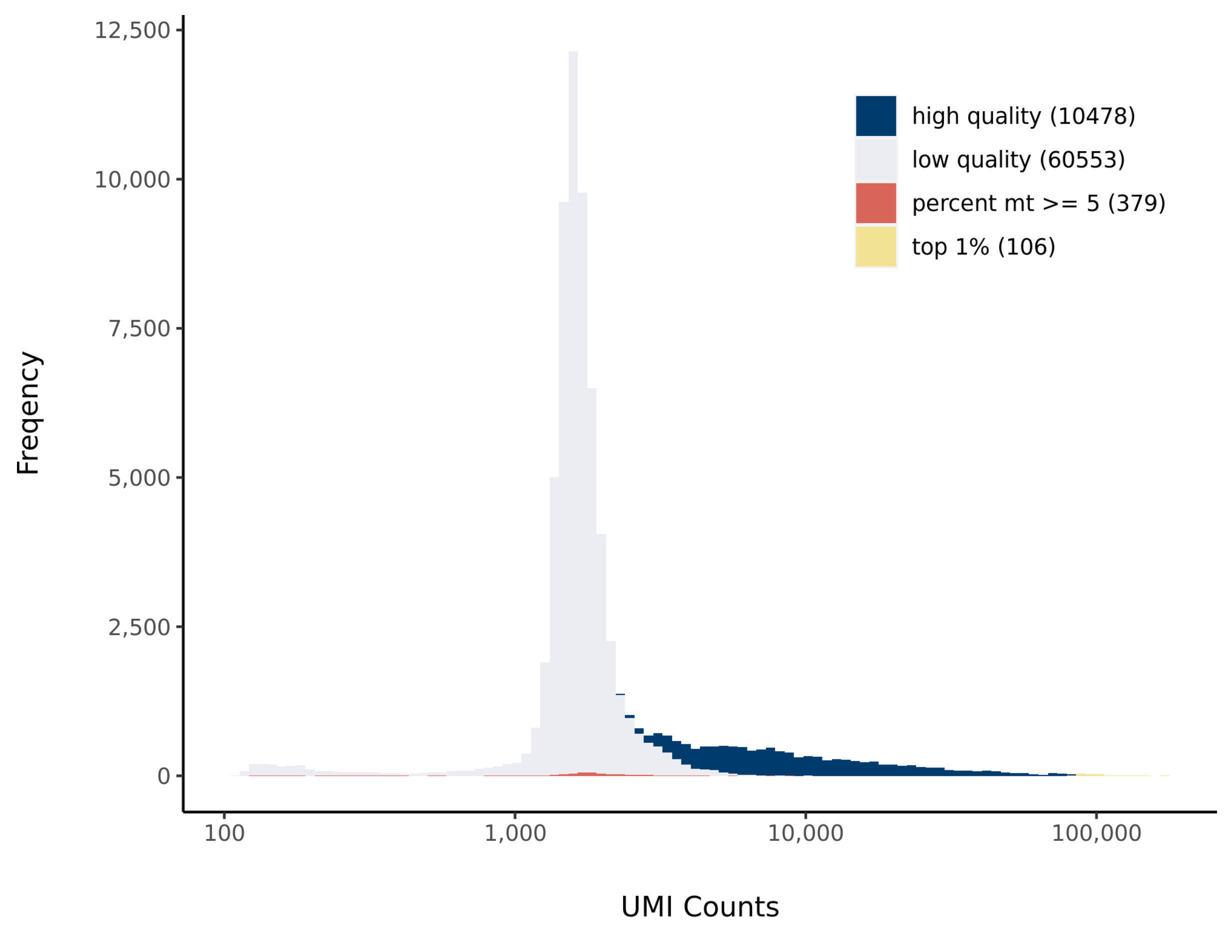


#### Number of Genes Histogram

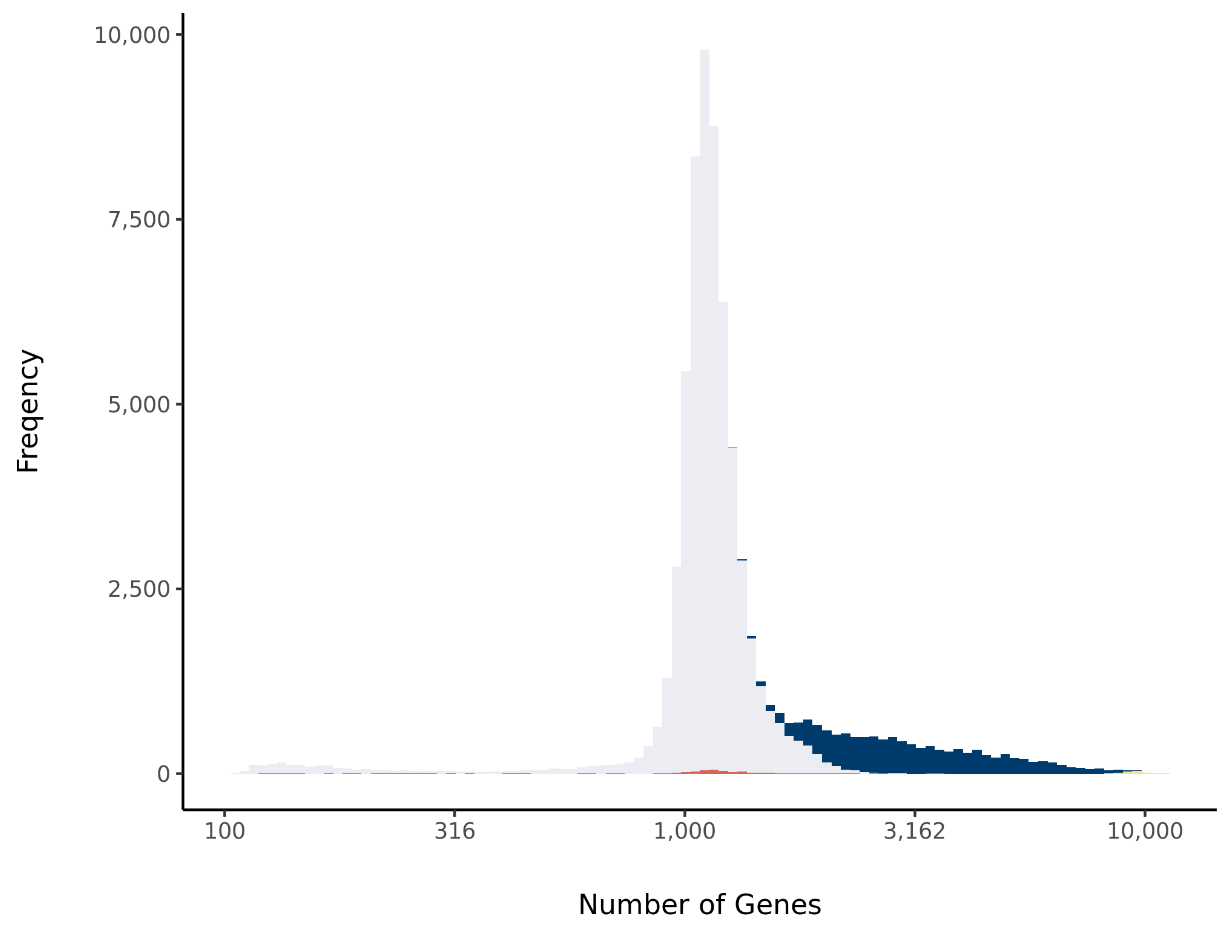


#### Barcode Rank Plot

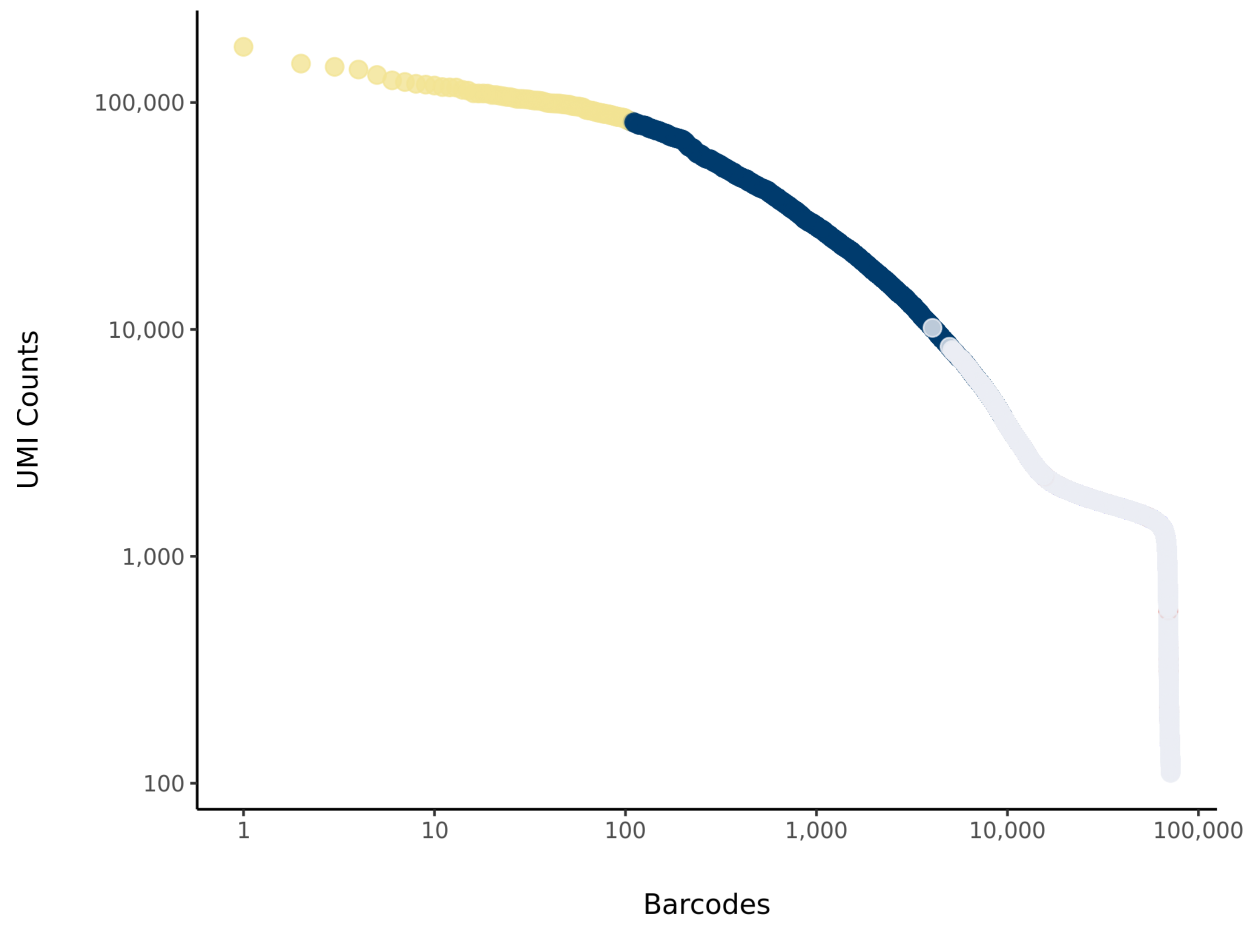

### sc_25 Summary

Processed by COPILOT

Summary Analysis

#### Parameters

| | |
|---|---|
| Iteration of Filtering | 1 |
| Mitochondrial Expression Threshold | 5 % |
| Top High Quality Cell Filtered | 1 % |
| Doublet Removed | Yes |

#### Cell Stats

| | |
|---|---|
| Estimated Number of High Quality Cell | 8,246 |
| High Quality Cell | 12.34 % |
| Total UMI Counts in High Quality Cell | 151,165,418 |
| UMI Counts in High Quality Cell | 59.19 % |
| Median UMI Counts per High Quality Cell | 11,167 |
| Median Genes per High Quality Cell | 3,465 |
| Total Genes Detected in High Quality Cell | 25,744 |
| Cell above Mitochondrial Expression Threshold | 1.47 % |
| Estimated Doublet Rate in High Quality Cell | 6.19 % |

#### Sequencing Stats

| | |
|---|---|
| Number of Reads Processed | 357,488,578 |
| Reads Pseudoaligned | 94.6 % |
| Reads on Whitelist | 97.17 % |
| Total UMI Counts | 255,403,407 |
| Sequencing Technology | 10xv3 |
| Species | Arabidopsis thaliana |
| Transcriptome | TAIR10 |

#### Sample Stats

| | |
|---|---|
| Sample | sc_25 |
| Name | scr-4_1 |
| Source | Benfey lab |
| Genotype | scr-4 (Ws backcrossed to Col?) |
| Transgene | NA |
| Treatment | Untreated |
| Age | 5_day |
| Timepoint | NA |
| Rep | NA |
| Target Cells | 10,000 |
| Date | NA |
| Seq Run | Shahan_6177 |

#### UMI Counts Histogram

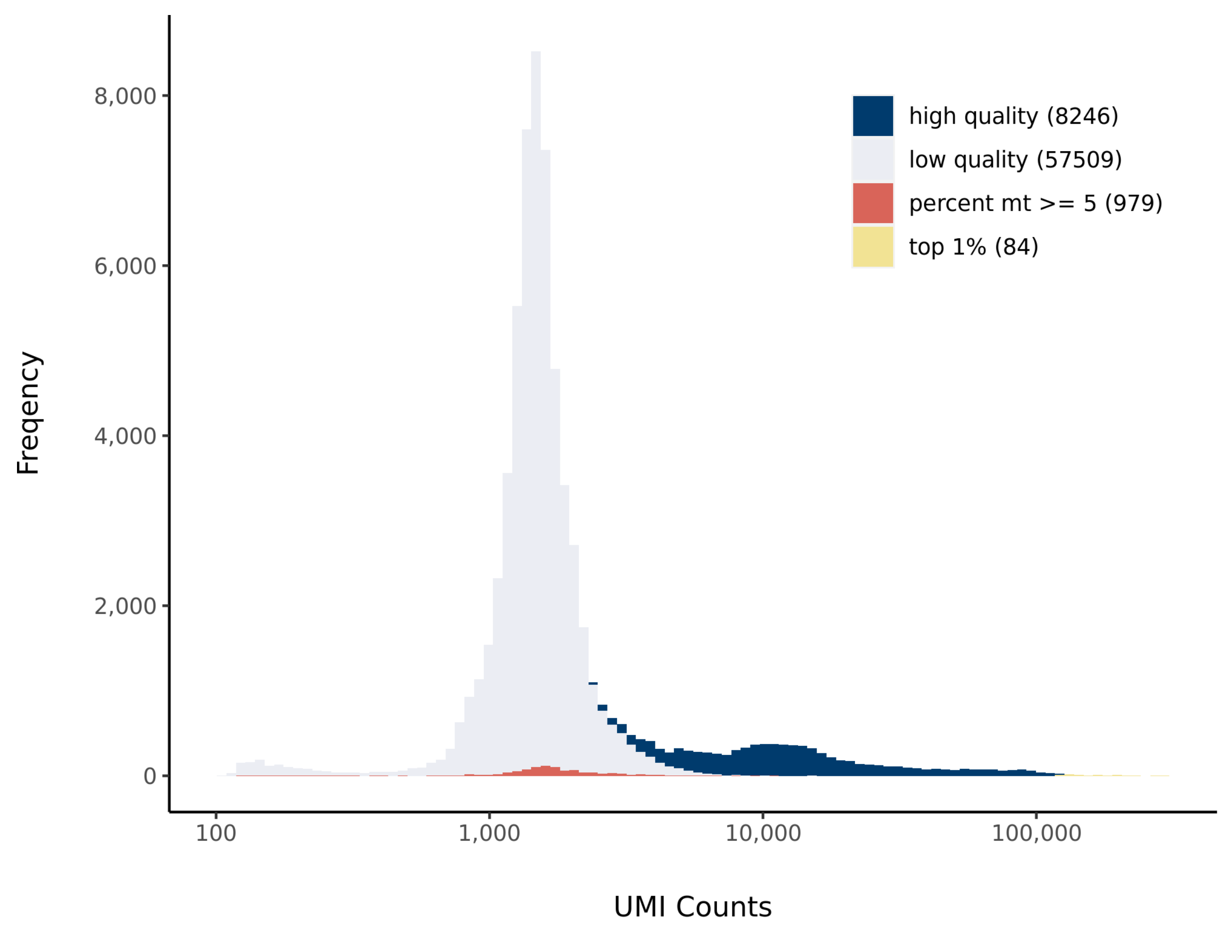


#### Number of Genes Histogram

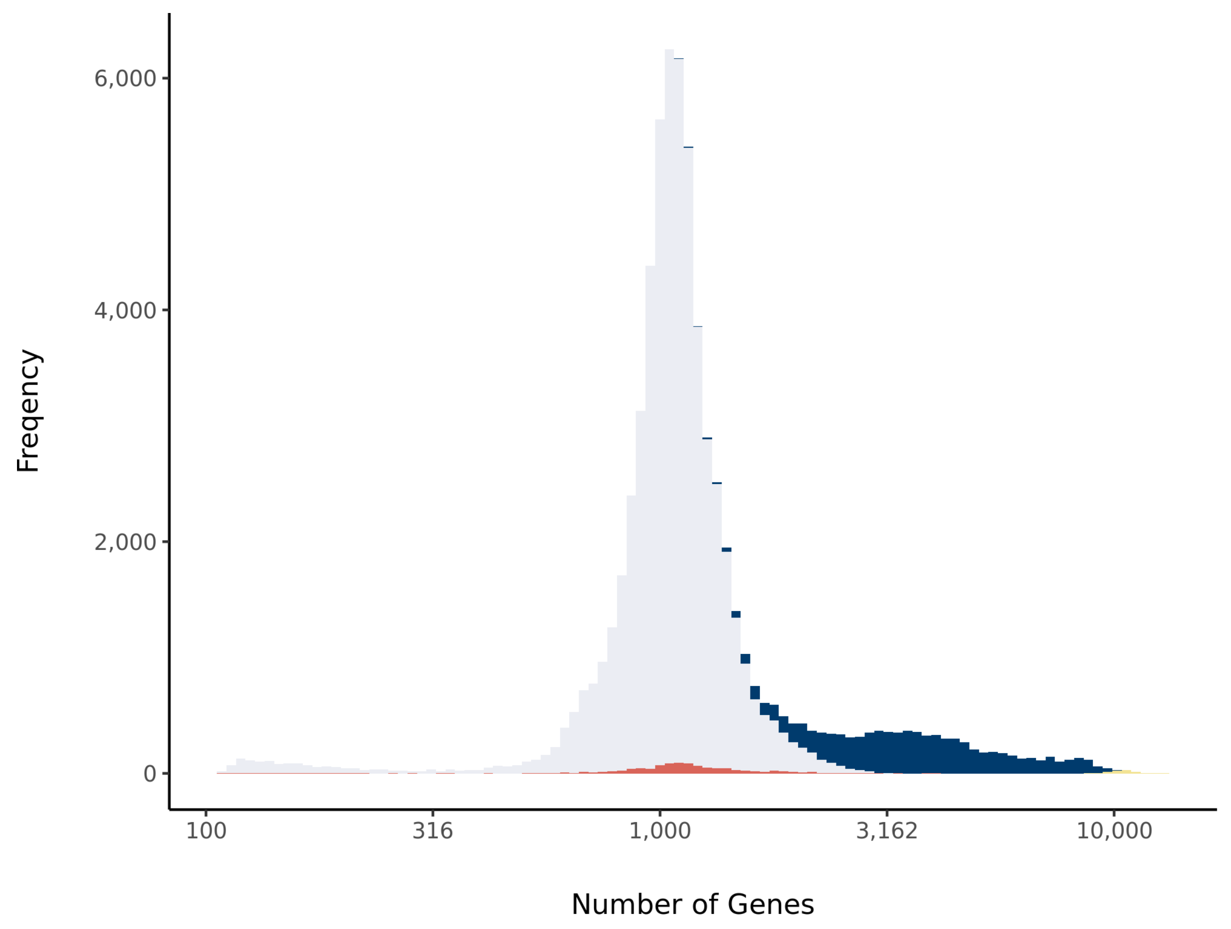


#### Barcode Rank Plot

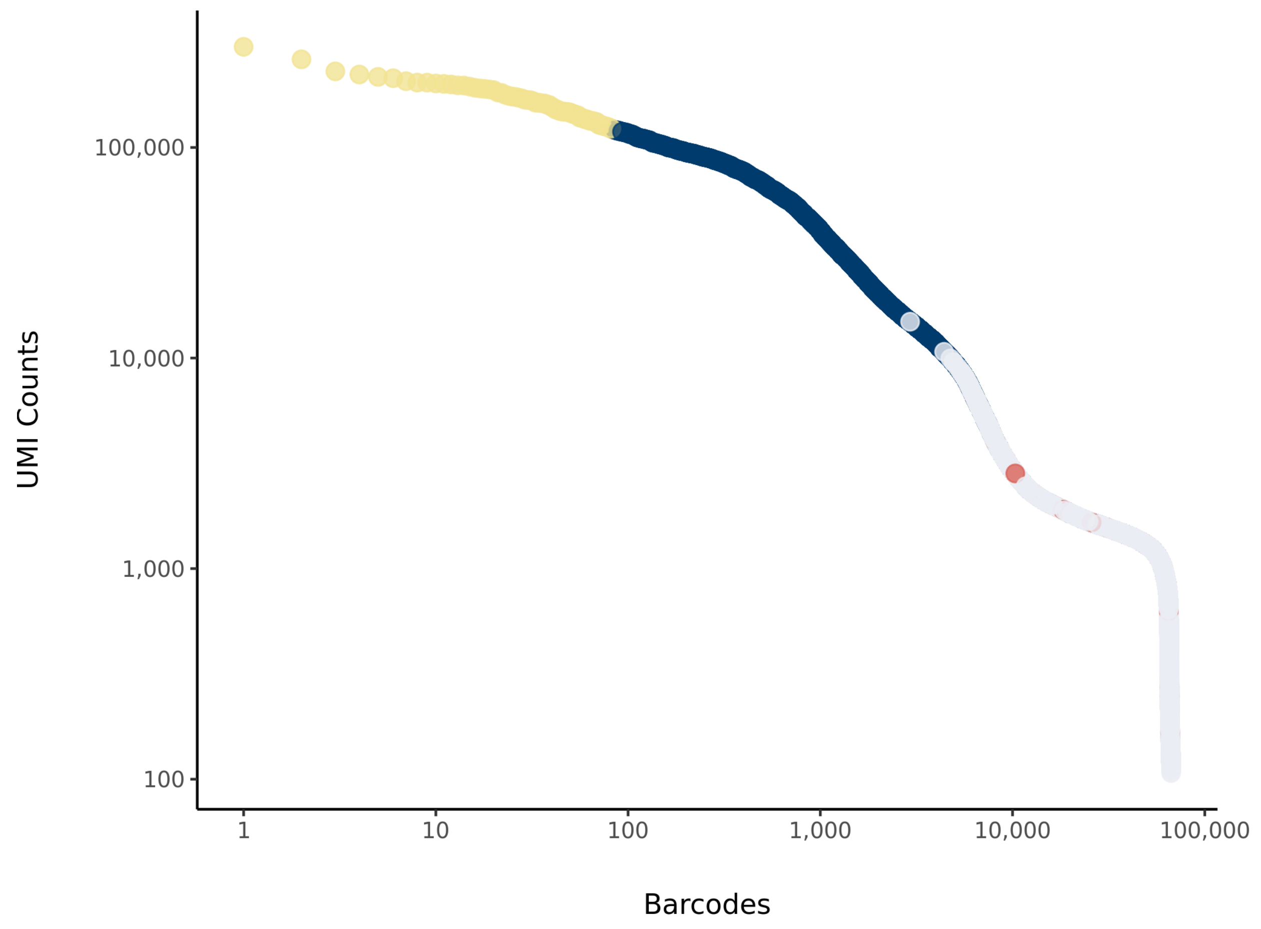

### sc_30 Summary

Processed by COPILOT



#### Parameters

| | |
|---|---|
| Iteration of Filtering | 1 |
| Mitochondrial Expression Threshold | 5 % |
| Top High Quality Cell Filtered | 1 % |
| Doublet Removed | Yes |

#### Cell Stats

| | |
|---|---|
| Estimated Number of High Quality Cell | 11,118 |
| High Quality Cell | 14.83 % |
| Total UMI Counts in High Quality Cell | 137,632,134 |
| UMI Counts in High Quality Cell | 79.01 % |
| Median UMI Counts per High Quality Cell | 6,495 |
| Median Genes per High Quality Cell | 2,466.5 |
| Total Genes Detected in High Quality Cell | 24,739 |
| Cell above Mitochondrial Expression Threshold | 4.71 % |
| Estimated Doublet Rate in High Quality Cell | 8.3 % |

#### Sequencing Stats

| | |
|---|---|
| Number of Reads Processed | 263,408,174 |
| Reads Pseudoaligned | 93.5 % |
| Reads on Whitelist | 96.22 % |
| Total UMI Counts | 174,187,605 |
| Sequencing Technology | 10xv3 |
| Species | Arabidopsis thaliana |
| Transcriptome | TAIR10 |

#### Sample Stats

| | |
|---|---|
| Sample | sc_30 |
| Name | WT Col-0_RS3 |
| Source | Benfey lab |
| Genotype | WT Col-0 |
| Transgene | NA |
| Treatment | Untreated |
| Age | 5_day |
| Timepoint | NA |
| Rep | NA |
| Target Cells | 10,000 |
| Date | NA |
| Seq Run | Nolan_6199 (NextSeq); Nolan_6226 (NovaSeq S4) |

#### UMI Counts Histogram

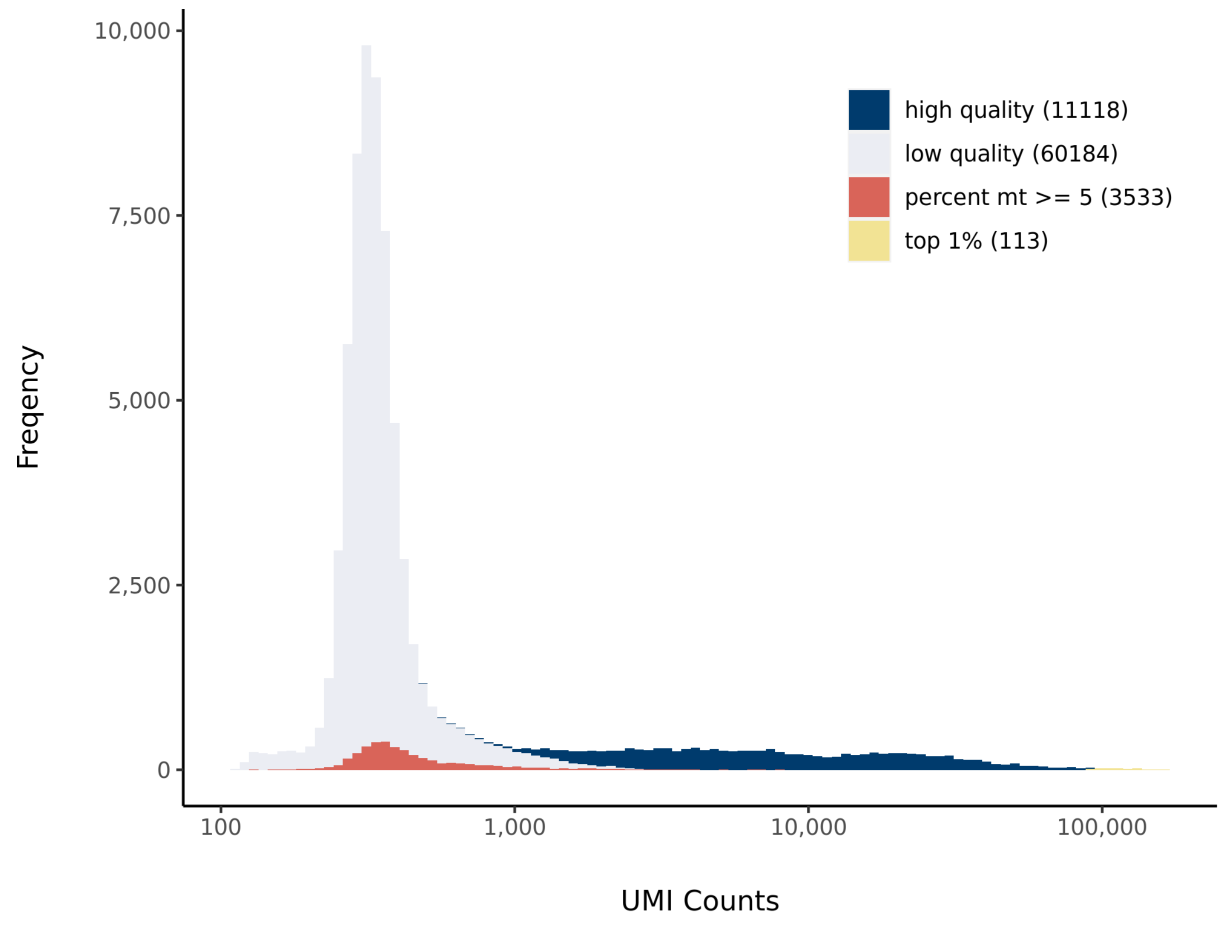


#### Number of Genes Histogram

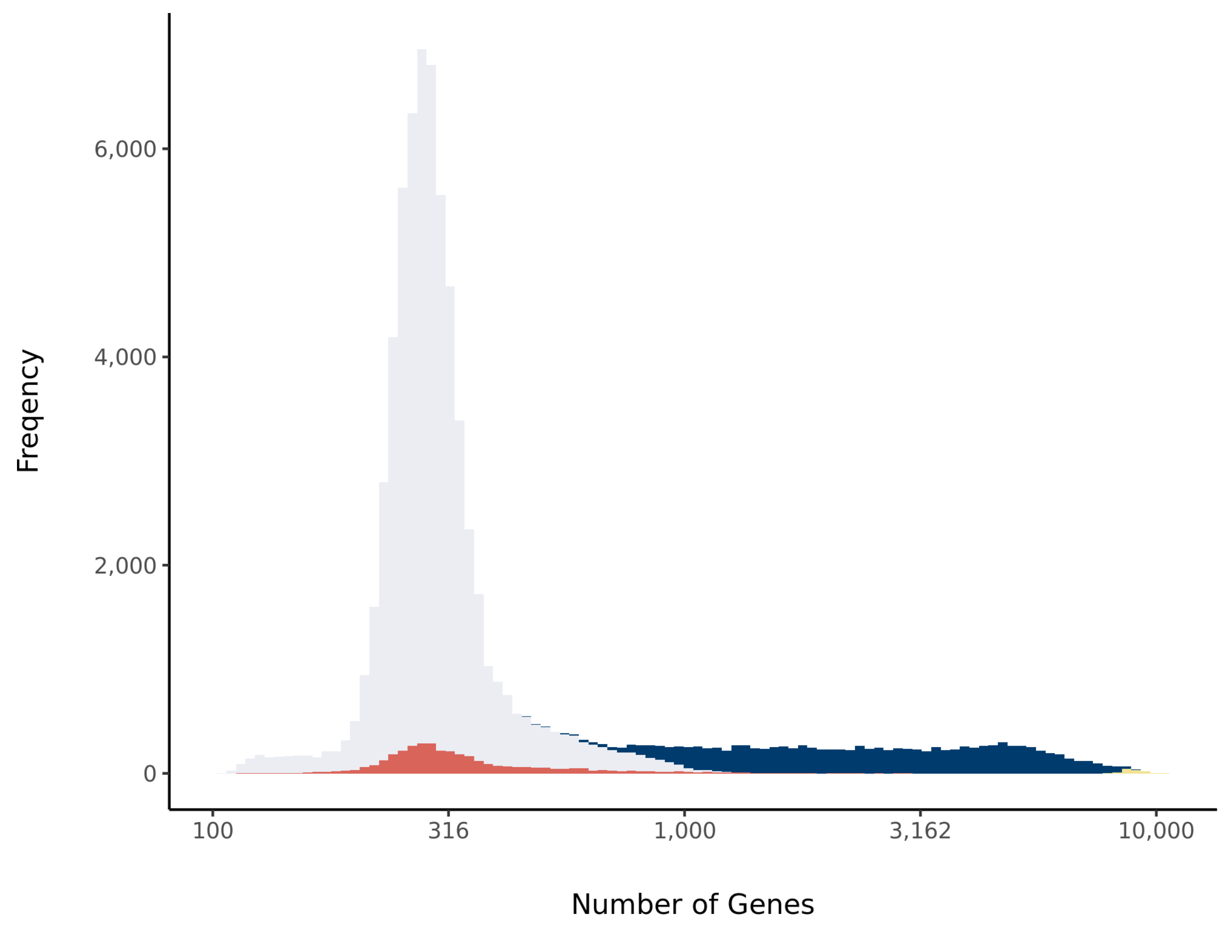


#### Barcode Rank Plot

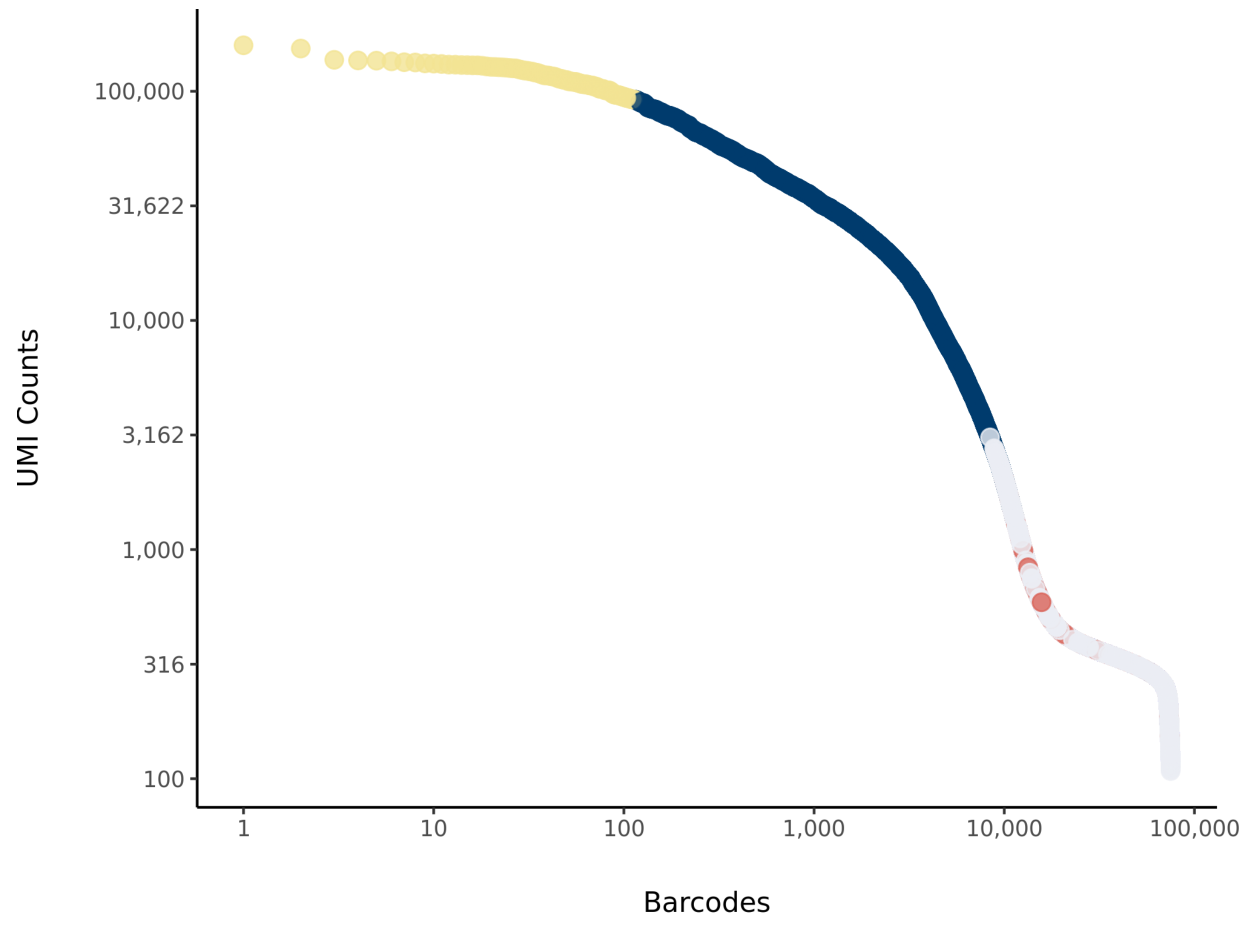

### sc_31 Summary

Processed by COPILOT

Summary | Analysis

#### Parameters

| | |
|---|---|
| Iteration of Filtering | 1 |
| Mitochondrial Expression Threshold | 5 % |
| Top High Quality Cell Filtered | 1 % |
| Doublet Removed | Yes |

#### Cell Stats

| | |
|---|---|
| Estimated Number of High Quality Cell | 9,443 |
| High Quality Cell | 12.54 % |
| Total UMI Counts in High Quality Cell | 128,218,174 |
| UMI Counts in High Quality Cell | 76.36 % |
| Median UMI Counts per High Quality Cell | 6,034 |
| Median Genes per High Quality Cell | 2,308 |
| Total Genes Detected in High Quality Cell | 24,826 |
| Cell above Mitochondrial Expression Threshold | 16.13 % |
| Estimated Doublet Rate in High Quality Cell | 7.07 % |

#### Sequencing Stats

| | |
|---|---|
| Number of Reads Processed | 266,362,743 |
| Reads Pseudoaligned | 92.5 % |
| Reads on Whitelist | 95.91 % |
| Total UMI Counts | 167,906,304 |
| Sequencing Technology | 10xv3 |
| Species | Arabidopsis thaliana |
| Transcriptome | TAIR10 |

#### Sample Stats

| | |
|---|---|
| Sample | sc_31 |
| Name | WT Col-0_RS4 |
| Source | Benfey lab |
| Genotype | WT Col-0 |
| Transgene | NA |
| Treatment | Untreated |
| Age | 5_day |
| Timepoint | NA |
| Rep | NA |
| Target Cells | 10,000 |
| Date | NA |
| Seq Run | Nolan_6199 (NextSeq); Nolan_6226 (NovaSeq S4) |

#### UMI Counts Histogram

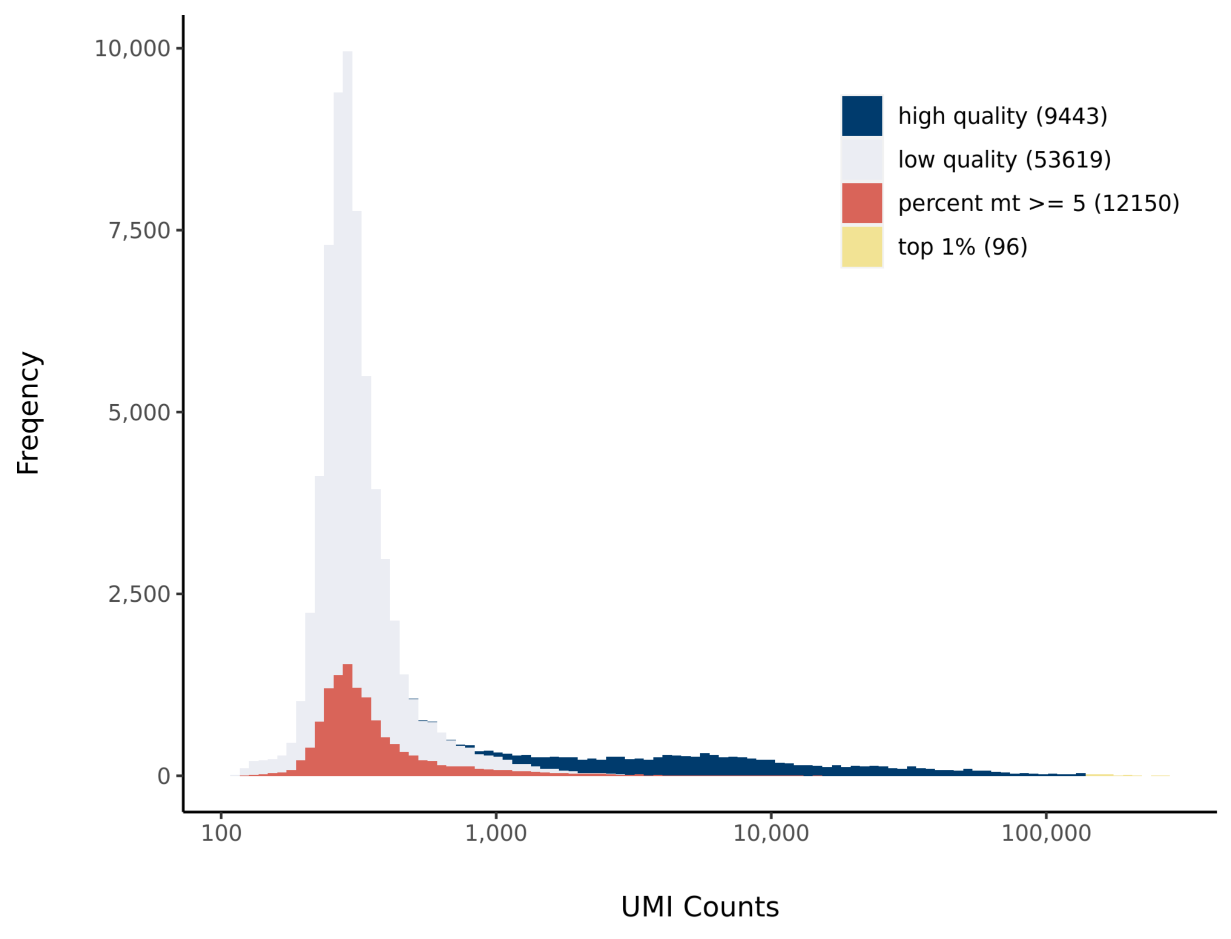


#### Number of Genes Histogram

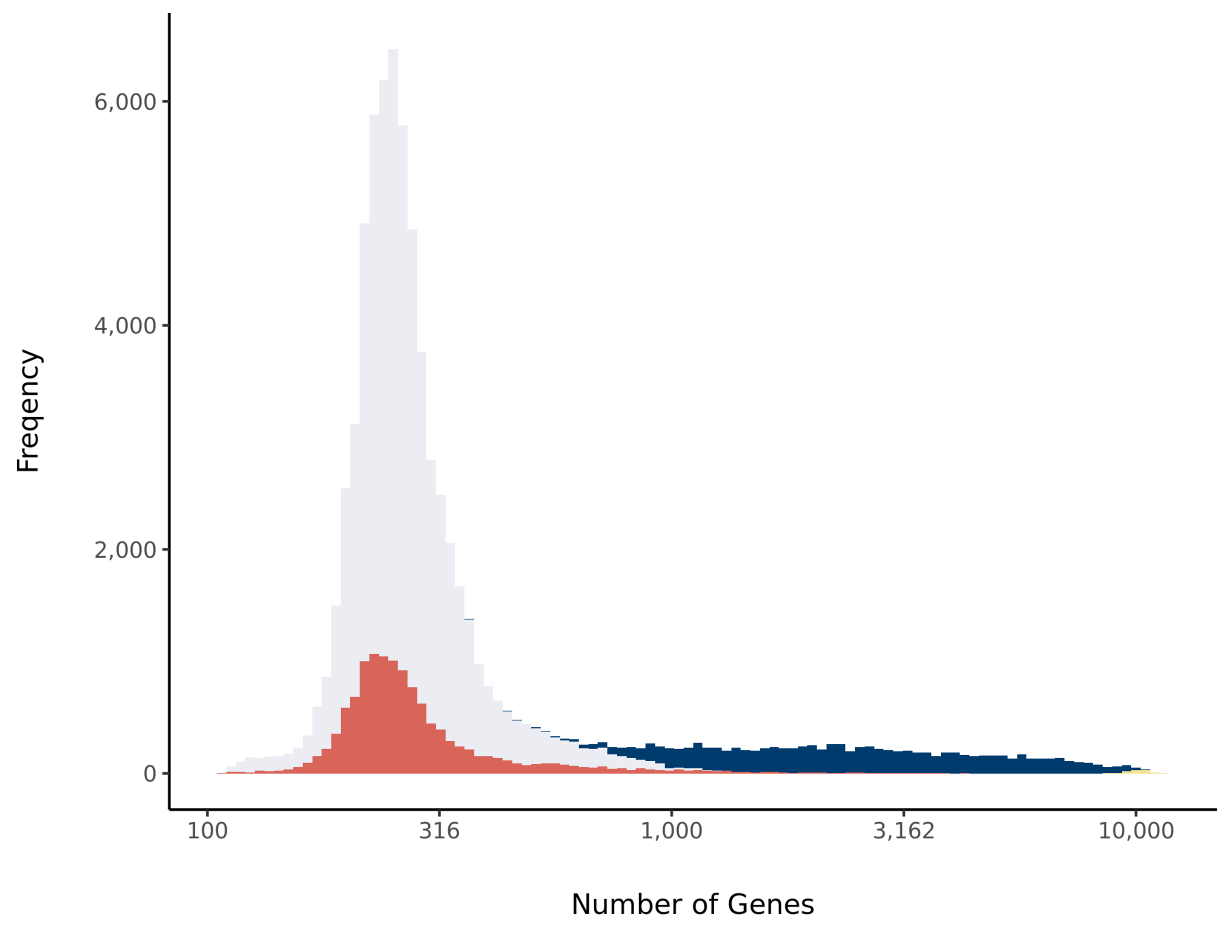


#### Barcode Rank Plot

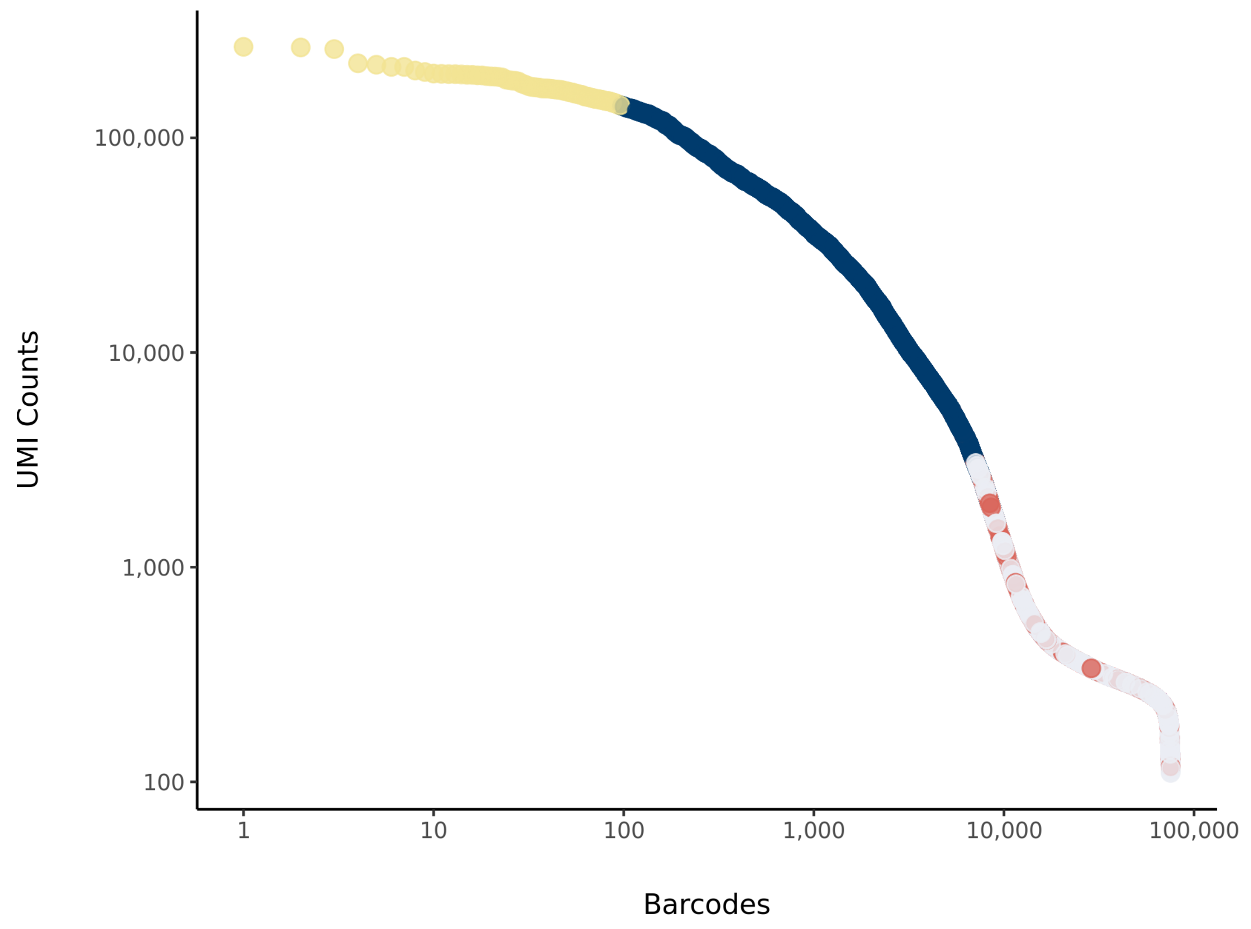

### sc_36 Summary

Processed by COPILOT

Summary Analysis

#### Parameters

| | |
|---|---|
| Iteration of Filtering | 1 |
| Mitochondrial Expression Threshold | 5 % |
| Top High Quality Cell Filtered | 1 % |
| Doublet Removed | Yes |

#### Cell Stats

| | |
|---|---|
| Estimated Number of High Quality Cell | 6,306 |
| High Quality Cell | 7.84 % |
| Total UMI Counts in High Quality Cell | 114,638,530 |
| UMI Counts in High Quality Cell | 76.67 % |
| Median UMI Counts per High Quality Cell | 9,849 |
| Median Genes per High Quality Cell | 3,209 |
| Total Genes Detected in High Quality Cell | 25,147 |
| Cell above Mitochondrial Expression Threshold | 33.79 % |
| Estimated Doublet Rate in High Quality Cell | 4.76 % |

#### Sequencing Stats

| | |
|---|---|
| Number of Reads Processed | 280,421,300 |
| Reads Pseudoaligned | 91.3 % |
| Reads on Whitelist | 95.85 % |
| Total UMI Counts | 149,515,088 |
| Sequencing Technology | 10xv3 |
| Species | Arabidopsis thaliana |
| Transcriptome | TAIR10 |

#### Sample Stats

| | |
|---|---|
| Sample | sc_36 |
| Name | scr_4_2 |
| Source | Benfey lab |
| Genotype | scr-4 |
| Transgene | NA |
| Treatment | Untreated |
| Age | 5_day |
| Timepoint | NA |
| Rep | NA |
| Target Cells | 10,000 |
| Date | NA |
| Seq Run | Nolan_6199 (NextSeq); Nolan_6226 (NovaSeq S4) |

#### UMI Counts Histogram

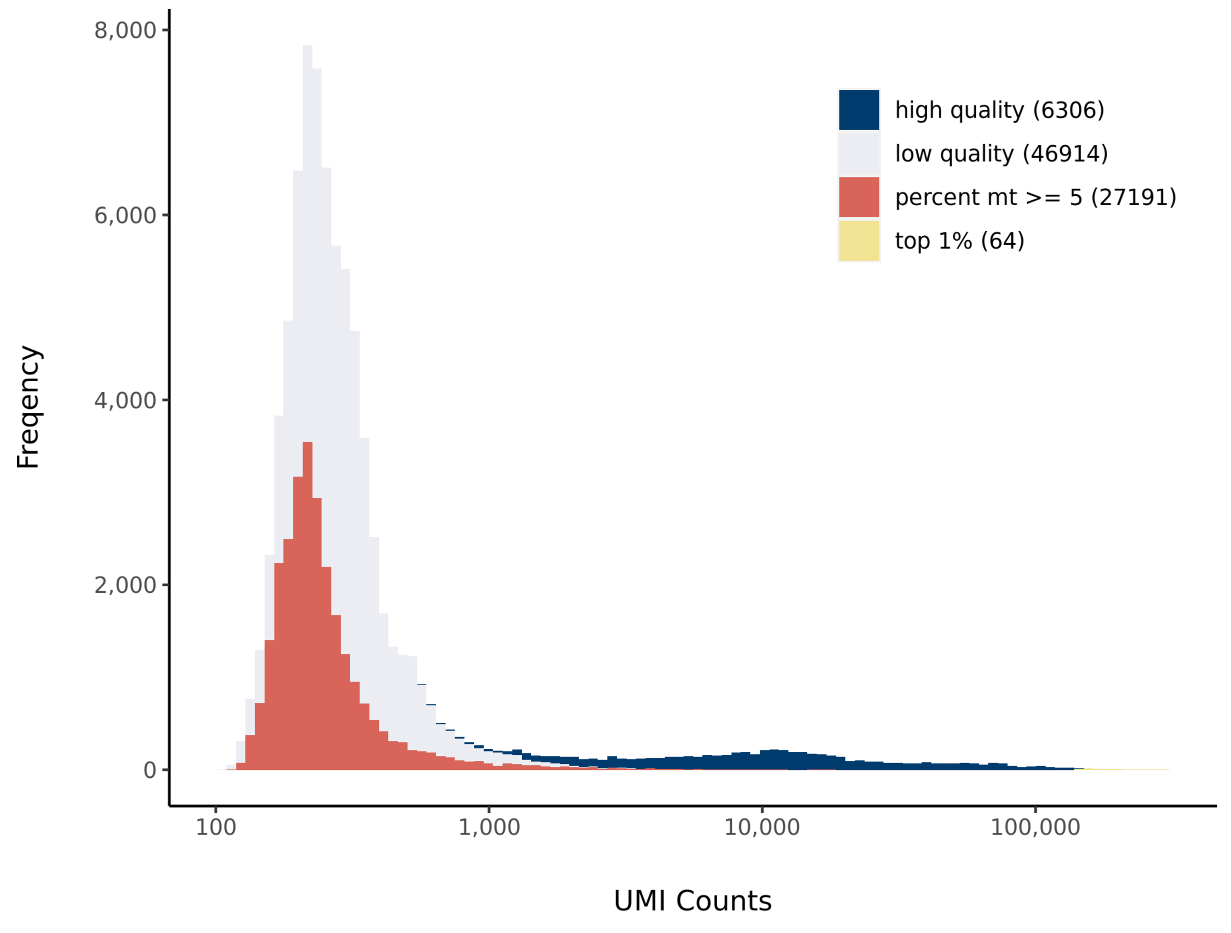


#### Number of Genes Histogram

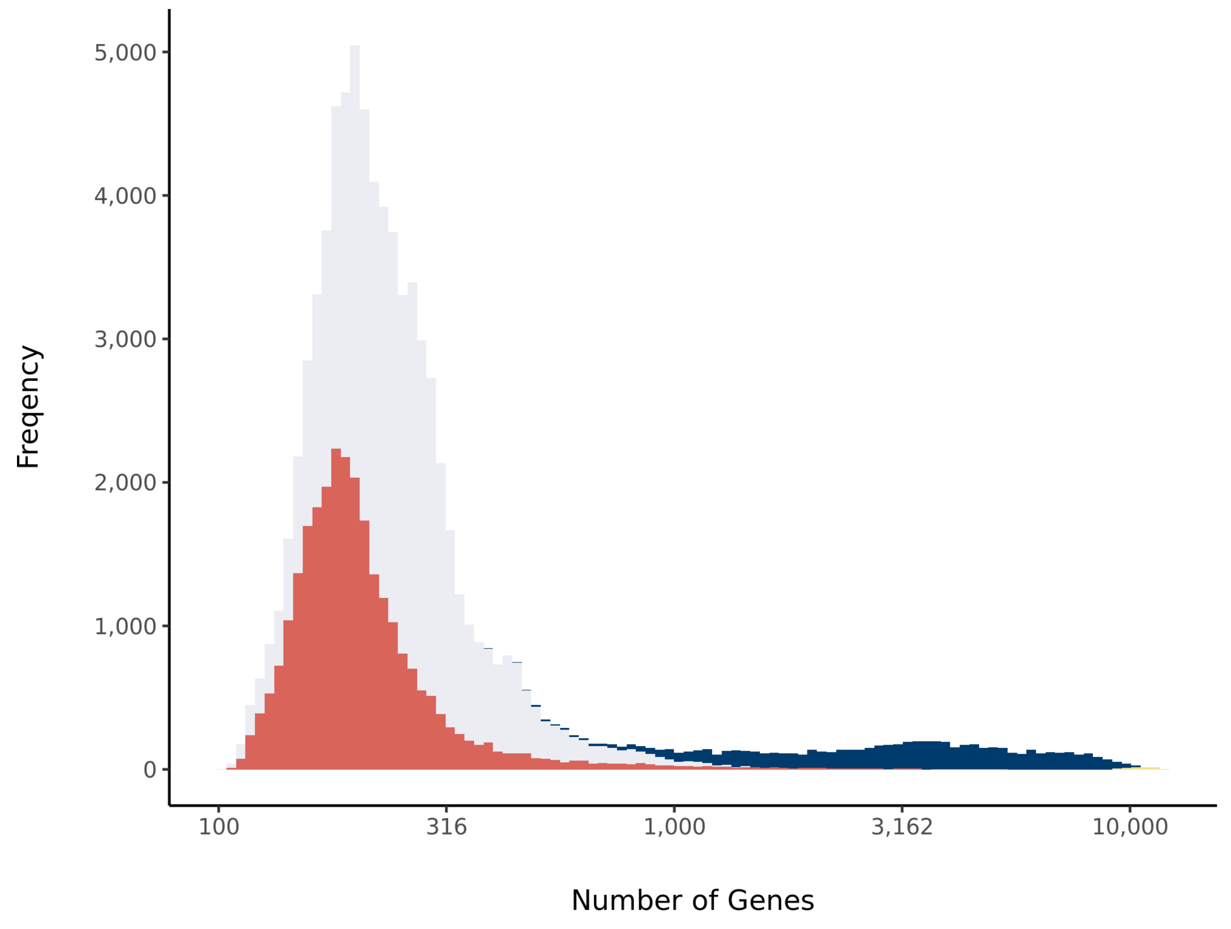


#### Barcode Rank Plot

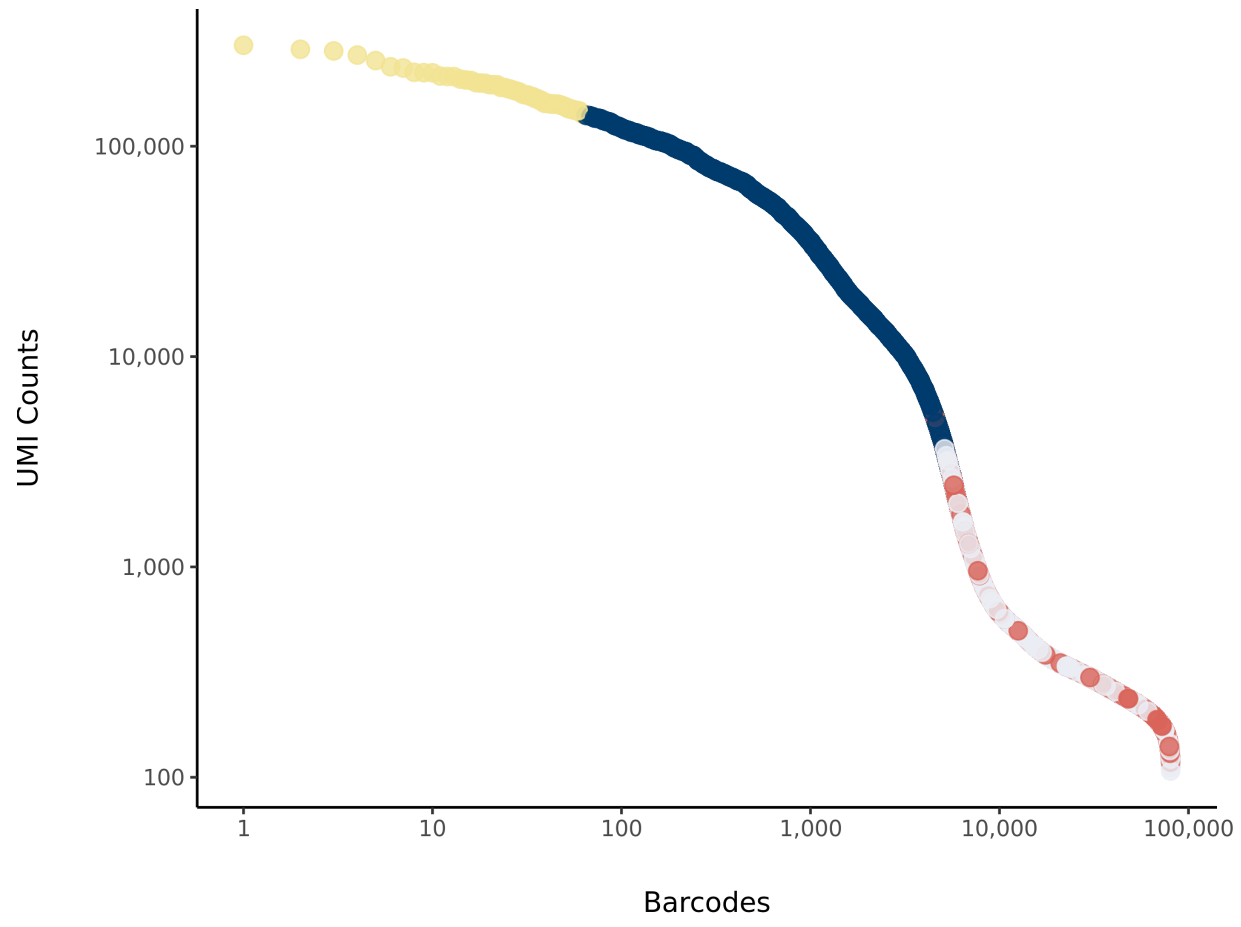

### sc_37 Summary

Processed by COPILOT

Summary Analysis

#### Parameters

| | |
|---|---|
| Iteration of Filtering | 1 |
| Mitochondrial Expression Threshold | 5 % |
| Top High Quality Cell Filtered | 1 % |
| Doublet Removed | Yes |

#### Cell Stats

| | |
|---|---|
| Estimated Number of High Quality Cell | 6,156 |
| High Quality Cell | 7.98 % |
| Total UMI Counts in High Quality Cell | 106,794,708 |
| UMI Counts in High Quality Cell | 77.91 % |
| Median UMI Counts per High Quality Cell | 5,611 |
| Median Genes per High Quality Cell | 1,984 |
| Total Genes Detected in High Quality Cell | 25,262 |
| Cell above Mitochondrial Expression Threshold | 3 % |
| Estimated Doublet Rate in High Quality Cell | 4.65 % |

#### Sequencing Stats

| | |
|---|---|
| Number of Reads Processed | 286,251,558 |
| Reads Pseudoaligned | 94.3 % |
| Reads on Whitelist | 96.52 % |
| Total UMI Counts | 137,082,512 |
| Sequencing Technology | 10xv3 |
| Species | Arabidopsis thaliana |
| Transcriptome | TAIR10 |

#### Sample Stats

| | |
|---|---|
| Sample | sc_37 |
| Name | WT Col-0 untreated |
| Source | Benfey lab |
| Genotype | WT Col-0 |
| Transgene | NA |
| Treatment | Untreated |
| Age | 6_day |
| Timepoint | 0 |
| Rep | NA |
| Target Cells | 10,000 |
| Date | 2020-02-11 |
| Seq Run | Nolan_6199 (NextSeq); Nolan_6226 (NovaSeq S4) |

#### UMI Counts Histogram

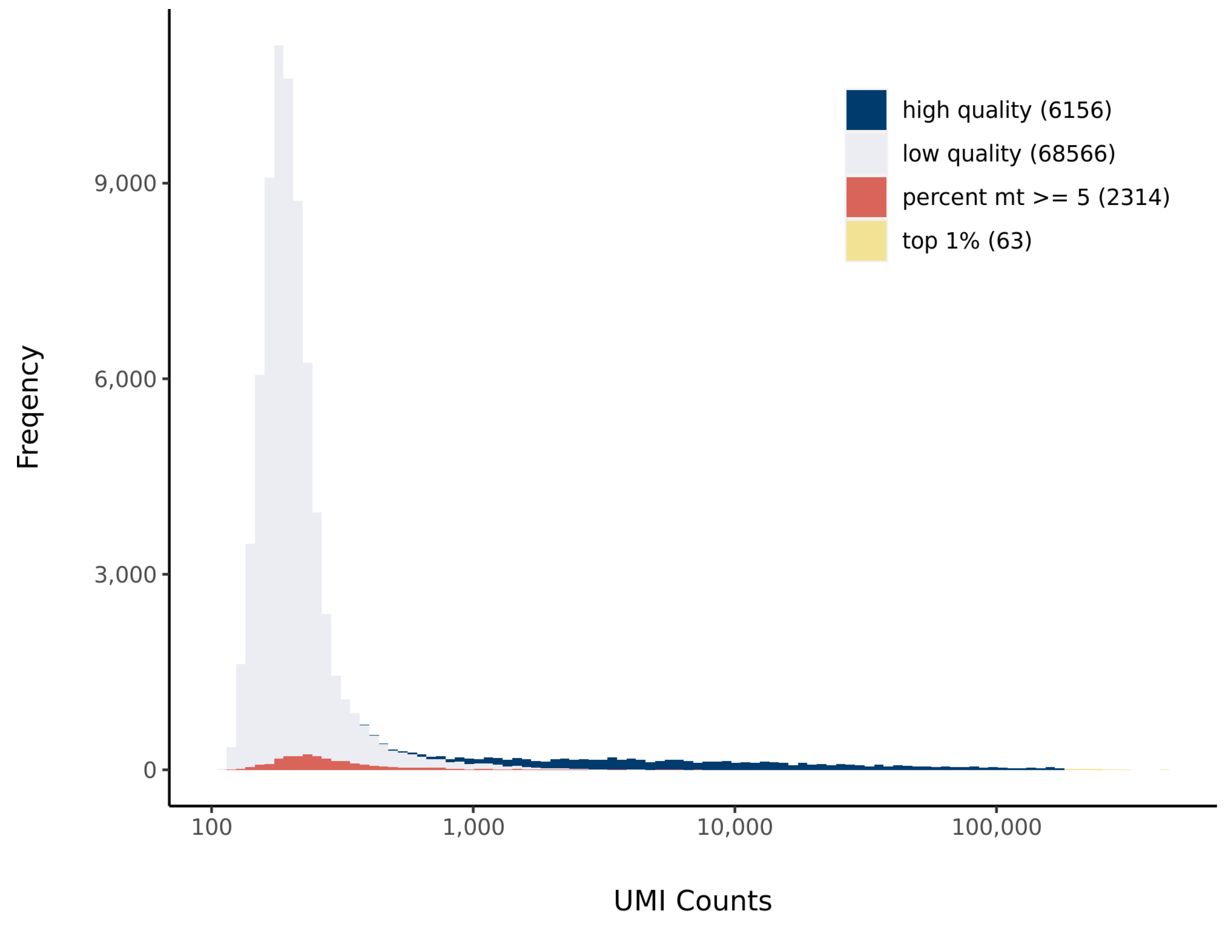


#### Number of Genes Histogram

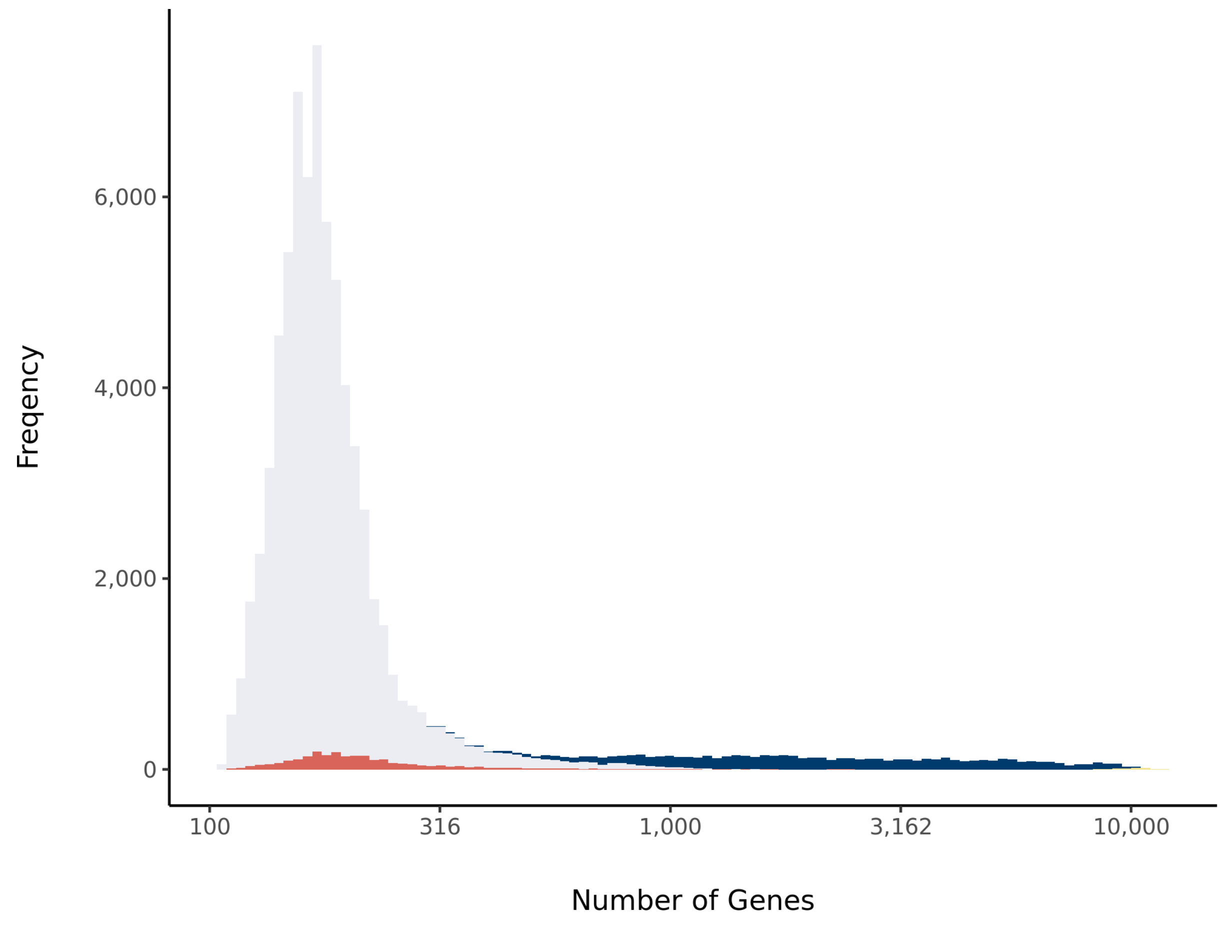


#### Barcode Rank Plot

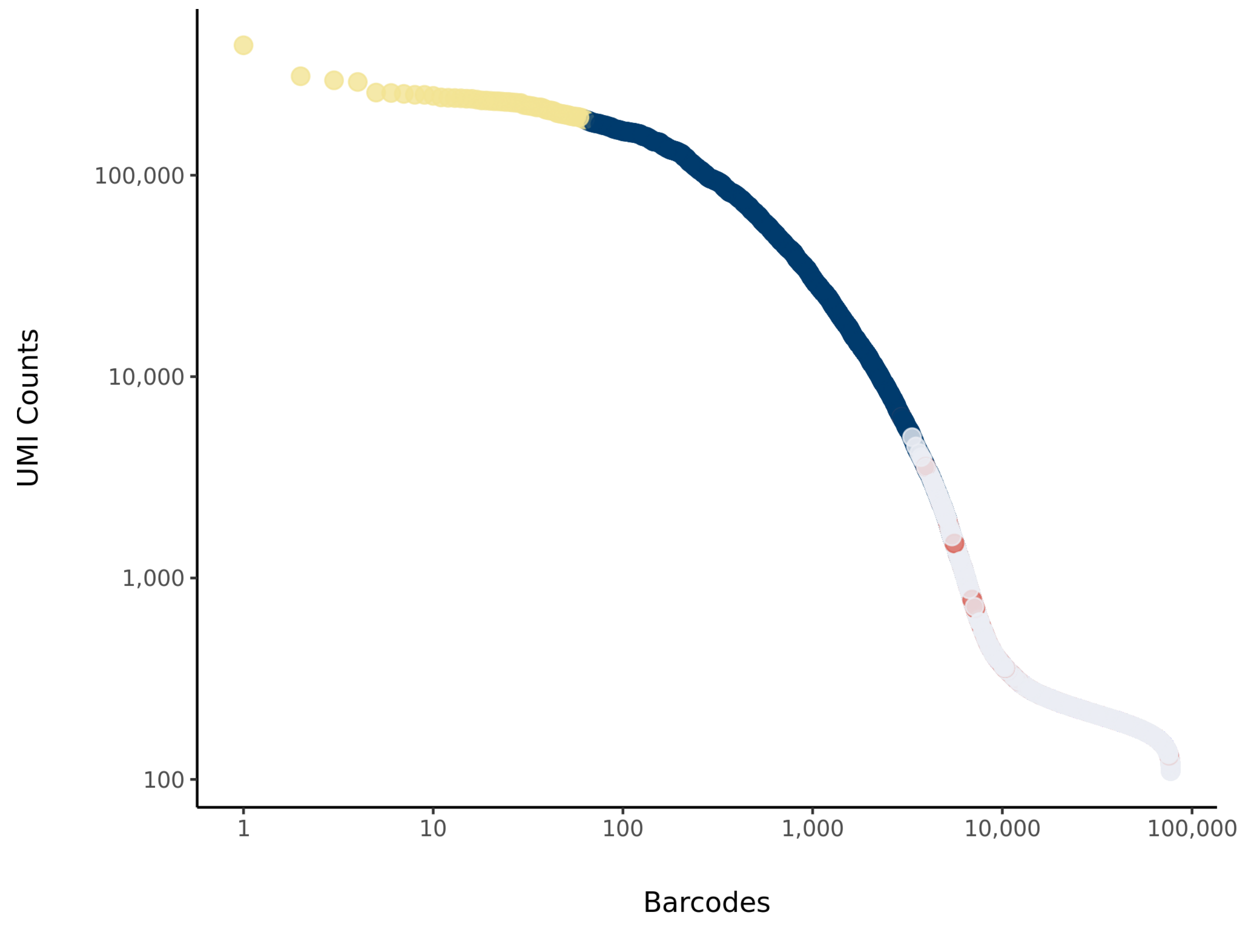

### sc_40 Summary

Processed by COPILOT

Summary Analysis

#### Parameters

| | |
|---|---|
| Iteration of Filtering | 1 |
| Mitochondrial Expression Threshold | 5 % |
| Top High Quality Cell Filtered | 1 % |
| Doublet Removed | Yes |

#### Cell Stats

| | |
|---|---|
| Estimated Number of High Quality Cell | 7,733 |
| High Quality Cell | 30.25 % |
| Total UMI Counts in High Quality Cell | 95,968,452 |
| UMI Counts in High Quality Cell | 84.61 % |
| Median UMI Counts per High Quality Cell | 4,244 |
| Median Genes per High Quality Cell | 1,681 |
| Total Genes Detected in High Quality Cell | 25,693 |
| Cell above Mitochondrial Expression Threshold | 16.34 % |
| Estimated Doublet Rate in High Quality Cell | 5.81 % |

#### Sequencing Stats

| | |
|---|---|
| Number of Reads Processed | 276,614,617 |
| Reads Pseudoaligned | 93 % |
| Reads on Whitelist | 96.28 % |
| Total UMI Counts | 113,425,205 |
| Sequencing Technology | 10xv3 |
| Species | Arabidopsis thaliana |
| Transcriptome | TAIR10 |

#### Sample Stats

| | |
|---|---|
| Sample | sc_40 |
| Name | WT Col-0 untreated |
| Source | Benfey lab |
| Genotype | WT Col-0 |
| Transgene | NA |
| Treatment | Untreated |
| Age | 6_day |
| Timepoint | 0 |
| Rep | NA |
| Target Cells | 10,000 |
| Date | 2020-02-11 |
| Seq Run | Nolan_6199 (NextSeq); Nolan_6226 (NovaSeq S4) |

#### UMI Counts Histogram

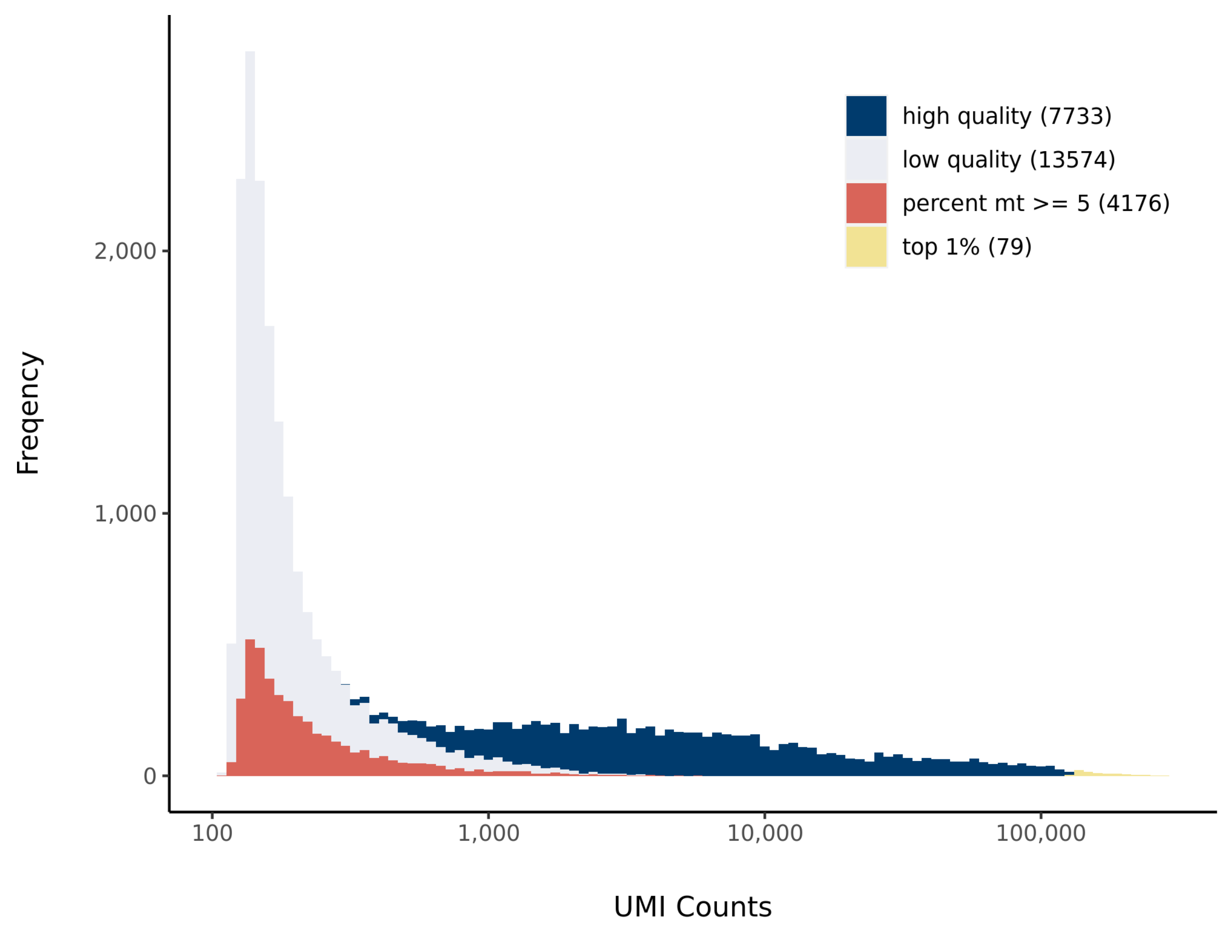


#### Number of Genes Histogram

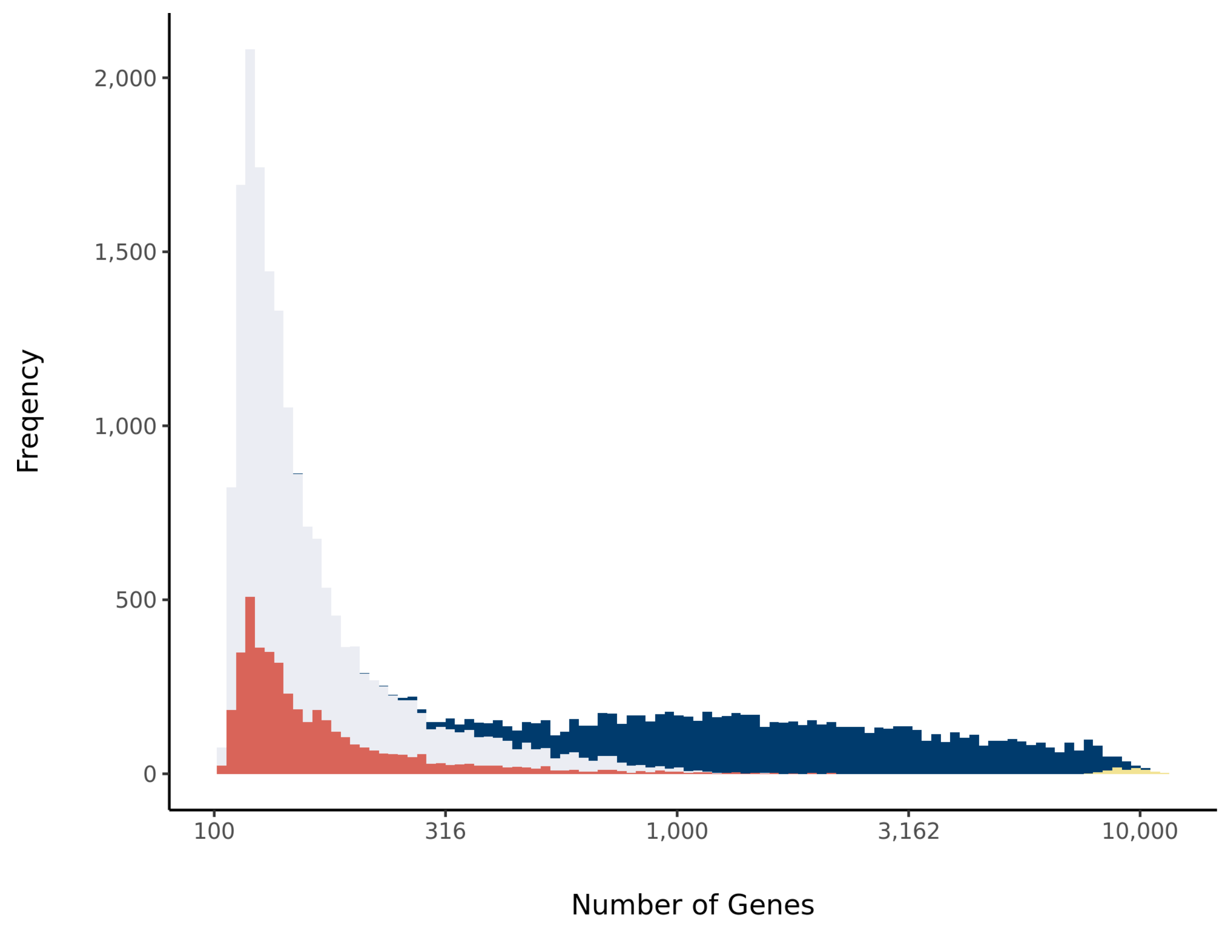


#### Barcode Rank Plot

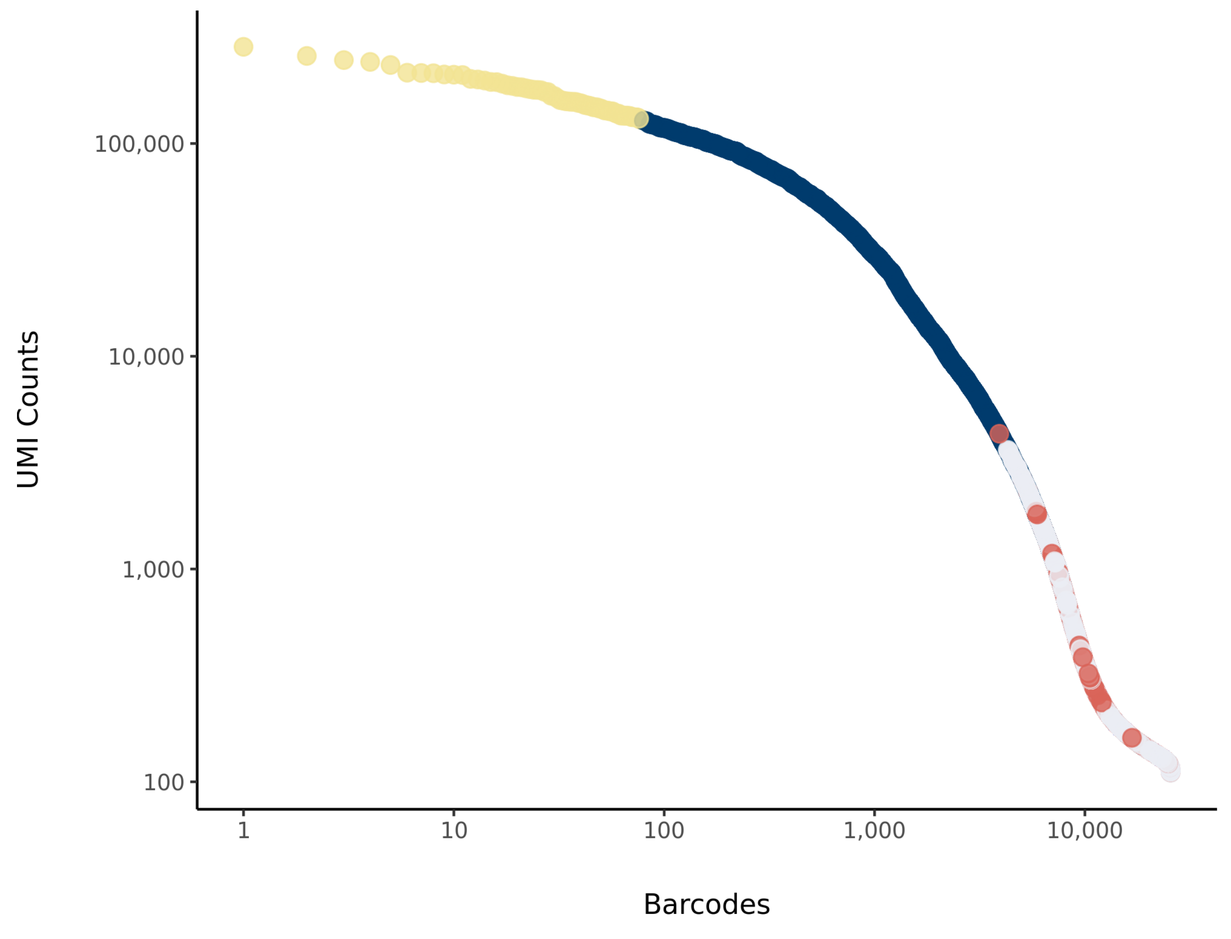

### sc_51 Summary

Processed by COPILOT

Summary Analysis

#### Parameters

| | |
|---|---|
| Iteration of Filtering | 1 |
| Mitochondrial Expression Threshold | 5 % |
| Top High Quality Cell Filtered | 1 % |
| Doublet Removed | Yes |

#### Cell Stats

| | |
|---|---|
| Estimated Number of High Quality Cell | 6,736 |
| High Quality Cell | 9.44 % |
| Total UMI Counts in High Quality Cell | 119,520,842 |
| UMI Counts in High Quality Cell | 77.74 % |
| Median UMI Counts per High Quality Cell | 8,893.5 |
| Median Genes per High Quality Cell | 2,997.5 |
| Total Genes Detected in High Quality Cell | 24,754 |
| Cell above Mitochondrial Expression Threshold | 11.29 % |
| Estimated Doublet Rate in High Quality Cell | 5.08 % |

#### Sequencing Stats

| | |
|---|---|
| Number of Reads Processed | 246,772,773 |
| Reads Pseudoaligned | 94 % |
| Reads on Whitelist | 96.29 % |
| Total UMI Counts | 153,752,550 |
| Sequencing Technology | 10xv3 |
| Species | Arabidopsis thaliana |
| Transcriptome | TAIR10 |

#### Sample Stats

| | |
|---|---|
| Sample | sc_51 |
| Name | WT Col (RS_5) |
| Source | Benfey lab |
| Genotype | WT Col-0 |
| Transgene | NA |
| Treatment | Untreated |
| Age | 5_day |
| Timepoint | NA |
| Rep | NA |
| Target Cells | 10,000 |
| Date | 2020-02-24 |
| Seq Run | Nolan_6226 |

#### UMI Counts Histogram

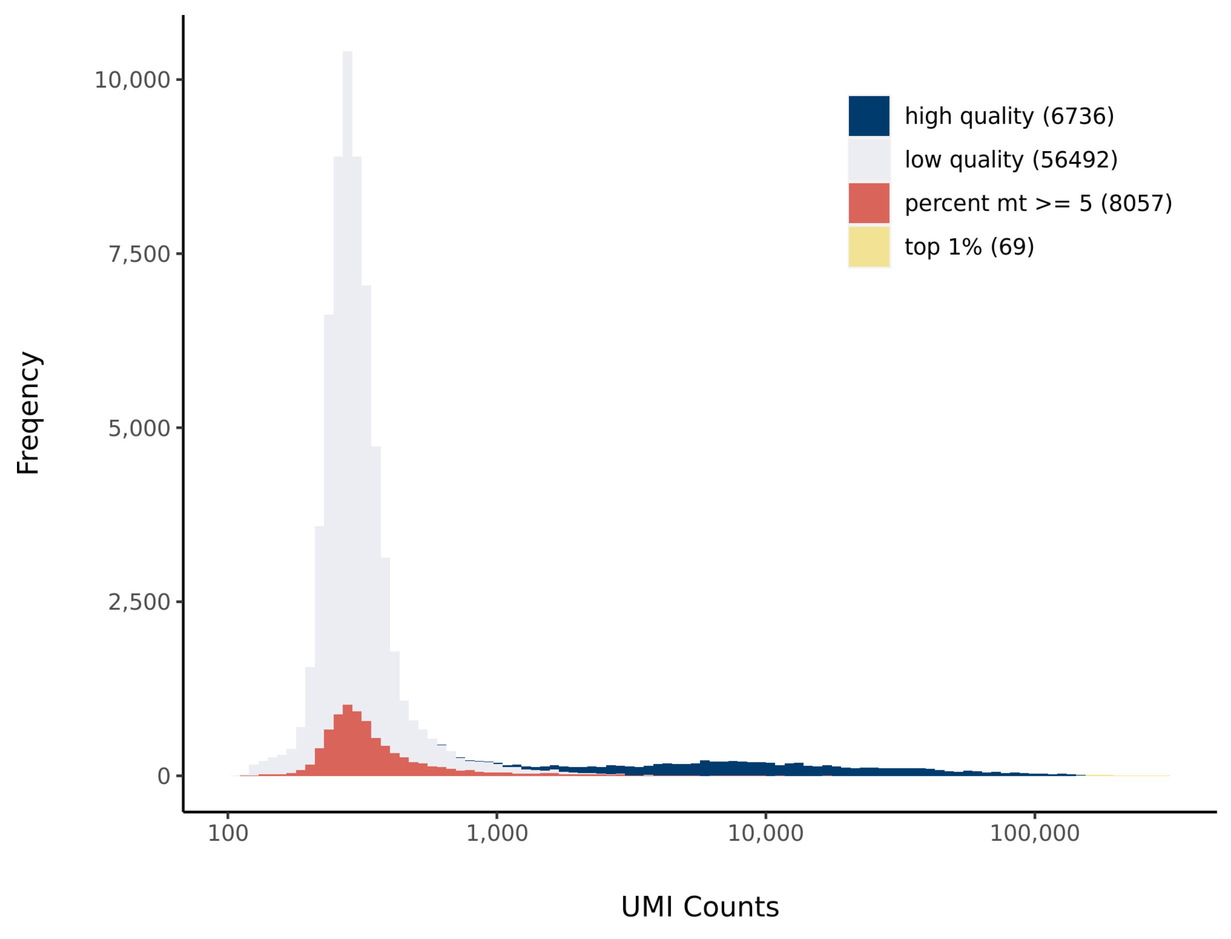


#### Number of Genes Histogram

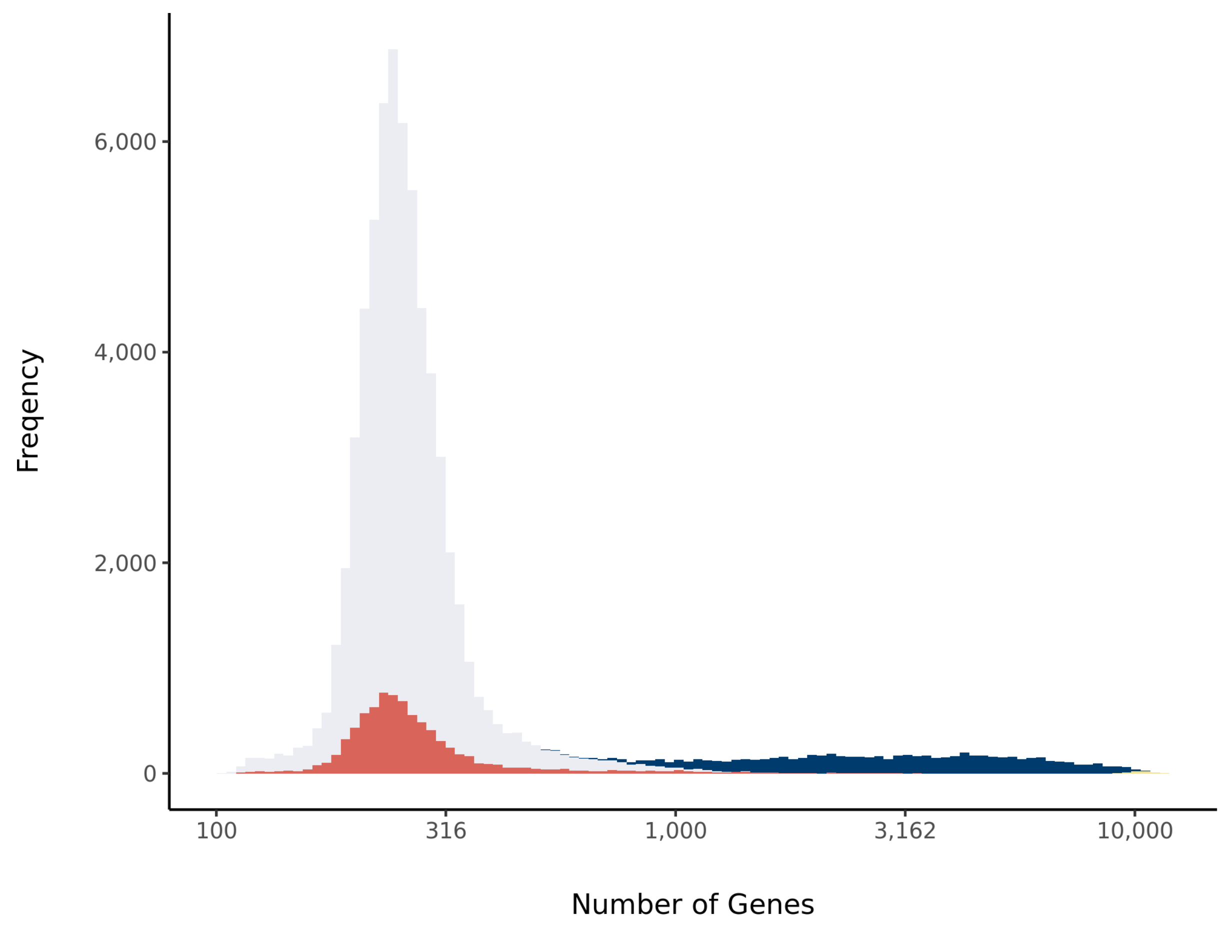


#### Barcode Rank Plot

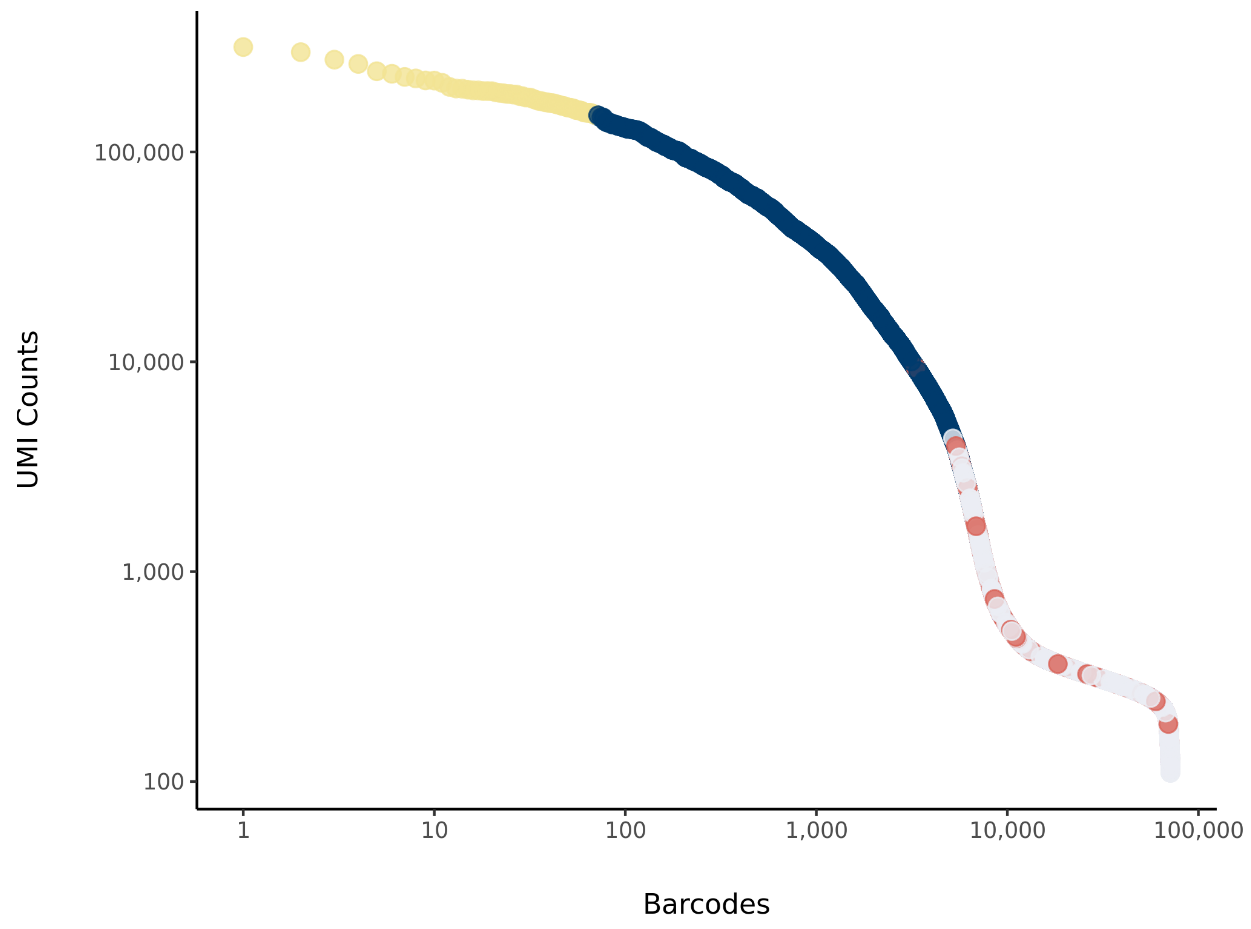

### sc_52 Summary

Processed by COPILOT

Summary Analysis

#### Parameters

| | |
|---|---|
| Iteration of Filtering | 1 |
| Mitochondrial Expression Threshold | 5 % |
| Top High Quality Cell Filtered | 1 % |
| Doublet Removed | Yes |

#### Cell Stats

| | |
|---|---|
| Estimated Number of High Quality Cell | 6,566 |
| High Quality Cell | 17.22 % |
| Total UMI Counts in High Quality Cell | 122,323,546 |
| UMI Counts in High Quality Cell | 87.97 % |
| Median UMI Counts per High Quality Cell | 9,925.5 |
| Median Genes per High Quality Cell | 3,122.5 |
| Total Genes Detected in High Quality Cell | 24,747 |
| Cell above Mitochondrial Expression Threshold | 20.25 % |
| Estimated Doublet Rate in High Quality Cell | 4.95 % |

#### Sequencing Stats

| | |
|---|---|
| Number of Reads Processed | 239,215,784 |
| Reads Pseudoaligned | 94.1 % |
| Reads on Whitelist | 96.37 % |
| Total UMI Counts | 139,046,144 |
| Sequencing Technology | 10xv3 |
| Species | Arabidopsis thaliana |
| Transcriptome | TAIR10 |

#### Sample Stats

| | |
|---|---|
| Sample | sc_52 |
| Name | shr-2 |
| Source | Benfey lab |
| Genotype | shr-2 |
| Transgene | NA |
| Treatment | Untreated |
| Age | 5_day |
| Timepoint | NA |
| Rep | NA |
| Target Cells | 10,000 |
| Date | 2020-02-24 |
| Seq Run | Nolan_6226 |

#### UMI Counts Histogram

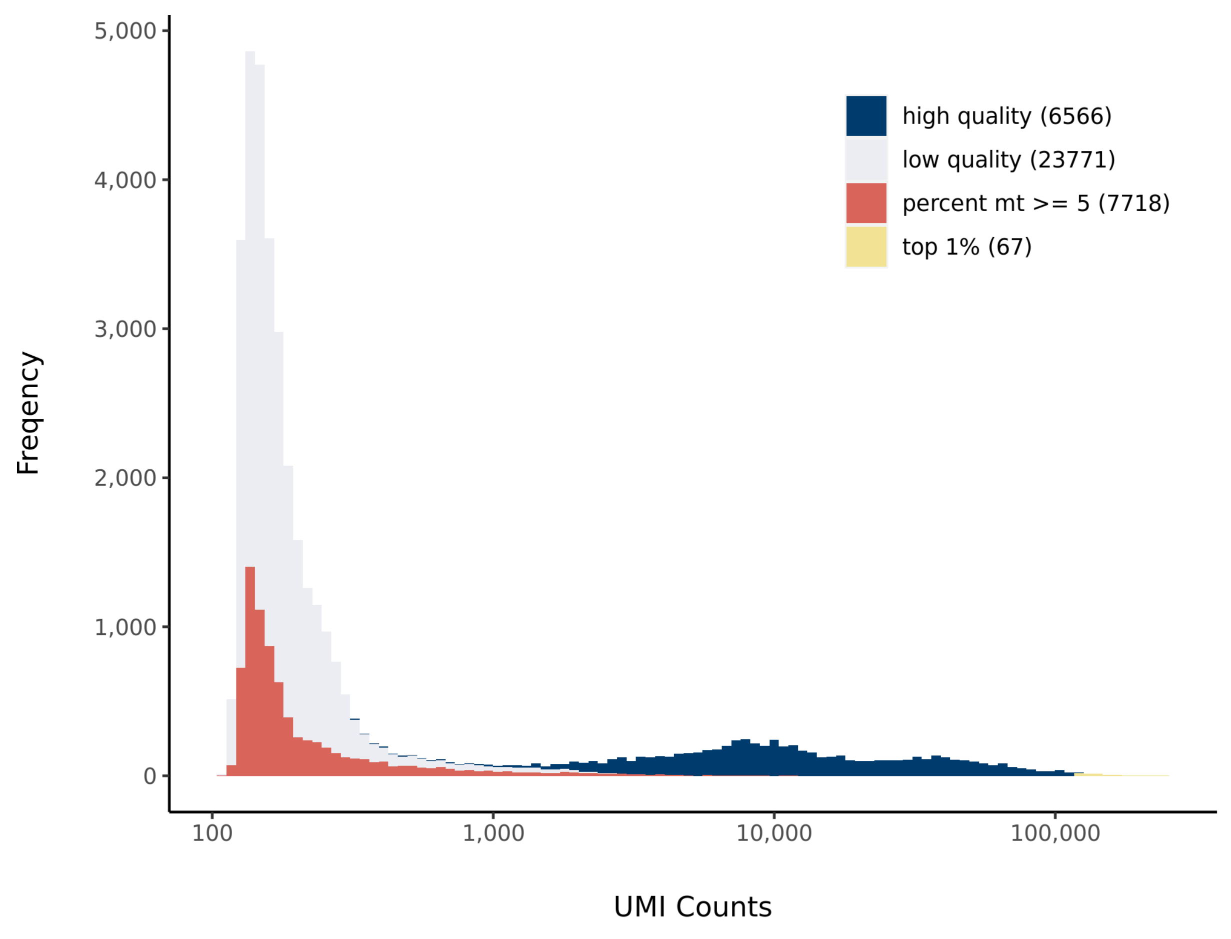


#### Number of Genes Histogram

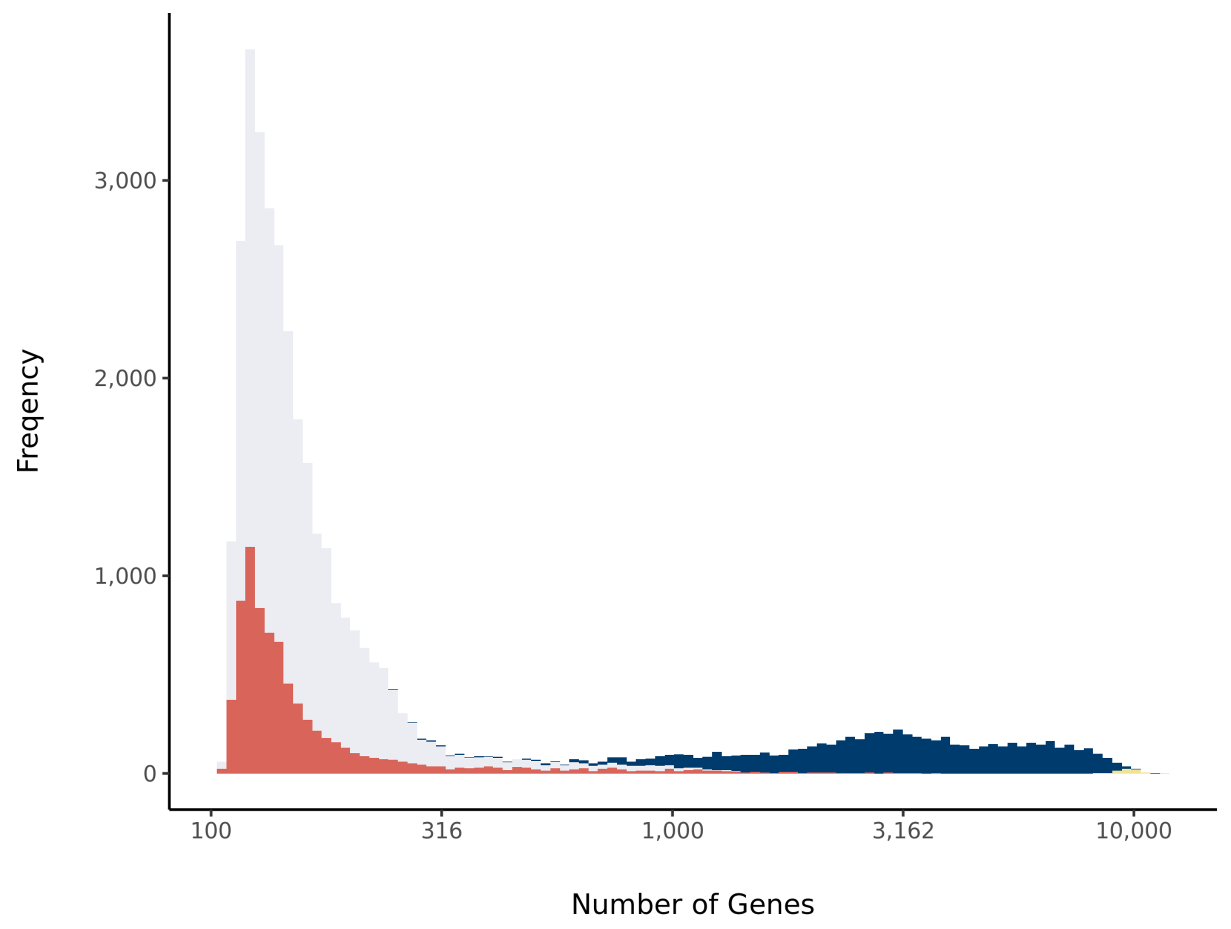


#### Barcode Rank Plot

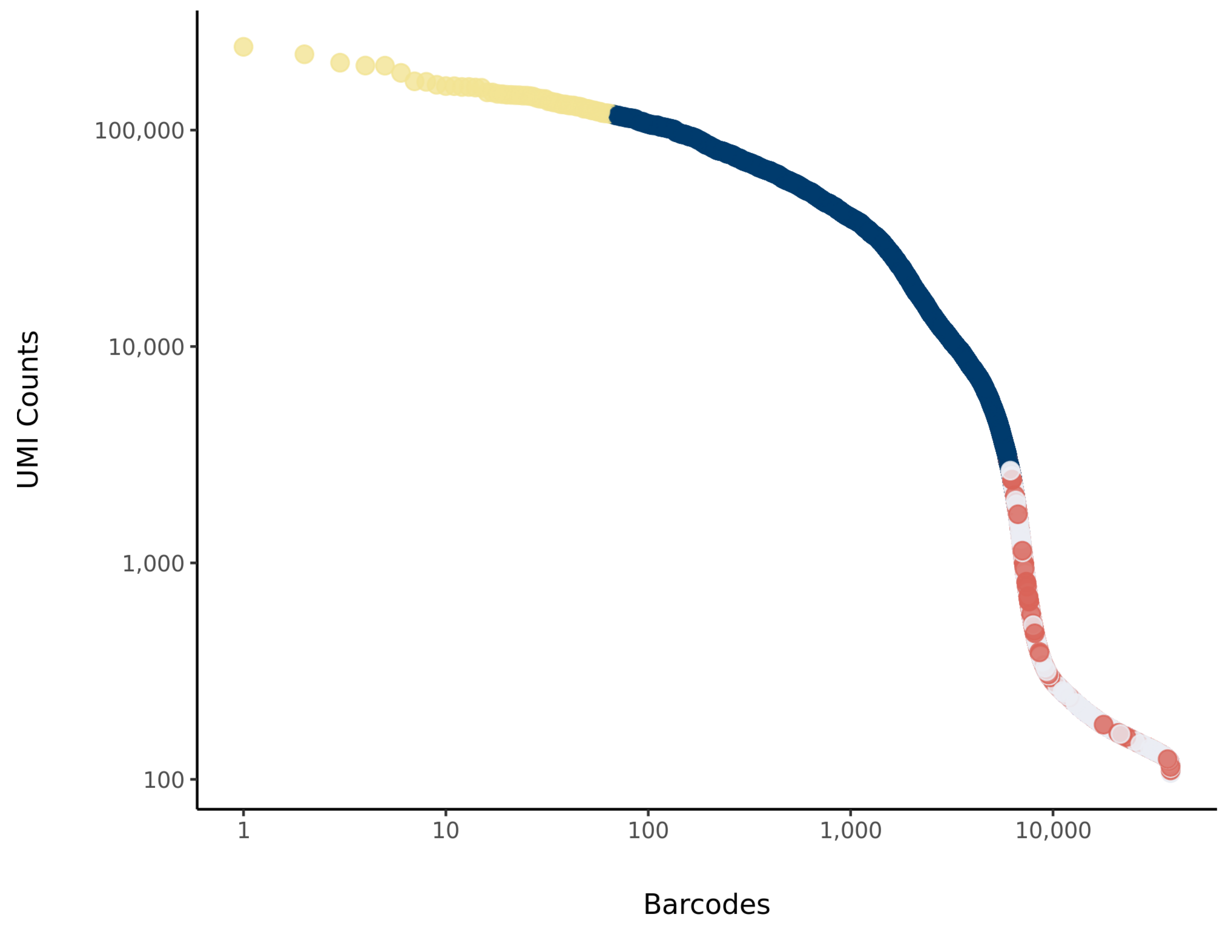

### sc_53 Summary

Processed by COPILOT

Summary Analysis

#### Parameters

| | |
|---|---|
| Iteration of Filtering | 1 |
| Mitochondrial Expression Threshold | 5 % |
| Top High Quality Cell Filtered | 1 % |
| Doublet Removed | Yes |

#### Cell Stats

| | |
|---|---|
| Estimated Number of High Quality Cell | 8,975 |
| High Quality Cell | 28.61 % |
| Total UMI Counts in High Quality Cell | 130,874,774 |
| UMI Counts in High Quality Cell | 89.12 % |
| Median UMI Counts per High Quality Cell | 7,585 |
| Median Genes per High Quality Cell | 2,784 |
| Total Genes Detected in High Quality Cell | 24,809 |
| Cell above Mitochondrial Expression Threshold | 15.34 % |
| Estimated Doublet Rate in High Quality Cell | 6.73 % |

#### Sequencing Stats

| | |
|---|---|
| Number of Reads Processed | 244,261,968 |
| Reads Pseudoaligned | 93.9 % |
| Reads on Whitelist | 96.36 % |
| Total UMI Counts | 146,853,427 |
| Sequencing Technology | 10xv3 |
| Species | Arabidopsis thaliana |
| Transcriptome | TAIR10 |

#### Sample Stats

| | |
|---|---|
| Sample | sc_53 |
| Name | shr-2 |
| Source | Benfey lab |
| Genotype | shr-2 |
| Transgene | NA |
| Treatment | Untreated |
| Age | 5_day |
| Timepoint | NA |
| Rep | NA |
| Target Cells | 10,000 |
| Date | 2020-02-24 |
| Seq Run | Nolan_6226 |

#### UMI Counts Histogram

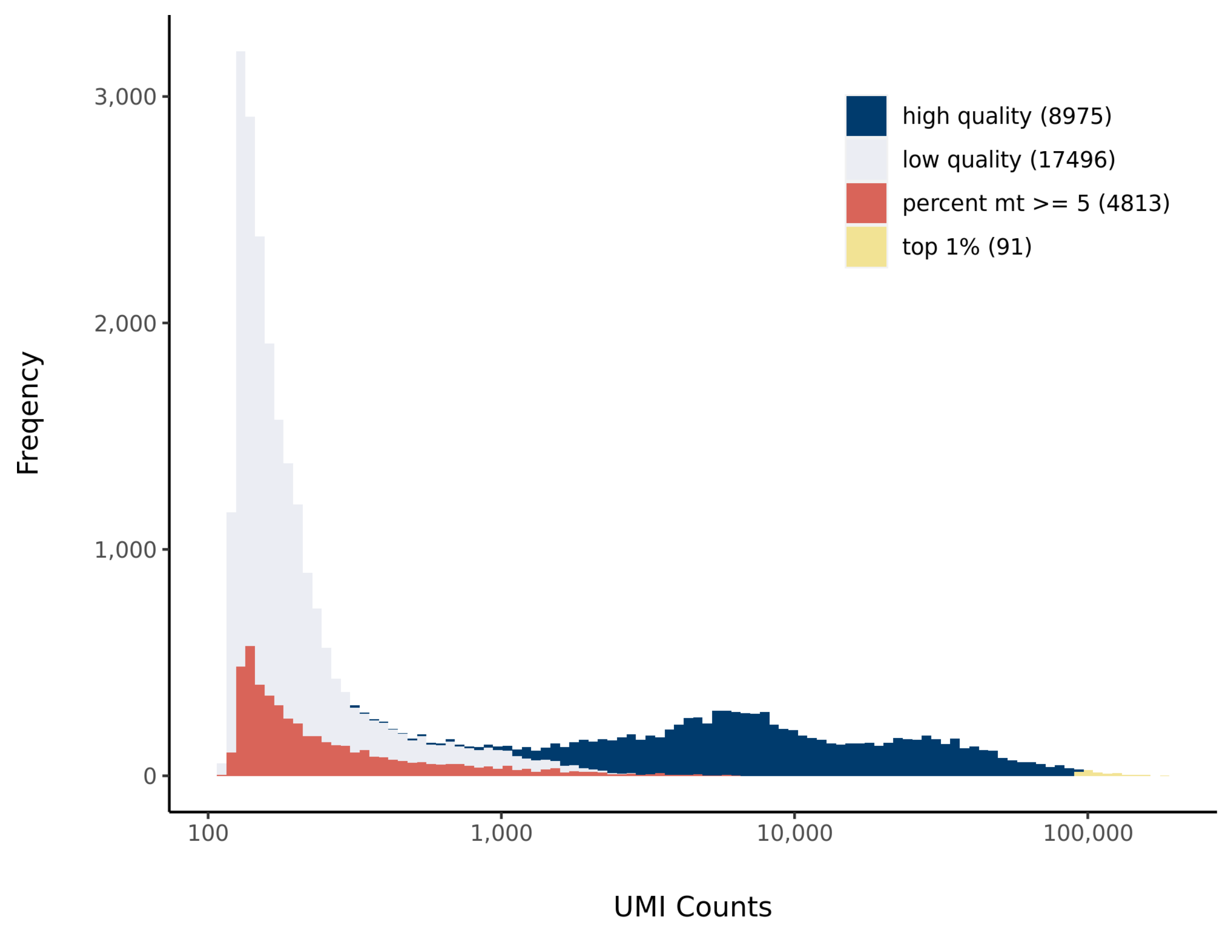


#### Number of Genes Histogram

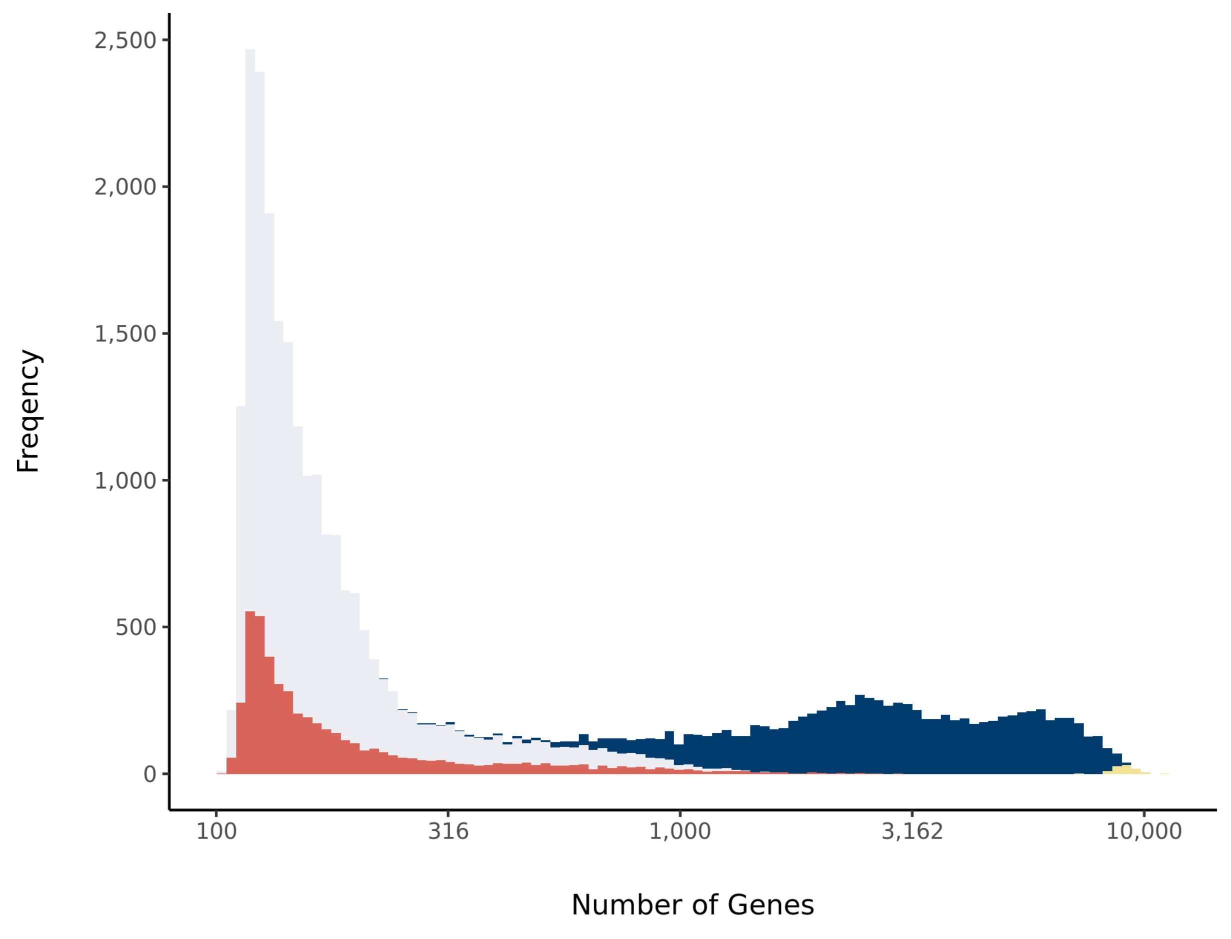


#### Barcode Rank Plot

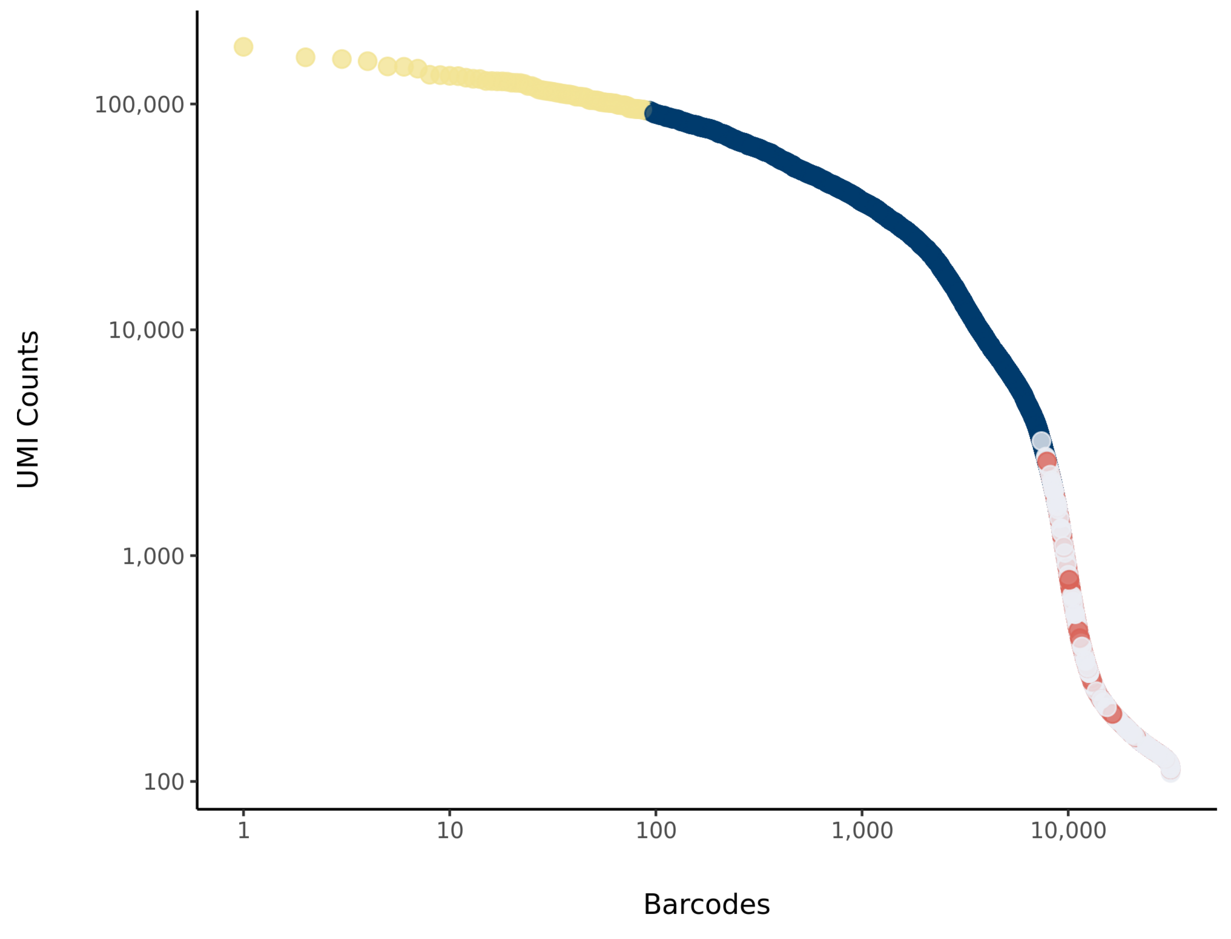

### tnw1 Summary

Processed by COPILOT

Summary | Analysis

#### Parameters

| | |
|---|---|
| Iteration of Filtering | 1 |
| Mitochondrial Expression Threshold | 5 % |
| Top High Quality Cell Filtered | 1 % |
| Doublet Removed | Yes |

#### Cell Stats

| | |
|---|---|
| Estimated Number of High Quality Cell | 5,083 |
| High Quality Cell | 6.03 % |
| Total UMI Counts in High Quality Cell | 108,012,603 |
| UMI Counts in High Quality Cell | 49.04 % |
| Median UMI Counts per High Quality Cell | 12,798 |
| Median Genes per High Quality Cell | 3,426 |
| Total Genes Detected in High Quality Cell | 24,940 |
| Cell above Mitochondrial Expression Threshold | 4.84 % |
| Estimated Doublet Rate in High Quality Cell | 3.86 % |

#### Sequencing Stats

| | |
|---|---|
| Number of Reads Processed | 487,124,902 |
| Reads Pseudoaligned | 91.5 % |
| Reads on Whitelist | 95.99 % |
| Total UMI Counts | 220,259,890 |
| Sequencing Technology | 10xv3 |
| Species | Arabidopsis thaliana |
| Transcriptome | TAIR10 |

#### Sample Stats

| | |
|---|---|
| Sample | tnw1 |
| Name | WT control1 |
| Source | Benfey lab |
| Genotype | WT Col-0 |
| Transgene | NA |
| Treatment | Untreated |
| Age | 5_day |
| Timepoint | 4 |
| Rep | 1 |
| Target Cells | 5,000 |
| Date | 2019-10-17 |
| Seq Run | Nolan_6013 |

#### UMI Counts Histogram

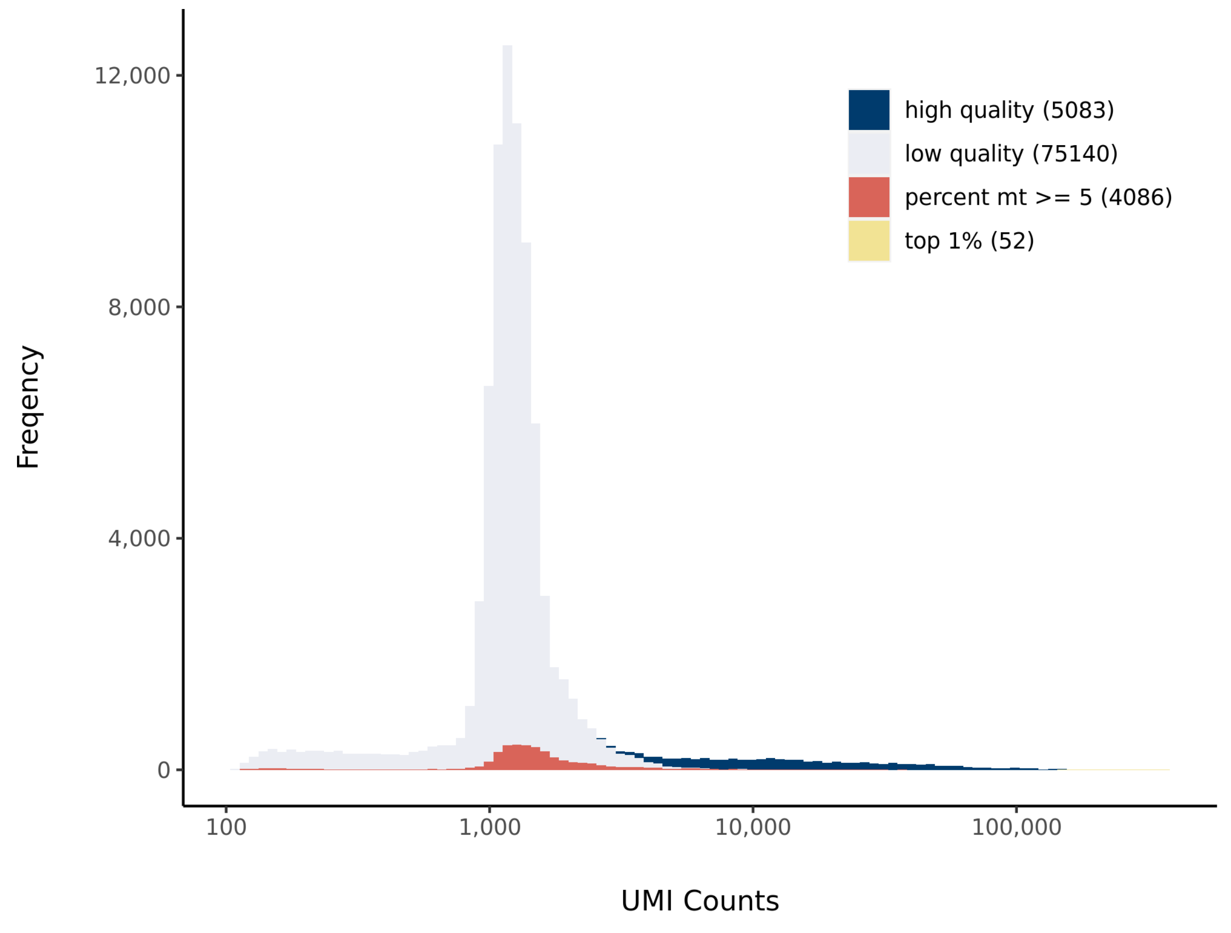


#### Number of Genes Histogram

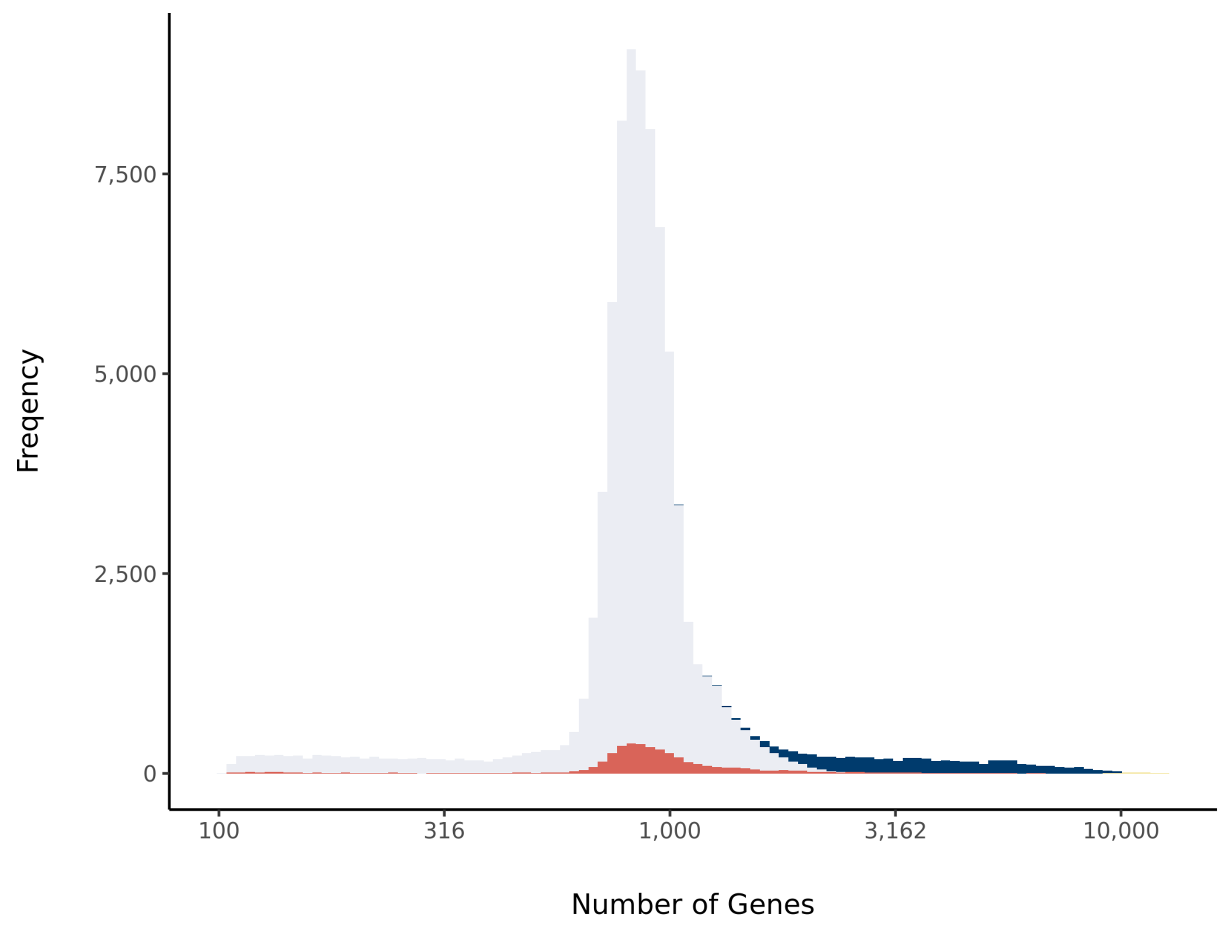


#### Barcode Rank Plot

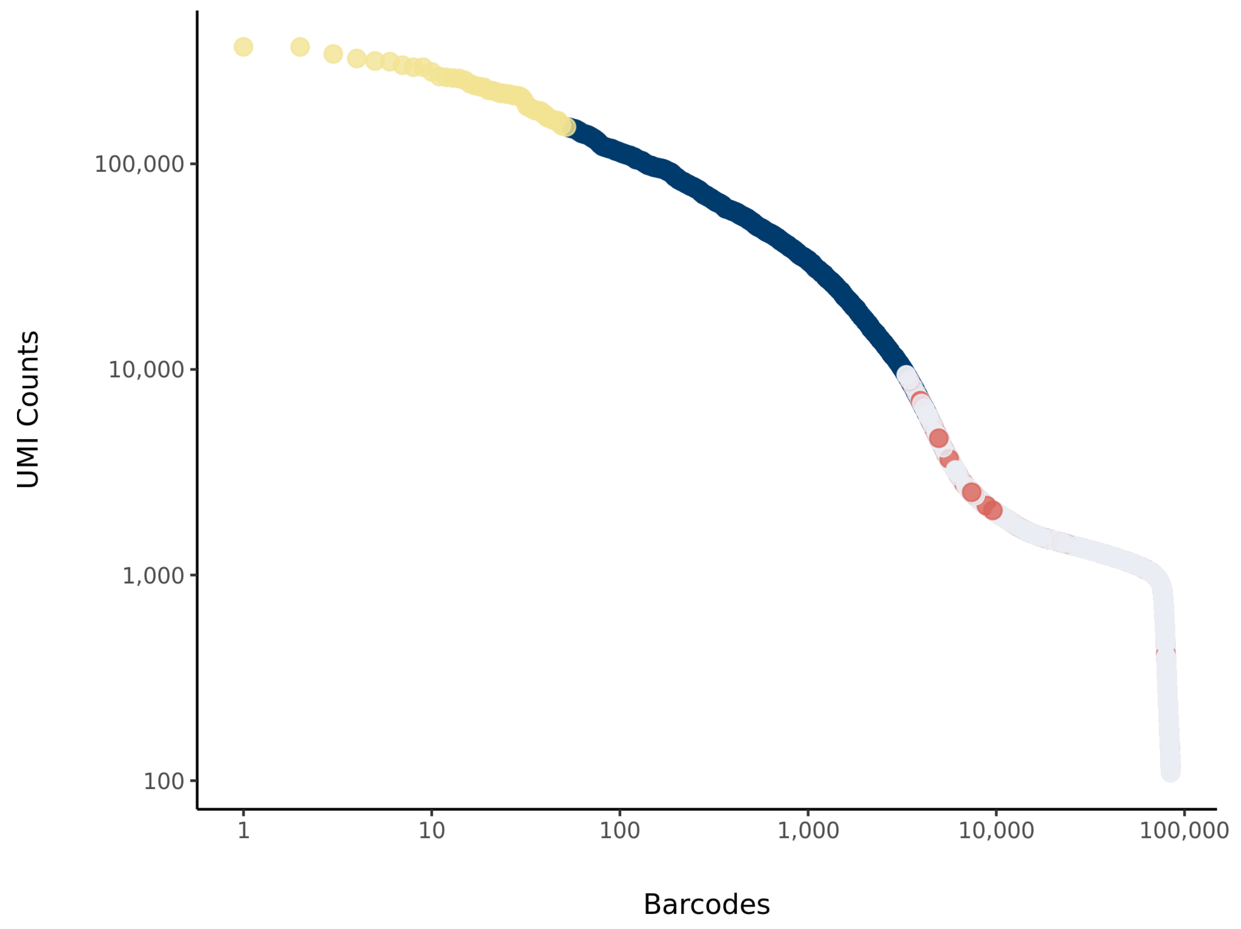

### tnw2 Summary

Processed by COPILOT

Summary Analysis

#### Parameters

| | |
|---|---|
| Iteration of Filtering | 1 |
| Mitochondrial Expression Threshold | 5 % |
| Top High Quality Cell Filtered | 1 % |
| Doublet Removed | Yes |

#### Cell Stats

| | |
|---|---|
| Estimated Number of High Quality Cell | 4,065 |
| High Quality Cell | 5.23 % |
| Total UMI Counts in High Quality Cell | 114,947,259 |
| UMI Counts in High Quality Cell | 59.81 % |
| Median UMI Counts per High Quality Cell | 15,794 |
| Median Genes per High Quality Cell | 3,557 |
| Total Genes Detected in High Quality Cell | 25,145 |
| Cell above Mitochondrial Expression Threshold | 14.57 % |
| Estimated Doublet Rate in High Quality Cell | 3.12 % |

#### Sequencing Stats

| | |
|---|---|
| Number of Reads Processed | 481,231,001 |
| Reads Pseudoaligned | 93.1 % |
| Reads on Whitelist | 96.28 % |
| Total UMI Counts | 192,190,179 |
| Sequencing Technology | 10xv3 |
| Species | Arabidopsis thaliana |
| Transcriptome | TAIR10 |

#### Sample Stats

| | |
|---|---|
| Sample | tnw2 |
| Name | WT control2 |
| Source | Benfey lab |
| Genotype | WT Col-0 |
| Transgene | NA |
| Treatment | Untreated |
| Age | 5_day |
| Timepoint | 4 |
| Rep | 2 |
| Target Cells | 5,000 |
| Date | 2019-10-17 |
| Seq Run | Nolan_6013 |

#### UMI Counts Histogram

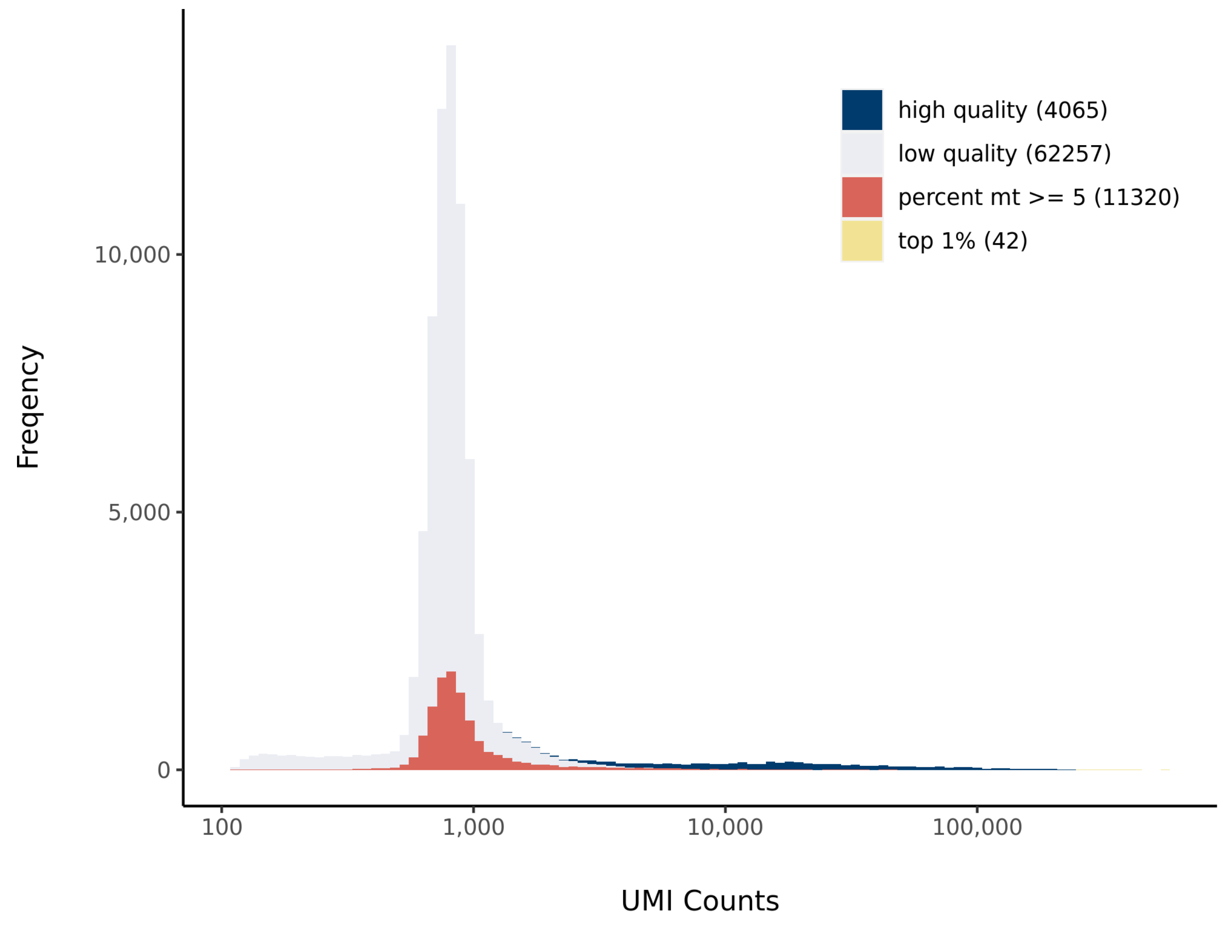


#### Number of Genes Histogram

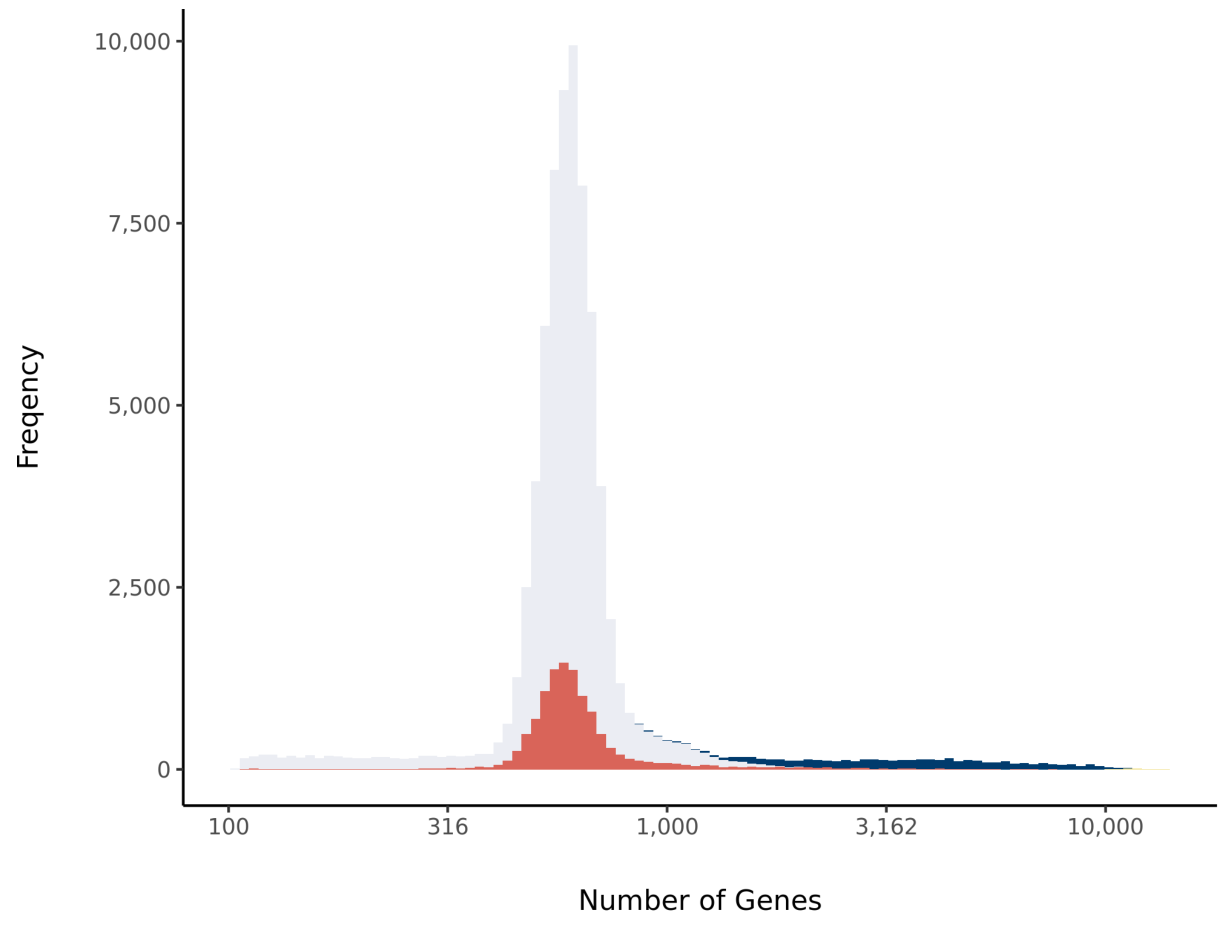


#### Barcode Rank Plot

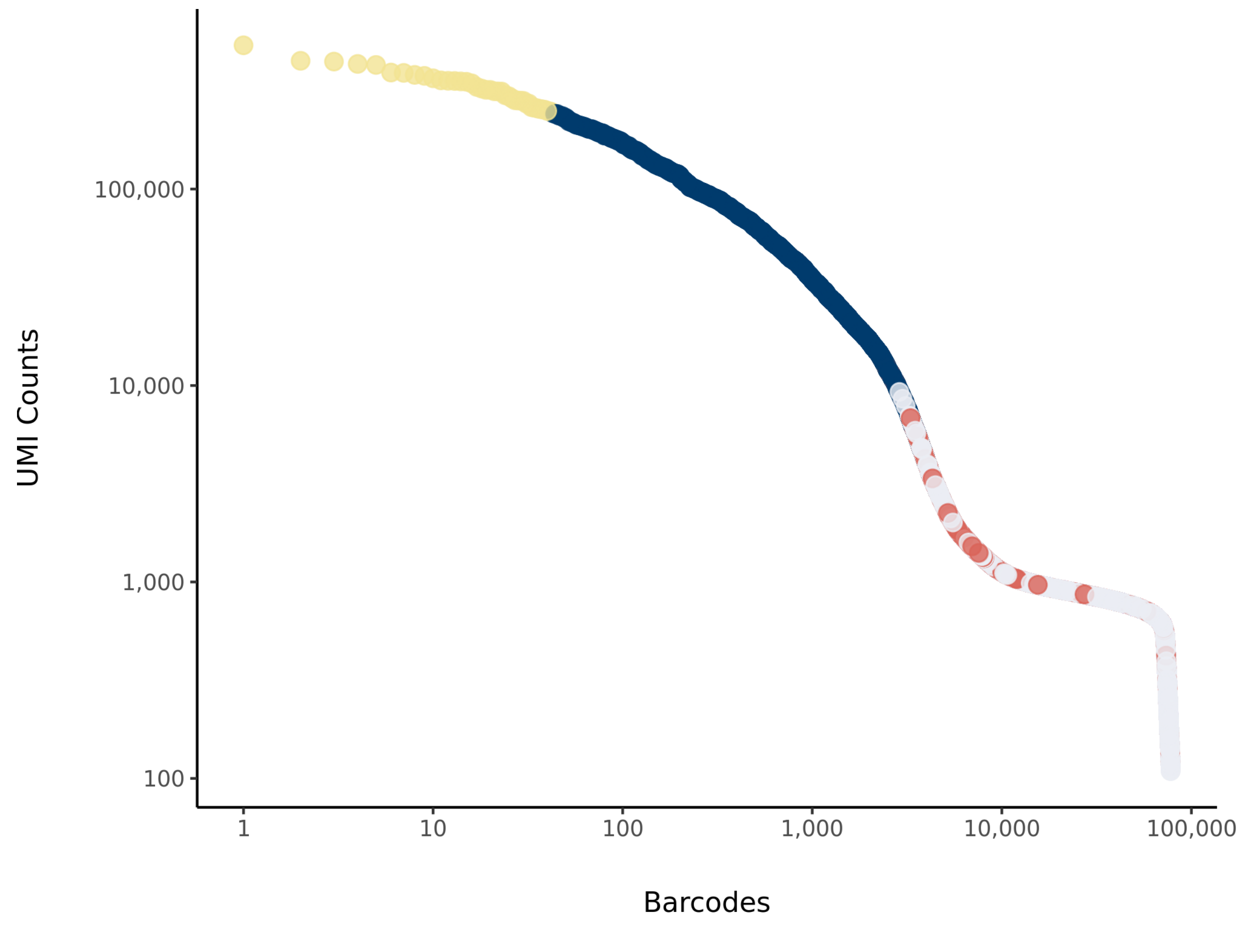
